# Comprehensive profiling of genomic and transcriptomic differences between risk groups of lung adenocarcinoma and lung squamous cell carcinoma

**DOI:** 10.1101/2020.12.31.424952

**Authors:** Talip Zengin, Tuğba Önal-Süzek

## Abstract

Lung cancer is the second frequently diagnosed cancer type and responsible for the highest number of cancer deaths worldwide. Lung adenocarcinoma and lung squamous cell carcinoma are subtypes of non-small cell lung cancer which has the highest frequency of lung cancer cases. We aimed to analyze genomic and transcriptomic variations including simple nucleotide variations (SNVs), copy number variations (CNVs) and differential expressed genes (DEGs) in order to find key genes and pathways for diagnostic and prognostic prediction for lung adenocarcinoma and lung squamous cell carcinoma. We performed univariate cox model and then lasso regularized cox model with leave-one-out cross-validation using TCGA gene expression data in tumor samples. We generated a 35-gene signature and a 33-gene signature for prognostic risk prediction based on the overall survival time of the patients with LUAD and LUSC, respectively. When we clustered patients into high-risk and low-risk groups, the survival analysis showed highly significant results with high prediction power for both training and test datasets. Then we characterized the differences including significant SNVs, CNVs, DEGs, active subnetworks, and the pathways. We described the results for the risk groups and cancer subtypes separately to identify specific genomic alterations between both high-risk groups and cancer subtypes. Both LUAD and LUSC high-risk groups have more down-regulated immune pathways and upregulated metabolic pathways. On the other hand, low-risk groups have both upregulated and downregulated genes on cancer-related pathways. Both LUAD and LUSC have important gene alterations such as CDKN2A and CDKN2B deletions with different frequencies. SOX2 amplification occurs in LUSC and PSMD4 amplification in LUAD. EGFR and KRAS mutations are mutually exclusive in LUAD samples. EGFR, MGA, SMARCA4, ATM, RBM10, and KDM5C genes are mutated only in LUAD but not in LUSC. CDKN2A, PTEN, and HRAS genes are mutated only in LUSC samples. Low-risk groups of both LUAD and LUSC, tend to have a higher number of SNVs, CNVs, and DEGs. The signature genes and altered genes have the potential to be used as diagnostic and prognostic biomarkers for personalized oncology.

## Introduction

Lung cancer is the second frequently diagnosed cancer type and the leading cause of cancer-related mortality worldwifde [1]. Lung cancer treatments used in the clinic are surgery, radiotherapy, chemotherapy, targeted therapy and emerging immunotherapy. The clinical treatment decisions are made based on TNM stage, histology, genetic alterations of a few driver oncogenes for targeted therapies, and patient’s condition [2]. However, most of the patients are diagnosed at an advanced and metastatic stage, with high mortality and poor benefit from therapies [3]. Although the introduction of targeted therapeutics and immunotherapeutics including immune-checkpoint inhibitors for patients with advanced-stage, these options are beneficial only for limited subsets of patients and also these patients can develop resistance [4]. Therefore, the majority of patients with advanced-stage lung cancer die within 5 years of diagnosis [5].

Histologically there are 4 major types of lung cancer, including small cell carcinoma (SCLC), and adenocarcinoma, squamous cell carcinoma, large cell carcinoma as grouped non-small cell carcinoma (NSCLC). Lung adenocarcinoma (LUAD) and lung squamous cell carcinoma (LUSC) account for 50% and 23% of all lung cancers, respectively [6].

Lung cancer is both histologically and molecularly heterogeneous disease and characterizing the genomics and transcriptomics of its nature is very important for effective therapies. Lung cancer therapy has changed from the use of cytotoxic therapeutics to medicating patients with targeting therapeutics according to their genomic and transcriptomic alterations and the status of programmed death ligand-1 (PD-L1) which is target of immune checkpoint blockers [7]. Lung cancer has many subtypes with distinct genetic characteristics, resulting in intra-tumoral heterogeneity [8]. Kirsten rat sarcoma (KRAS) and epidermal growth factor receptor (EGFR) genes play a major role in tumorigenesis and are strong candidates for targeted therapy. KRAS and EGFR mutations are usually mutually exclusive, but if they are co-occur, KRAS mutations can cause resistance to EGFR inhibitors [9].

The most commonly mutated genes in LUAD are KRAS and EGFR oncogenes, and TP53, KEAP1, STK11 and NF1 the tumor suppressor genes. The frequency of EGFR-activating mutations varies greatly by region and ethnicity [7]. KEAP1 inactivation in the presence of KRAS mutations enhances sensitivity to glutaminase inhibitors, providing a potential therapeutic strategy for patients carrying these mutations [10].

Commonly mutated gene in LUSC is TP53, which is present in more than 90% of tumors, and CDKN2A tumor suppressors [7]. The recurrent mutations in NFE2L2, KEAP1, BAI3, FBXW7, GRM8, MUC16, RUNX1T1, STK11 and ERBB4 have been reported in LUSC [11,12]. Somatic copy number variations also have been identified in LUSC, including deletion of CDKN2A and amplification of SOX2, PDGFRA, FGFR1 and WHSC1L1 genes 9,10. DDR2 mutations and FGFR1 amplification have been nominated as therapeutic targets [13–15].

The Cancer Genome Atlas (TCGA) database serves different types of data such as transcriptome profiling, simple nucleotide variation, copy number variation, DNA methylation, clinical and biospecimen data of 84392 cancer patients with 68 primary sites [16]. The Cancer Genome Atlas Research Network reported molecular profiling of 230 lung adenocarcinoma samples using mRNA, microRNA and DNA sequencing integrated with copy number, methylation and proteomic analyses. They identified 18 significantly mutated genes, including TP53, KRAS which is mutually exclusive with EGFR, BRAF, PIK3CA, MET, STK11, KEAP1, NF1, RB1, CDKN2A, GTPase gene RIT1 including activating mutations and MGA including loss-of-function mutations. DNA and mRNA sequence from the same tumor highlighted splicing alterations including exon 14 skipping in MET mRNA in 4% of cases. They also showed DNA hyper-methylation of several key genes: CDKN2A, GATA2, GATA4, GATA5, HIC1, HOXA9, HOXD13, RASSF1, SFRP1, SOX17, WIF1 and MYC over-expression was significantly associated with the hyper-methylation phenotype as well [17].

The Cancer Genome Atlas Research Network also profiled 178 lung squamous cell carcinomas and detected mutations in 11 genes, including mutations in TP53 (81%), CDKN2A, PTEN, PIK3CA, KEAP1, MLL2, HLA-A, NFE2L2, RB1, NOTCH1 including truncating mutations and loss-of-function mutations in the HLA-A class I major histocompatibility gene. They identified altered pathways such as NFE2L2 and KEAP1 in 34%, squamous differentiation genes in 44%, PI3K pathway genes in 47%, and CDKN2A and RB1 in 72% of tumors. CNV analysis revealed the amplification of NFE2L2, MYC, CDK6, MDM2, BCL2L1 and EYS, and deletions of FOXP1, PTEN and NF1 genes with previously identified CNV genes, SOX2, PDGFRA, KIT, EGFR, FGFR1, WHSC1L1, CCND1 and CDKN2A. They identified overexpression and amplification of SOX2 and TP63, loss-of-function mutations in NOTCH1, NOTCH2 and ASCL4 and focal deletions in FOXP1 which have known roles in squamous cell differentiation. CDKN2A is down-regulated in over 70% of samples through epigenetic silencing by methylation (21%), inactivating mutation (18%), exon 1β skipping (4%), or homozygous deletion (29%) [18].

Recently, many studies have been published on gene expression signatures predicting survival risk of patients with lung adenocarcinoma. These recent studies have been mostly using TCGA data but their methods generated different gene signatures. Seven-gene expression signature including ASPM, KIF15, NCAPG, FGFR1OP, RAD51AP1, DLGAP5 and ADAM10 genes, was obtained for early stage cases from seven published lung adenocarcinoma cohorts and the signature showed high hazard rations in cox regression analysis [19]. Shukla et al. developed TCGA RNAseq data based prognostic signature including four protein-coding genes RHOV, CD109, FRRS1, and the lncRNA gene LINC00941, which showed high hazard ratios for stage I, EGFR wild-type, and EGFR mutant groups [20]. A prognostic signature which was independent of other clinical factors, was developed and validated based on the TCGA data. Patients were grouped into risk groups using signature genes, and patients with high-risk scores tended to have poor survival rate at 1-, 3- and 5-year follow-up. The developed eight-gene signature including TTK, HMMR, ASPM, CDCA8, KIF2C, CCNA2, CCNB2, and MKI67 were highly expressed in A549 and PC-9 cells [21].

Twelve-gene signature (RPL22, VEGFA, G0S2, NES, TNFRSF25, DKFZP586P0123, COL8A2, ZNF3, RIPK5, RNFT2, ARHGEF12 and PTPN20A/B) was established by using published microarray dataset from 129 patients and the signature was independently prognostic for lung squamous carcinoma but not for lung adenocarcinoma [22]. A four-gene clustering model in 14-Genes (DPPA, TTTY16, TRIM58, HKDC1, ZNF589, ALDH7A1, LINC01426, IL19, LOC101928358, TMEM92, HRASLS, JPH1, LOC100288778, GCGR) was established and these genes plays role in positive regulation of ERK1 and ERK2 cascade, angiogenesis, platelet degranulation, cell–matrix adhesion, extracellular matrix organization and macrophage activation [23].

Lu et.al. identified differentially expressed genes between lung adenocarcinoma and lung squamous cell carcinoma by using microarray data from the Gene Expression Omnibus database. They identified 95 upregulated and 241 downregulated DEGs in lung adenocarcinoma samples, and 204 upregulated and 285 downregulated DEGs in lung squamous cell carcinoma samples, compared to the normal lung tissue samples. The genes play role in cell cycle, DNA replication and mismatch repair. Top five genes from global network, HSP90AA1, BCL2, CDK2, KIT and HDAC2 have differential expression profiles between lung adenocarcinoma and lung squamous cell carcinoma [24]. Recently, Wu et.al. identified diagnostic and prognostic genes for lung adenocarcinoma and squamous cell carcinoma by using weighted gene expression profiles. The five-gene diagnostic signature including KRT5, MUC1, TREM1, C3 and TMPRSS2 and the five-gene prognostic signature including ADH1C, AZGP1, CLU, CDK1 and PEG10 obtained a log-rank P-value of 0.03 and a C-index of 0.622 on the test set [25].

Considerable number of genetic and transcriptomic alterations have been identified in mostly LUAD and poorly in LUSC. Although many gene expression signatures have been identified in LUAD recently, there is less work on LUSC expression signatures. Therebeside, the molecular differences between risk groups of LUAD and LUSC have not yet been systematically described. In this study, we aimed to identify the genomic and transcriptomic differences between risk groups of lung adenocarcinoma and lung squamous cell carcinoma. We performed univariate Cox model and then Lasso Regularized Cox Model with Leave-One-Out Cross-Validation (LOOCV) by using TCGA gene expression data in tumor samples, and identified best gene signatures to cluster patients into low-risk and high-risk groups. We generated 35-gene signature and 33-gene signature for prognostic risk prediction based on the overall survival time of the patients with LUAD and LUSC. When we clustered patients into high-risk and low-risk groups, the survival analysis showed highly significant results for both training and test datasets. Then we characterized the differences including significant SNVs, CNVs, DEGs and active subnetwork DEGs between risk groups in LUAD and LUSC.

## Materials and Methods

### Data

Simple Nucleotide Variation (SNV), Transcriptome Profiling, Copy Number Variation (CNV) and Clinical data of patients who have all of these data types in LUAD and LUSC projects, was downloaded separately using *TCGAbiolinks* R package [26]. Using the same package and the reference of hg38; Simple Nucleotide Variations (SNVs) and Copy Number Variations (CNVs); and transcriptomic variations were processed to identify the genomic alterations of the LUAD and LUSC patients (Table 1).

**Table 1.** Summary of clinical variables of train and test group of patients with LUAD and LUSC analyzed in the study

| Category | LUAD |  | LUSC |  |
| --- | --- | --- | --- | --- |
|  | Train Group<br>(n: 436) | Test Group<br>(n: 56) | Train Group<br>(n: 431) | Test Group<br>(n: 47) |
| Age at diagnosis (median; range) | 66; 33-88 | 66.5; 42-86 | 68; 39-90 | 69; 45-85 |
| Gender |  |  |  |  |
| Female | 232 | 33 | 112 | 14 |
| Male | 204 | 23 | 319 | 33 |
| Tumor stage |  |  |  |  |
| I | 241 | 28 | 211 | 25 |
| II | 106 | 13 | 138 | 16 |
| III | 68 | 13 | 76 | 5 |
| IV | 23 | 2 | 6 | 1 |
| Vital status |  |  |  |  |
| Alive | 284 | 30 | 275 | 18 |
| Dead | 152 | 26 | 156 | 29 |
| Smoked years (median; range) | 33; 2-61 | 31.5; 4-64 | 40; 8-62 | 40; 10-60 |
| Smoked packs per year (median; range) | 40; 0.15-154 | 48; 5-94.5 | 50; 1-240 | 50; 2-157.5 |

### Gene Expression Signature Analysis

Clinical data and Gene Expression Quantification data (HTSeq counts) of patients with unpaired RNAseq data (tumor samples without normal samples) was downloaded from the TCGA database using the *TCGAbiolinks* R package. Raw HTSeq counts of tumor samples were normalized by TMM (trimmed mean of M values) method and Log_2_ transformed after filtering to remove genes that consistently have zero or low counts. Univariate Cox Proportional Hazards Regression analysis was performed using *survival* R package [27] to identify survival-related genes. For these survival-related potential biomarker genes (p <= 0.05), Lasso Regularized Cox Model with Leave-One-Out Cross-Validation (LOOCV) was performed to determine a gene expression signature using *glmnet* R package [28]. Multivariate Cox Regression for the signature genes was performed and the predictive performance of the model was scored using *riskRegression* R package [29]. Risk score of each patient was predicted based on multivariate Cox regression model using the *survival* R package. Patients were clustered into high-risk and the low-risk group based on the best cutoff value for ROC, calculated by *cutoff* R package [30].

For the validation of the gene signature, HTSeq counts belonging to the tumor samples of patients who have paired RNAseq data (tumor samples with the paired adjacent normal samples) were downloaded from the TCGA database, filtered, normalized by TMM method and Log_2_ transformed. Multivariate Cox Regression for the signature genes was performed and the predictive performance of the model was scored. Risk score of every patient in the validation group was predicted based on multivariate Cox regression model and each patient was assigned to high-risk or the low-risk group using the best cutoff value for ROC.

Scatter plots showing risk score and survival time of patients were generated by *ggrisk* R package [31] and Kaplan-Meier (KM) survival curves were plotted by *survminer* R package [32] displaying the overall survival difference between the risk groups stratified on the proposed gene signature. ROC curves were plotted for the risk scores based on each gene signature using *survivalROC* R package [33]. Univariate and multivariate cox regression analysis were performed and forest plots were generated for risk score with clinical variables using *survival* and *forestmodel* [34] R packages. Gene and pathway enrichment analysis was performed by *biomaRt* [35] and *clusterProfiler* [36] R packages and plotted by *enrichplot* R package [37]. Heatmap plots was generated using *ComplexHeatmap* R package [38]. Mosaic plots to compare the categorical variables were generated using the *vcd* R package [39,40]. These analyses were performed for LUAD and LUSC patients separately.

### Differential Expression Analysis

Gene Expression Quantification data (HTSeq counts) of both the primary tumor (TP) and the paired normal tissue adjacent to the tumor (NT) was downloaded from the TCGA database. Raw HTSeq counts of both tumor and normal samples were normalized by TMM method after filtering to remove genes which have zero or low counts. Differentially expressed (q < 0.01) genes were determined using *limma* [41] and *edgeR* [42] R packages by limma-voom method with duplicate-correlation function. HUGO symbols and NCBI Gene identifiers of the differentially expressed genes were downloaded using the *biomaRt* R package. Pathway analysis was performed using *clusterProfiler* package and plotted by *enrichplot* R package. Heatmap plots was generated using *ComplexHeatmap* R package. All possible relations between DEGs of LUAD and LUSC risk groups were identified by using *VennDiagram* R package [43]. This analysis was performed for high-risk and low-risk group patients of LUAD and LUSC, separately.

### Active Subnetwork Analysis

Active subnetworks of the differentially expressed genes were determined using *DEsubs* R package [44]. *DEsubs* package accepts the differentially expressed genes output of the *limma* package along with their FDR adjusted p values (q values). *DEsubs* package both computes and plots the active subnetworks. Pathway enrichment of the active subnetwork genes was performed and plotted using *clusterProfiler* and *enrichplot* R packages. Heatmap plots was generated using *ComplexHeatmap* R package. All possible relations between active subnetwork DEGs of LUAD and LUSC risk groups were identified by using *VennDiagram* R package. All the plots and computations were generated for the high-risk and low-risk group patients of the LUAD and LUSC projects, separately.

### Principal Component Analysis (PCA) Analysis

Gene Expression Quantification in tumor samples of LUAD and LUSC risk groups were used for Principal Component Analysis (PCA). Filtered, TMM normalized and Log_2_ transformed HTSeq counts data was used for PCA analysis. All genes in RNAseq data for total patients and test groups, and intersected DEGs and intersected active subnetwork genes for test groups of LUAD and LUSC patients were analyzed by using *PCAtools* R package [45].

### Copy Number Variation Analysis

The Copy Number Variation data of the primary tumor samples of patients was downloaded using *TCGAbiolinks* package (Masked Copy Number Segment as data type). The chromosomal regions which are significantly aberrant in tumor samples were determined and plotted by *gaia* R package [46]. Gene enrichment from genomic regions which have significant differential copy number was performed using *GenomicRanges* [47] and *biomaRt* R packages. R codes used in this analysis were modified from the codes presented at “TCGA Workflow” article [48]. Pathway enrichment analysis of genes which have CNVs was performed and plotted using *clusterProfiler* and *enrichplot* R packages. OncoPrint showing CNVs among patient samples was generated using *ComplexHeatmap* R package. Relations between CNV genes of LUAD and LUSC risk groups were identified by using *VennDiagram* R package. All the computations and the plots were generated for the high-risk and low-risk groups of LUAD and LUSC projects, separately.

### Simple Nucleotide Variations Analysis

The masked Mutation Annotation Format (maf) files of the TCGA mutect2 pipeline in tumor samples were downloaded to obtain the somatic mutations. The maf files are filtered using the *maftools* [49] to obtain the subset of the mutations corresponding to the patient barcodes. Summary plot and oncoplot were generated to summarize the mutation data using *maftools* R package. Somatic mutations were filtered and assigned to either oncogene (OG) or tumor suppressor gene (TSG) groups along with a significance score (q <0.05) using the *SomInaClust* R package [50]. *SomInaClust* computes a background mutation value to identify the hot spots using the known set of somatic mutations in “COSMIC” and the “Cancer Gene Census” (v92) datasets of COSMIC database for GRCh38 [51]. Pathway enrichment analysis of mutated genes was performed and plotted using *clusterProfiler* and *enrichplot* R packages. Relations between SNV genes of LUAD and LUSC risk groups were identified by using *VennDiagram* R package. OncoPlot for significant mutated genes was drawn using *maftools*, and oncoPrint showing SNVs and CNVs together was generated using *ComplexHeatmap* R package. Circos plot showing all non-synonymous SNVs in original data of risk groups and significant CNVs at genome scale were generated using *circlize* R package [52]. All the plots were generated for high-risk and low-risk group patients of LUAD and LUSC projects, separately.

## Results

### Gene Expression Signature Analysis of LUAD and LUSC Patients

In order to identify gene expression prognosis risk model, clinical data and gene expression quantification data of tumor samples of patients with LUAD and h LUSC with unpaired RNAseq data as two separate training groups (Table 1) were downloaded from the TCGA database. A 35-gene expression signature for LUAD and a 33-gene expression signature for LUSC were identified by Lasso Regularized Cox Model with LOOCV after univariate Cox regression analysis. Risk scores of each patient in training groups and test groups were predicted using signature genes, then patients were clustered into high-risk and low-risk groups based on the cutoff values.

The genes of the LUAD expression signature model identified are AC005077.4, AC113404.3, ADAMTS15, AL365181.2, ANGPTL4, ASB2, ASCL2, CCDC181, CCL20, CD200R1, CPXM2, DKK1, ENPP5, EPHX1, GNPNAT1, GRIK2, IRX2, LDHA, LDLRAD3, LINC00539, LINC00578, MS4A1, OGFRP1, RAB9B, RGS20, RHOQ, SAMD13, SLC52A1, STAP1, TLE1, U91328.1, WBP2NL, ZNF571-AS1, ZNF682, ZNF835. Twenty-seven of them are protein-coding genes while two of them are long intergenic non-protein coding RNA (LINC00539, LINC00578), one is antisense RNA (ZNF571-AS1), three of them are pseudogenes (AC005077.4, AC113404.3, OGFRP1) and two of them are novel transcripts (AL365181.2, U91328.1) (Table S1). Pathway enrichment analysis by using *clusterProfiler* R package didn’t give any results for this 35-gene list, therefore enrichment analysis was performed manually using the online KEGG Mapper tool. The genes play role in metabolic pathways, cancer and immune system-related pathways such as Central carbon metabolism in cancer, Glycolysis, Cholesterol metabolism, Amino sugar and Nucleotide sugar metabolism, HIF-1 signaling pathway, TNF signaling pathway, IL-17 signaling pathway, Chemokine signaling pathway and Wnt signaling pathway (Table S2). Multi-variate Cox regression analysis was performed for the signature genes and the predictive performance of the model was scored. The AUC was 0.963 (p = 1.1e-15) for LUAD training group. Risk score of each patient was predicted and patients were clustered into high-risk and low-risk groups based on the cutoff value. Low-risk and high-risk groups have different expression patterns of the signature genes and significantly different survival probabilities (p <0.0001). The prediction power of the risk score is around 0.78 (AUC) for 1, 3, 5 and 8 years for LUAD training group (Figure S1). Risk group clustering is independent from tumor stages because risk groups have also significantly different survival probability for each tumor stage (Figure S2). Vital status is highly correlated with risk groups that high-risk group is positively correlated with death (p = 1.5e-13), while only tumor stage IA and III are associated with risk groups (Figure S3). Risk score has highly significant prognostic ability (HR:2.59, p <0.001) when multivariate Cox regression analysis was performed with other clinical variables (Figure S4, S5).

In order to validate the gene expression signature, gene expression quantification data of tumor samples of patients with LUAD who have paired RNAseq data were downloaded from the TCGA database. Risk scores of each patient in test group were predicted using gene signature lists and patients were clustered into high-risk and low-risk groups based on the best cutoff values for ROC (Figure 1B). Risk groups have differential signature gene expression pattern; high-risk group has lower survival time and higher number of deaths resulting a significantly different survival probability (p <0.0001). Risk score has high prediction powers, 0.97, 0.92, 0.93 and 0.92 (AUC) for 1, 3, 5 and 8 years respectively, for LUAD test group (Figure 1).

**Figure 1.**
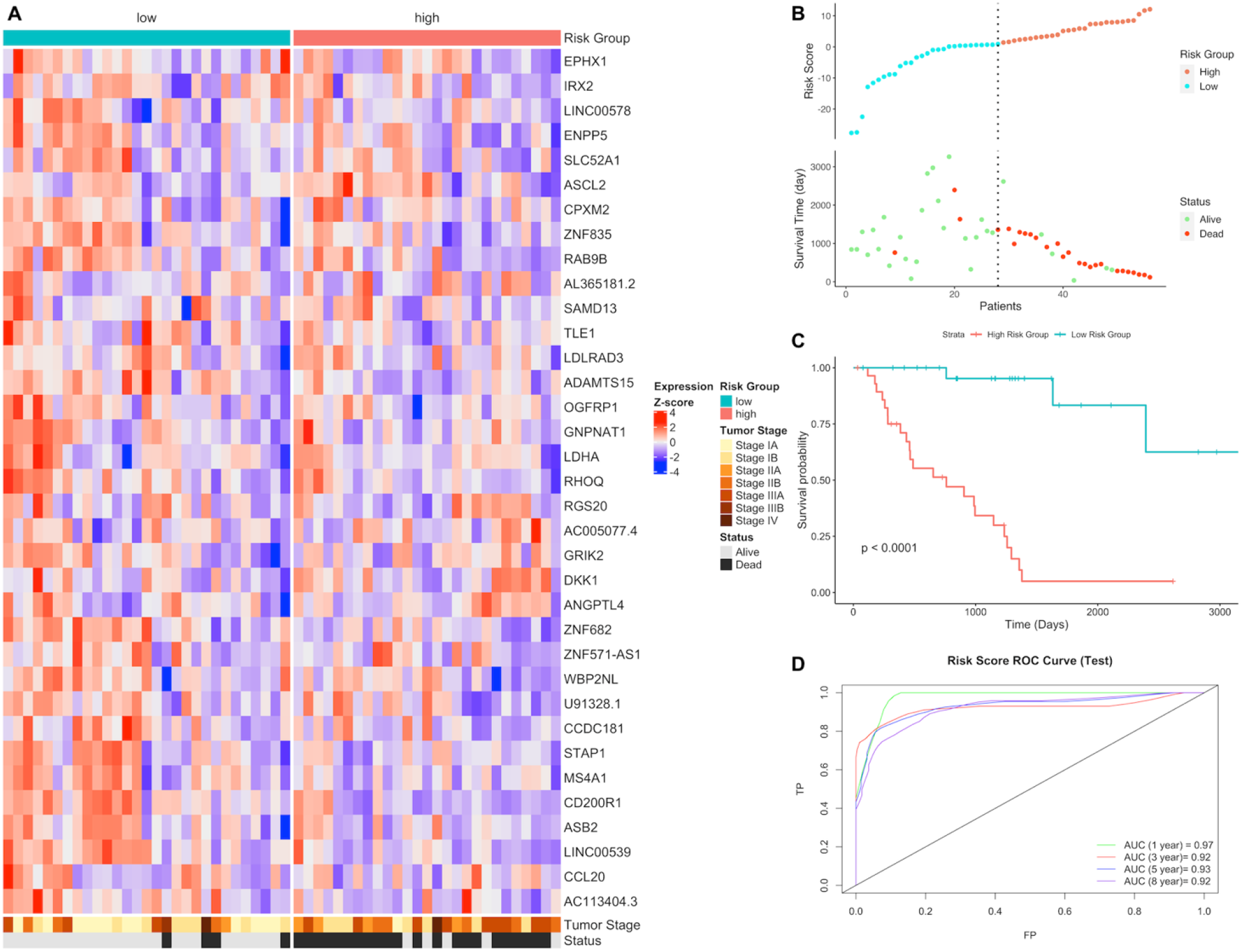
Gene expression signature and risk clustering of LUAD test dataset. Test dataset patients were clustered into high-risk and low-risk groups based on risk scores of patients calculated by predicting the effect of the signature genes of the signature genes expression on overall survival. (**A**) Expression heatmap of the signature genes in tumor samples of LUAD patients in test dataset. (**B**) Scatter plot showing risk scores, survival time and separation point of the patients into risk groups. (**C**) KM survival plot showing the overall survival probability between risk groups. (**D**) ROC curve showing prediction power of risk score in test dataset for 1, 3, 5 and 8 years.

Risk groups have significantly different survival probability for each tumor stage in LUAD test group as well (Figure S6). Vital status is highly correlated with risk groups. High-risk group is positively correlated with death (p = 3.87e-7), while only tumor stage I is positively associated with low-risk group (p = 0.016) (Figure S7). Risk score has highly significant prognostic ability (HR:2.79, p <0.001) as the result of multi-variate Cox regression analysis was performed with other clinical variables (Figure S8).

Expression signature model identified for LUSC includes these genes: AC078883.1, AC096677.1, AC106786.1, ADAMTS17, ALDH7A1, ALK, COL28A1, EDN1, FABP6, HKDC1, IGSF1, ITIH3, JHY, KBTBD11, LINC01426, LINC01748, LPAL2, NOS1, PLAAT1, PNMA8B, RGMA, RPL37P6, S100A5, SLC9A9, SNX32, SRP14-AS1, STK24, UBB, UGGT2, WASH8P, Y_RNA, ZNF160, ZNF703. Twenty-three of them are protein coding genes while two of them are long intergenic non-protein coding RNA (LINC01748, LINC01426), one is antisense RNA (SRP14-AS1), three of them are pseudo-genes (LPAL2, RPL37P6, WASH8P), three of them are novel transcripts (AC106786.1, AC096677.1, AC078883.1) and one is Y RNA (Table S3). They play role in mostly in metabolic pathways, cancer and immunity related pathways such as Arginine and proline metabolism, Glycolysis / Gluconeogenesis, HIF-1 signaling pathway, Non-small cell lung cancer, PD-L1 expression and PD-1 checkpoint pathway in cancer and TGF-beta signaling pathway (Table S4).

The predictive performance score of the signature model is 80.8 (AUC) (p = 1.3e-6) in multivariate Cox regression analysis for LUSC training group. Risk score of each patient was predicted and patients were clustered into high-risk and low-risk groups based on the cutoff value. Low-risk and high-risk groups have different expression patterns of the signature genes and significant difference of survival probability (p <0.0001). The AUC values showing prediction power of the risk score are 0.76, 0.82, 0.87 and 0.92 for 1, 3, 5 and 8 years respectively, for LUSC training group (Figure S9). Risk groups have also significantly different survival probability for tumor stages I, II and III (Figure S10). Risk groups are highly correlated with vital status. High-risk group has highly significant positive correlation with death (p = 8.5e-15), while low-risk group is negatively correlated. Tumor stages didn’t show any association with risk groups (Figure S11). Risk score has highly significant prognostic ability (HR:2.85, p <0.001) when multivariate Cox regression analysis was performed with other clinical variables (Figure S12).

In order to validate the gene expression signature for LUSC, gene expression quantification data of tumor samples of patients with LUSC who have paired RNAseq data were downloaded. Risk scores of each patient in LUSC test group were predicted using gene signature lists and patients were clustered into high-risk and low-risk groups based on the best cutoff values for ROC (Figure 2B). Risk groups have differential signature gene expression pattern; high-risk group has lower survival time and higher number of deaths. Risk groups have significantly different survival probability (p <0.0001). Risk score has high prediction powers, 0.93, 0.95, 0.96 and 0.97 (AUC) for 1, 3, 5 and 8 years respectively, for LUSC test group (Figure 2).

**Figure 2.**
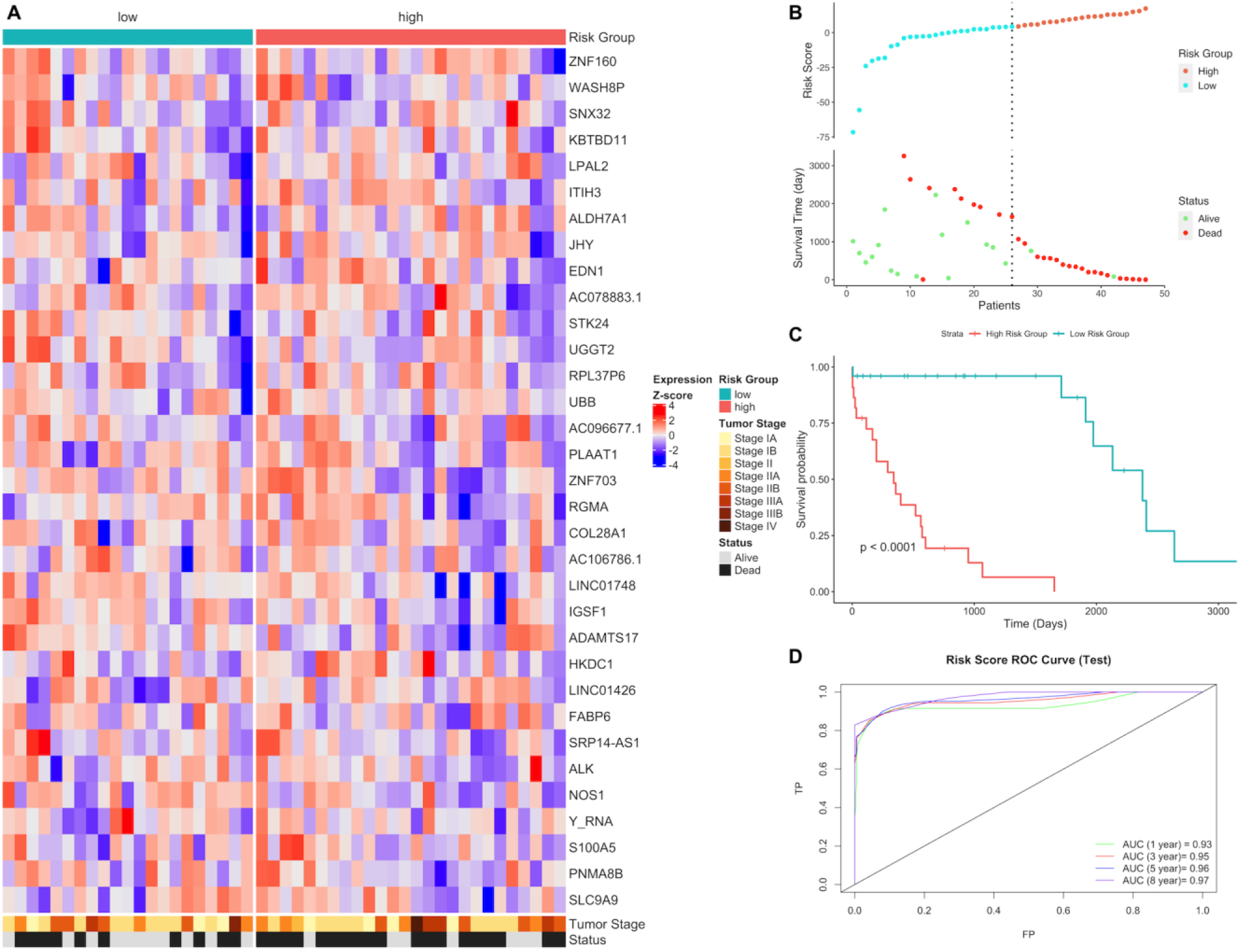
Gene expression signature and risk clustering of LUSC test dataset. Test dataset patients were clustered into high-risk and low-risk groups based on risk scores of patients calculated by predicting the effect of the signature genes of the signature genes expression on overall survival. (**A**) Expression heatmap of the signature genes in tumor samples of LUSC patients in test dataset. (**B**) Scatter plot showing risk scores, survival time and separation point of the patients into risk groups. (**C**) KM survival plot showing the overall survival probability between risk groups. (**D**) ROC curve showing prediction power of risk score in test dataset for 1, 3, 5 and 8 years.

Risk groups have also significantly different survival probability for tumor stages in test group (Figure S13). Vital status isn’t correlated with risk groups of LUSC test group that number of deaths is higher for high-risk group insignificantly (p = 0.07). Tumor stages aren’t associated with risk groups (Figure S14). Risk score has highly significant prognostic ability (HR:2.66, p <0.001) while other clinical variables have no effect on overall survival in multivariate Cox regression analysis (Figure S15).

The expression gene signatures of LUAD and LUSC don’t have any common gene, however they share 8 common pathways which are mostly metabolic pathways: Central carbon metabolism in cancer, Glycolysis / Gluconeogenesis, HIF-1 signaling pathway, Pyruvate metabolism, PPAR signaling pathway, Amino sugar and nucleotide sugar metabolism, TNF signaling pathway and Pathways of neurodegeneration - multiple diseases.

### Differential Expression and Active Subnetwork Analysis of Risk Groups

Gene expression quantification data of both primary tumor and adjacent normal tissues of patients who have paired RNAseq data (test groups) in LUAD and LUSC projects were downloaded from the TCGA database. Differentially expressed (q <0.01) genes (DEGs) were determined in tumor samples according to normal samples for high-risk and low-risk patient groups in test sets of LUAD and LUSC, separately. Then, active subnetworks of DEGs in tumor samples were determined using the DEGs with their q values.

In tumor samples of LUAD low-risk group, the number of the genes which are dysregulated significantly (q <0.01) more than 2-fold is 3615 (2439 down-, 1176 up-regulated) while 3610 genes (2239 down-, 1371 up-regulated) are dysregulated for LUAD highrisk group. LUAD low-risk and high-risk groups have 2745 common differentially expressed genes (Figure S16). Top 20 significant DEGs highlighted as purple at volcano plot in Figure 3A and B are different between LUAD risk groups as dysregulation pattern is different between risk groups albeit the shared 2745 DEGs.

**Figure 3.**
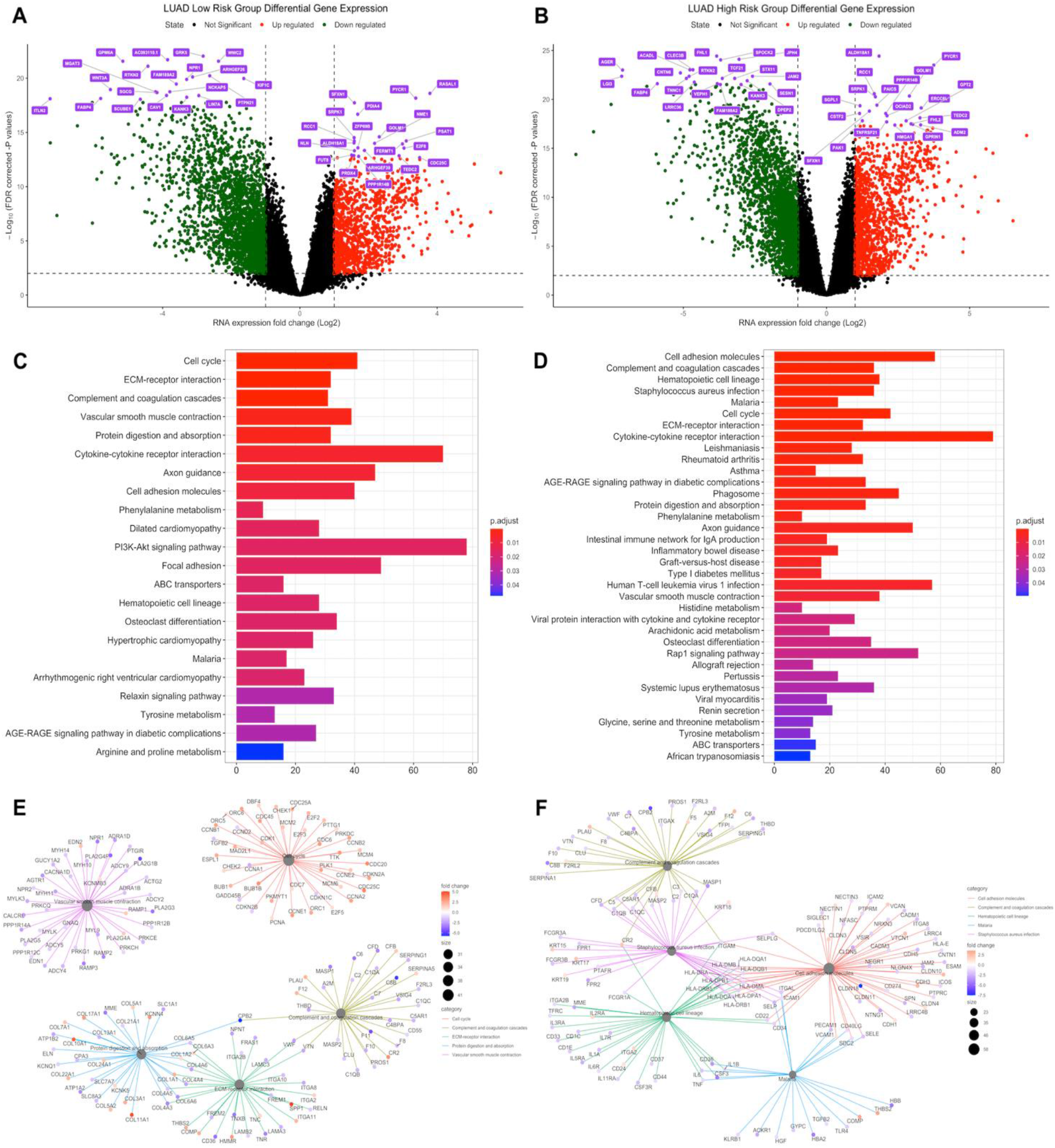
Differential expression analysis of LUAD risk groups. LUAD test dataset patients were clustered into high-risk and low-risk groups based on risk scores of patients and differentially expressed genes in tumor samples were determined based on expressions in normal tissues. (**A**) Volcano plot showing differentially expressed genes more than 2-fold (Log_2_=1) for LUAD low-risk group. Top 20 significant downregulated and upregulated genes are highlighted as purple. FDR corrected p-values threshold is 0.01 (-Log_10_ = 2). Red: Upregulated, Green: Downregulated, Black: Not significant or low than 2-fold. (**B**) Volcano plot showing differentially expressed genes more than two-fold (Log_2_ = 1) for the LUAD high-risk group. Top 20 significant downregulated and upregulated genes are highlighted as purple. FDR corrected p-values threshold is 0.01 (-Log_10_ = 2). Red: Upregulated, Green: Downregulated, Black: Not significant or low than 2-fold. (**C**) Dysregulated pathways of differentially expressed genes for LUAD low-risk group. (**D**) Dysregulated pathways of differentially expressed genes for LUAD high-risk group. (**E**) Top 5 gene-pathway networks of differentially expressed genes with their fold changes for the LUAD low-risk group. (**F**) Top 5 gene-pathway networks of differentially expressed genes with their fold changes for the LUAD high-risk group.

Seven of the signature genes (GNPNAT1, CCDC181, LDHA, ADAMTS15, IRX2, LINC00578, AC005077.4) are dysregulated in both risk groups. ANGPTL4 is upregulated in the high-risk group while MS4A1, GRIK2 and OGFRP1 are upregulated in low-risk group.

Risk groups of LUAD share dysregulated pathways (Figure 3C and D), highly related with cancer, such as Cell cycle, Biosynthesis of aminoacids and Protein digestion and absorption which are upregulated for both risk groups (Figure S17), on the other hand, they also share ECM-receptor interaction, Cell adhesion molecules pathways with immune system-related pathways such as Complement and coagulation cascades and Cytokine-cytokine receptor interaction which are downregulated for both risk groups (Figure S17). However high-risk group has more dysregulated immune system-related pathways such as Allograft rejection, Graft-versus-host disease, Inflammatory bowel disease, Intestinal immune network for IgA production, Rheumatoid arthritis, Staphylococcus aureus infection (Figure 3C and D), which are downregulated pathways in LUAD high-risk group (Figure S17).

Active subnetworks of differentially expressed genes in tumor samples of LUAD risk groups were identified and low-risk group has 191 genes while high-risk group has 168 genes including 112 common genes, which are acting on active subnetworks (Figure S16).

Pathway enrichment of DEGs at active subnetworks shows that the genes playing role in active subnetworks are much more related with cancer pathways such as PI3K-Akt signaling pathway, Ras signaling pathway, Small cell lung cancer, Breast cancer, Gastric cancer, Proteoglycans in cancer and Rap1 signaling pathway (Figure 4). LUAD risk groups have mostly similar cancer related active pathways, however only low-risk group has FoxO signaling pathway and TNF signaling pathway while high-risk group has Estrogen signaling pathway, Growth hormone synthesis, secretion and action with immune system pathways such as Antigen processing and presentation, Intestinal immune network for IgA production and Leukocyte trans-endothelial migration.

**Figure 4.**
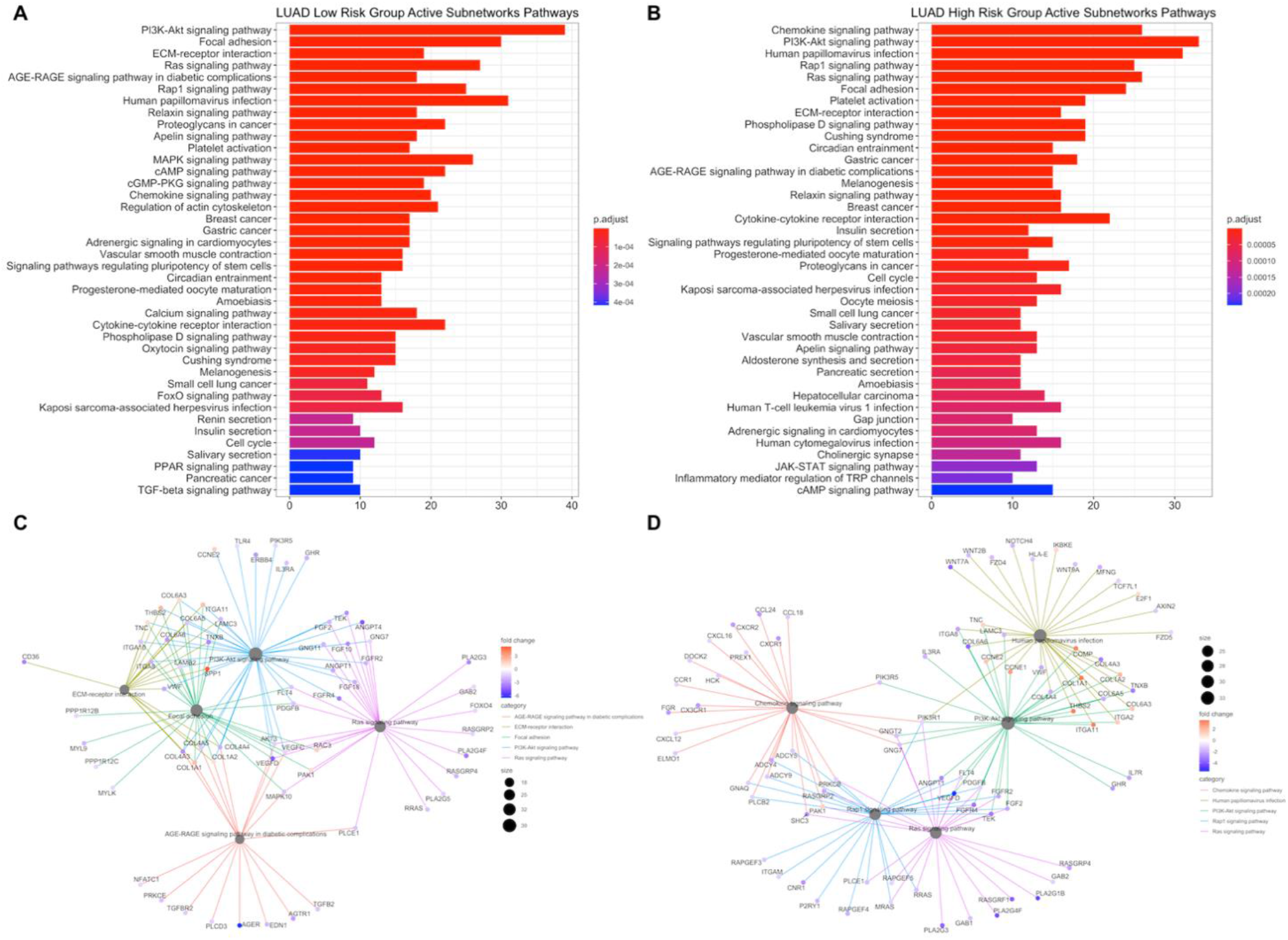
Pathway enrichment of differentially expressed genes at active subnetworks of LUAD risk groups. Active subnetworks were determined by using differential expression analysis results and pathway enrichment analysis was performed for the genes at subnetworks. (**A-B**) Active pathways of differentially expressed genes for LUAD low-risk and high-risk group. (**C-D**) Top 5 gene-pathway networks of active subnetwork genes with their fold changes for the LUAD low-risk group and high-risk group.

The number of dysregulated genes expressed significantly (q <0.01) more than 2-fold in tumor samples of LUSC low-risk group is 5596 (3394 downregulated, 2202 upregulated) while 5403 genes (3338 downregulated, 2065 upregulated) are dysregulated for LUSC high-risk group. LUSC low-risk and high-risk groups have 4562 common differentially expressed genes (Figure S16). Top 20 significant DEGs highlighted at volcano plot in Figure 5A and B include common genes and dysregulation pattern is similar between risk groups.

**Figure 5.**
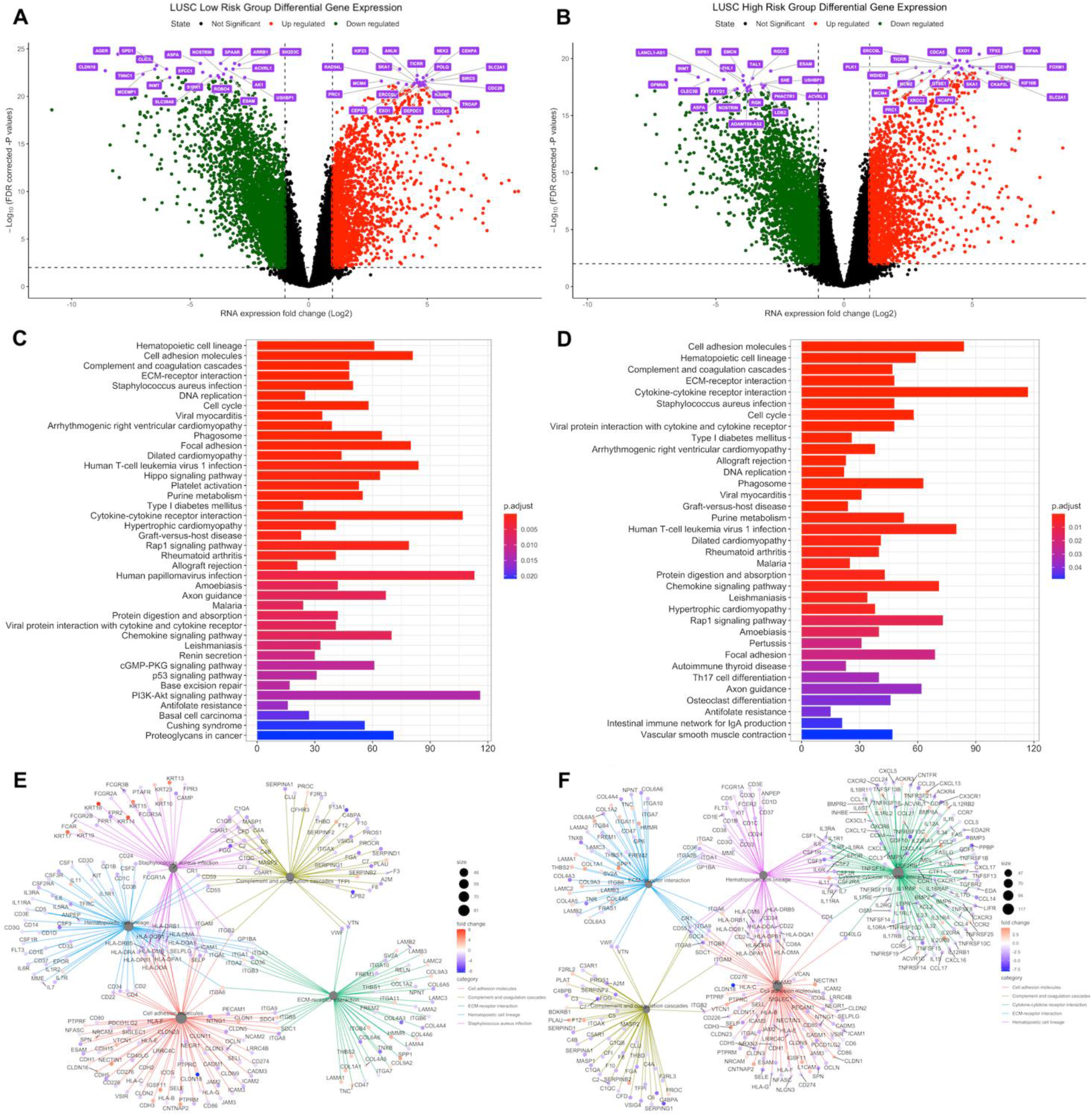
Differential expression analysis of LUSC risk groups. LUSC test dataset patients were clustered into high-risk and low-risk groups based on risk scores of patients and differentially expressed genes in tumor samples were determined based on expressions in normal tissues. (**A**) Volcano plot showing differentially expressed genes more than 2-fold (Log_2_ = 1) for LUSC low-risk group. Top 20 significant downregulated and upregulated genes are highlighted as purple. FDR corrected p-values threshold is 0.01 (-Log_10_ = 2). Red: Upregulated, Green: Downregulated, Black: Not significant or low than 2-fold. (**B**) Volcano plot showing differentially expressed genes more than two-fold (Log_2_ = 1) for LUSC high-risk group. Top 20 significant downregulated and upregulated genes are highlighted as purple. FDR corrected p-values threshold is 0.01 (-Log_10_ = 2). Red: Upregulated, Green: Downregulated, Black: Not significant or low than 2-fold. (**C**) Dysregulated pathways of differentially expressed genes for LUSC low-risk group. (**D**) Dysregulated pathways of differentially expressed genes for LUSC high-risk group. (**E**) Top 5 gene-pathway networks of differentially expressed genes with their fold changes for LUSC low-risk group. (**F**) Top 5 gene-pathway networks of differentially expressed genes with their fold changes for LUSC high-risk group.

LUSC signature genes have 10 common genes (EDN1, JHY, PLAAT1, HKDC1, ITIH3, KBTBD11, RGMA, ZNF703, S100A5, LPAL2) with DEGs of both risk groups. Three of the signature genes, ADAMTS17, IGSF1 and LINC01426, are upregulated in low-risk group; others, NOS1 and SRP14-AS1 are downregulated while Y_RNA is upregulated in the high-risk group.

Risk groups of LUSC have common dysregulated pathways (Figure 5C and D), which are highly related with cancer, such as Cell cycle, DNA replication, Base excision repair, p53 signaling pathway which are upregulated at both risk groups (Figure S18), on the other hand, they also share ECM-receptor interaction, Cell adhesion molecules, Focal adhesion pathways with immune system-related pathways such as Chemokine signaling pathway, Complement and coagulation cascades, Cytokine-cytokine receptor interaction, which are downregulated at both risk groups (Figure S18). However high-risk group has more upregulated metabolic pathways such as Central carbon metabolism in cancer, Protein digestion and absorption, Alanine, aspartate and glutamate metabolism, Arginine and proline metabolism, Cysteine and methionine metabolism, Glutathione metabolism, Ribosome biogenesis in eukaryotes; and down-regulated immune related pathways such as JAK-STAT signaling pathway, TNF signaling pathway, Primary immunodeficiency, T cell receptor signaling pathway distinctly from low-risk group (Figure S18). LUSC low-risk group has downregulated PI3K-Akt signaling pathway, Phenylalanine metabolism, Tyrosine metabolism, Phospholipase D signaling pathway, Proteoglycans in cancer and Tight junction pathways with upregulated Hippo signaling pathway and Small cell lung cancer distinctly from high-risk group (Figure S18).

Active subnetworks of differentially expressed genes in tumor samples of LUSC risk groups has 357 genes for low-risk group while 350 genes for high-risk group including 245 common genes (Figure S16). Active pathways of LUSC risk groups, are highly related with cancer pathways such as PI3K-Akt signaling pathway, Ras signaling pathway, Small cell lung cancer, Proteoglycans in cancer and Rap1 signaling pathway (Figure 6A and B). LUSC risk groups have mostly similar cancer related active pathways, however only low-risk group has Nucleotide excision repair, Adherens junction and Alpha-Linolenic acid metabolism pathways, while high-risk group has cancer and metabolism related pathways such as Basal cell carcinoma, Prolactin signaling pathway, Apoptosis, Mitophagy, Choline metabolism in cancer, Insulin signaling pathway, Carbohydrate digestion and absorption and Central carbon metabolism in cancer with immune system-related Measles and Influenza A pathways.

**Figure 6.**
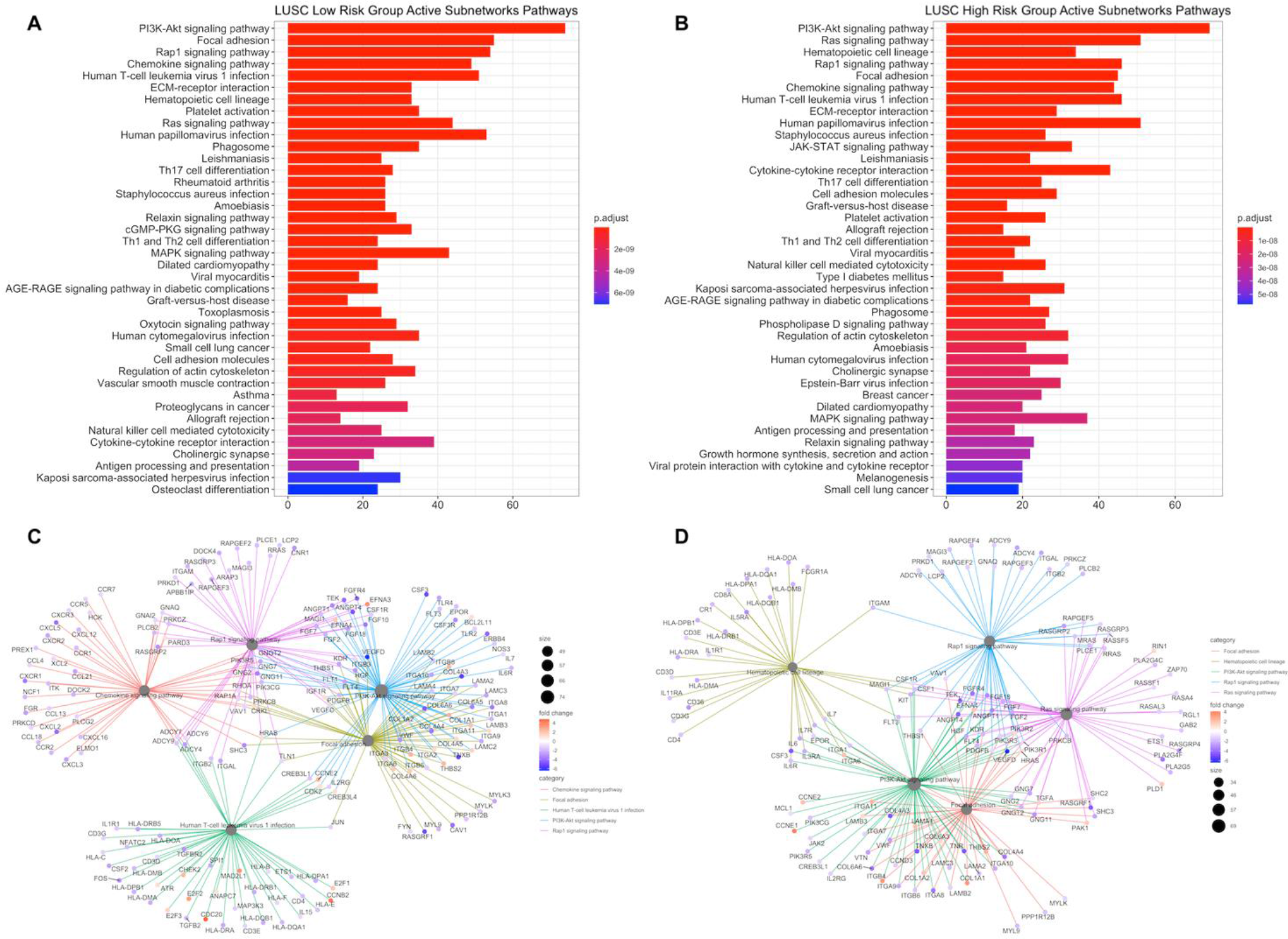
Pathway enrichment of differentially expressed genes at active subnetworks of LUSC risk groups. Active subnetworks were determined by using differential expression analysis results and pathway enrichment analysis was performed for the genes at subnetworks. (**A**) Active pathways of differentially expressed genes for LUSC low-risk group. **(B)** Active pathways of differentially expressed genes for LUSC high-risk group. (**C**) Top 5 gene-pathway networks of active subnetwork genes with their fold changes for LUSC low-risk group. (**D**) Top 5 gene-pathway networks of active subnetwork genes with their fold changes for LUSC high-risk group.

### PCA Analysis of Expression Profiles of LUAD and LUSC risk groups

Gene Expression Quantification data (HTSeq counts) was filtered, TMM normalized and Log_2_ transformed for Principal Component Analysis (PCA). All genes in RNAseq data for total patients and test groups, and intersected DEGs and intersected active subnetwork genes for test groups of LUAD and LUSC patients were analyzed. LUAD and LUSC patients are clustered almost separately based on PC2 when whole transcriptome is used (Figure 7A). However, many patients from different cancer subtypes are placed together and risk groups are not well separated in each cancer cohort (Figure 7A). Test group patients of cancer subtypes are separated better, by using whole transcriptome again, even if some patients are placed in another cancer type cluster. Moreover, risk groups of LUAD are clustered much more far from each other while risk groups of LUSC are clustered closely because LUSC patients in high-risk group are distributed more than patients in low-risk group (Figure 7B). PCA of test group patients give similar result when expression data of common DEGs are used. LUAD and LUSC patients are clustered separately with separation of LUAD risk groups, however, LUAD patients are more distributed (Figure 7C). When only expression of DEGs at active subnetworks are used, neither cancer sub-types nor risk groups are separated well because they have similar expression profiles of genes at active subnetworks (Figure 7D). PC1 and PC2 at Figure 7A and 7B are significantly correlated with gender, tumor stages and smoking data (Figure S19, S20). Significant correlation of the principal components might be due to the known stronger association of LUSC with smoking than LUAD which will be investigated further in our future study.

**Figure 7.**
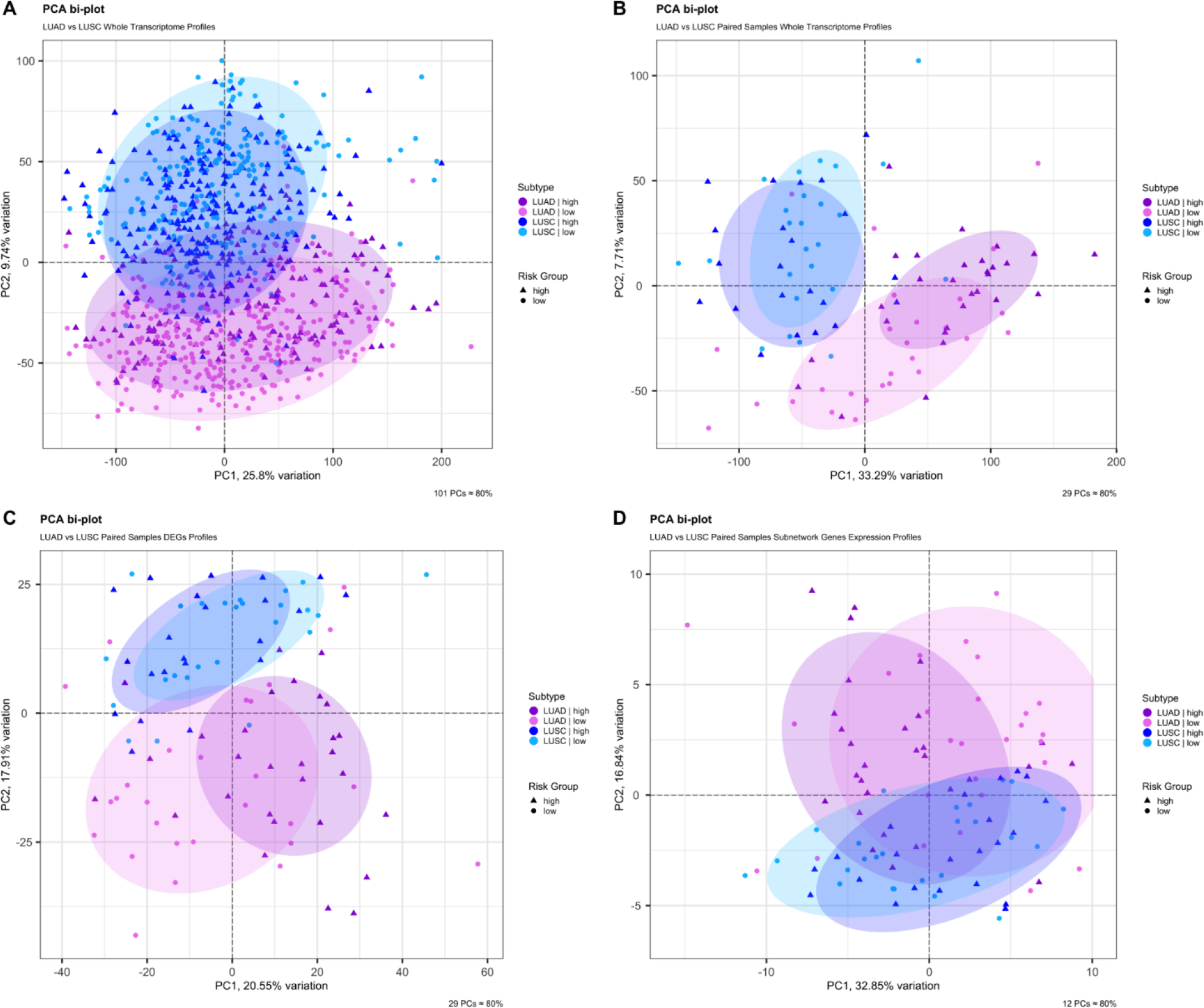
Principal Component Analysis (PCA) of LUAD and LUSC risk groups. PCA was performed by using whole transcriptome for total patients and test groups, and intersected DEGs and intersected subnetwork genes for test groups of LUAD and LUSC patients. (**A**) PCA for all patients including both train and test groups of LUAD and LUSC by using whole transcriptome. (**B**) PCA for test groups of LUAD and LUSC by using whole transcriptome. (**C**) PCA for test groups of LUAD and LUSC by using expression data of intersected DEGs. (**D**) PCA for test groups of LUAD and LUSC by using expression data of DEGs at active subnetworks.

### Copy Number Variations Analysis

The significant aberrant genomic regions in tumor samples of patients were determined and then gene enrichment from genomic regions which have differential copy number was performed. Pathway enrichment analysis of genes which have CNVs was performed and plotted. LUAD low-risk and high-risk groups have different CNV profiles as seen at CNV plots showing amplified or deleted genomic regions on chromosomes. Chromosomes 1, 6, 7, 10, 13, 16, 17, 28 and 20 have different significant aberrant genomic regions (q <0.01) between risk groups (Figure 8A and B). The highest frequencies of the amplified genes are 45%, 48-49% and the deleted genes are 31%, 45% in low-risk and high-risk groups, respectively. Top 25 the highest frequently amplified or deleted genes in tumor samples of risk groups are different and patients in same group may have different aberration patterns (Figure 8C and D). The numbers of the deleted genes and the amplified genes are 10144 and 10412 respectively, in tumor samples of LUAD low-risk group. LUAD high-risk group has 5379 deleted and 8442 amplified genes in tumor samples. Risk groups have 4921 deleted and 6559 amplified genes in common (Figure S23).

**Figure 8.**
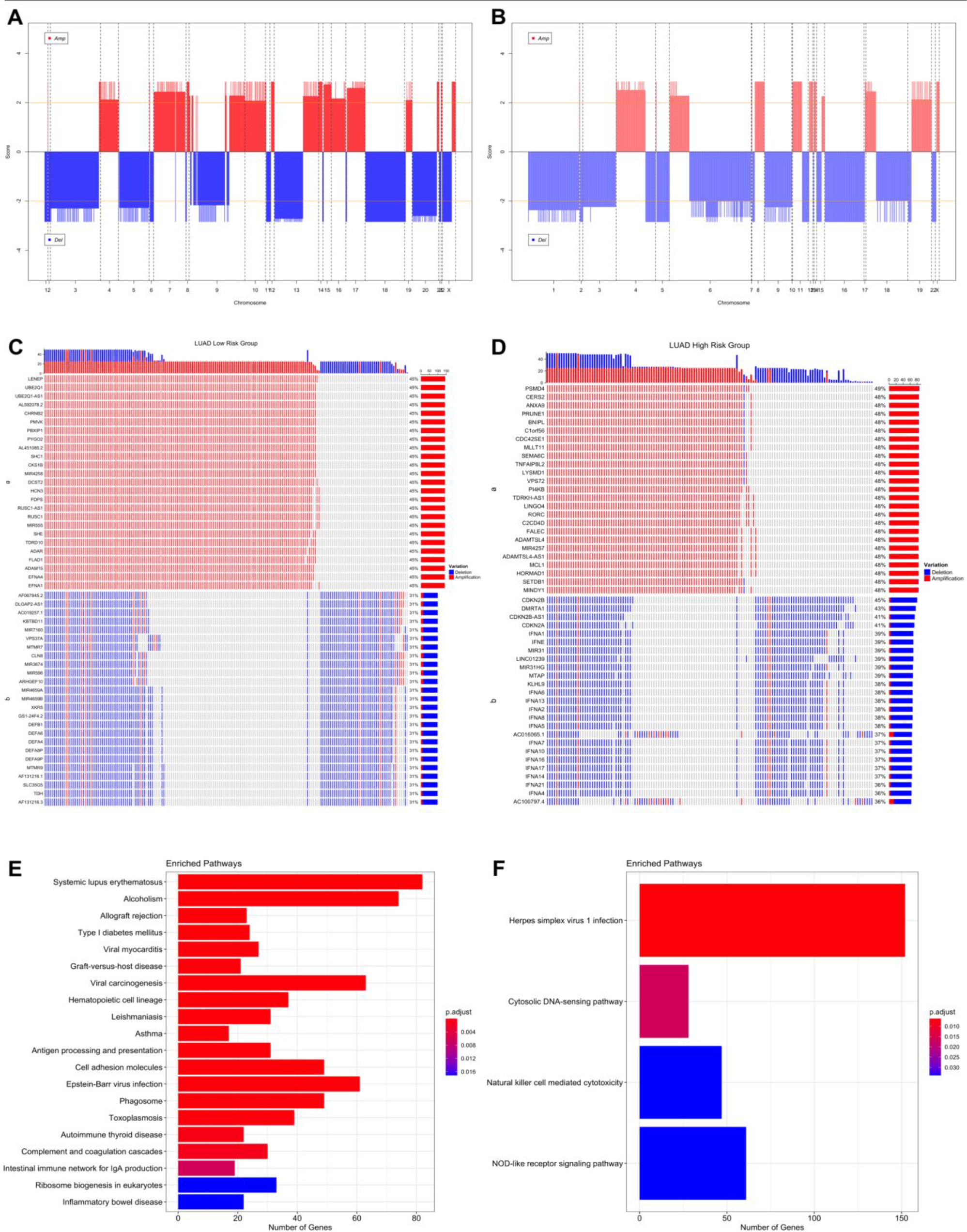
Significant Copy Number Variations (CNVs) of LUAD risk groups. (**A**) CNV plot at genome scale showing amplified or deleted genomic regions on chromosomes of LUAD low-risk group. Score: −Log_10_(q value), Horizontal orange line: 0.01 q value threshold. (**B**) CNV plot of LUAD high-risk group. (**C**) OncoPrint plot showing 25 the highest frequently amplified and deleted genes of LUAD low-risk group. (**D**) OncoPrint plot showing 25 the highest frequently amplified and deleted genes of LUAD high-risk group. (**E**) Pathways of CNV genes of LUAD low-risk group. (**F**) Pathways of CNV genes of LUAD high-risk group.

Pathways of CNV genes are different between LUAD risk groups; mostly immune system pathways such as Allograft rejection, Graft-versus-host disease, Antigen processing and presentation, Complement and coagulation cascades, Inflammatory bowel disease and Viral carcinogenesis pathways have amplified CNVs in low-risk group (Figure S21) while Herpes simplex virus 1, Cytosolic DNA sensing pathway, Natural killer cell mediated cytotoxicity and Nod-like receptor signaling pathways have deleted CNVs (Figure S21) in high-risk group (Figure 8). Complement and coagulation cascades pathway has amplified genes in both risk groups while Natural killer cell mediated cytotoxicity and Nod-like receptor signaling pathways have deleted genes in both risk groups (Figure S21). Low-risk group patients have immune system pathways with amplified genes whereas high-risk group patients have immune system pathways with deleted genes. On the other hand, high-risk group has amplified genes in metabolic pathways such as Gastric acid secretion and Insulin secretion (Figure S21).

LUSC risk groups have different significant aberrant genomic regions obviously on chromosomes 5, 6, 8 and X (Figure 9A and B). The highest frequencies of amplified genes are 84%, 77% and of the deleted genes are 55%, 51% in low-risk and high-risk groups, respectively. LUSC risk groups have higher frequency of amplified genes than deleted genes. Risk groups have common genes from top 25 the highest frequently amplified genes such as SOX2, GHSR, TNFSF10 and miRNAs, miR-7977 and miR-569, with variable frequencies. Risk groups have also common deleted genes such as CDK inhibitors, CDKN2A and CDKN2B, and miR-1284 (Figure 9C and D). LUSC low-risk group has 10720 deleted and 10264 amplified genes while LUSC high-risk group has 9477 deleted and 10250 amplified genes in tumor samples. Risk groups have 7820 deleted and 8659 amplified genes in common (Figure S23).

**Figure 9.**
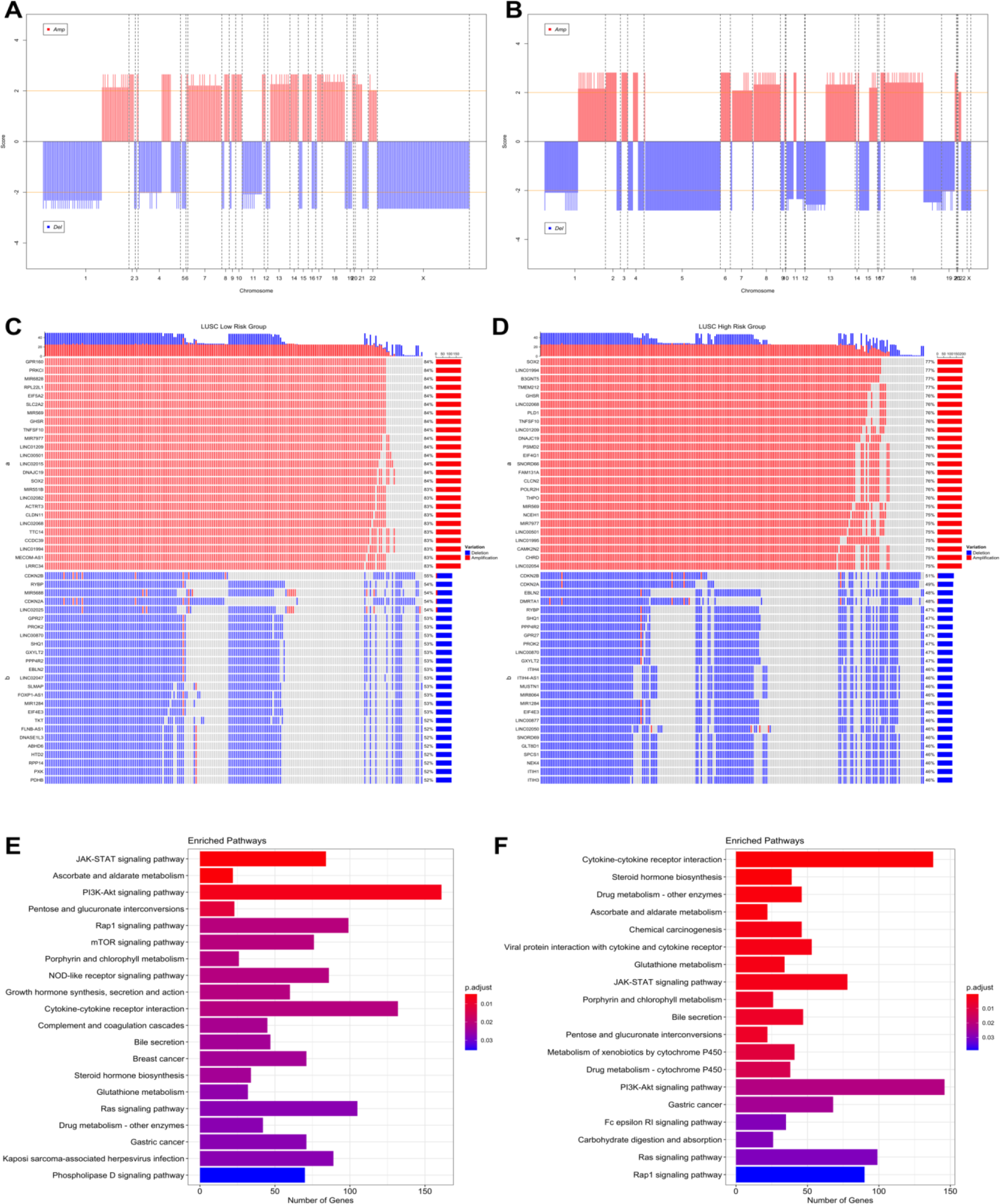
Significant Copy Number Variations (CNVs) of LUSC risk groups. (**A**) CNV plot at genome scale showing amplified or deleted genomic regions on chromosomes of LUSC low-risk group. (**B**) CNV plot of LUSC high-risk group. **(C)** OncoPrint plot showing 25 the highest frequently amplified and deleted genes of LUSC low-risk group. (**D**) OncoPrint plot showing 25 the highest frequently amplified and deleted genes of LUSC high-risk group. (**E**) Pathways of CNV genes of LUSC low-risk group. (**F**) Pathways of CNV genes of LUSC high-risk group.

Pathways of CNV genes highly overlap between LUSC risk groups and they share cancer-related pathways such as PI3K-Akt signaling pathway, JAK-STAT signaling pathway, Ras signaling pathway, Gastric cancer (Figure 9E and F). However, some pathways differ between risk groups, low-risk group has CNVs at mTOR signaling pathway, VEGF signaling pathways and Central carbon metabolism in cancer, while high-risk group has CNVs at Chemical carcinogenesis, Drug metabolism - cytochrome P450, Carbohydrate digestion and absorption pathways (Figure 9E and F). Steroid hormone biosynthesis and Bile secretion pathways have multiple amplified genes while NOD-like receptor signaling pathway has deleted genes, in both risk groups. Only low-risk group has multiple amplified genes at Growth hormone synthesis, secretion and action, and Complement and coagulation cascades pathways. Only high-risk group has amplified genes at Chemical carcinogenesis and Drug metabolism pathways while has deleted genes at Cytokine-cytokine receptor interaction and Fatty acid biosynthesis pathways (Figure S22).

### Simple Nucleotide Variations Analysis

Significantly (q <0.05) mutated genes classified as oncogene (OG) or tumor suppressor gene (TSG) based on TSG / OG scores of the genes and the Cancer Gene Census, were identified for LUAD and LUSC risk groups. COSMIC database was used as a reference mutation database for this analysis and Cancer Gene Census data.

LUAD low-risk group has 15376 mutated genes, while LUAD low-risk group has 12815 mutated genes, 11516 genes of which are common between LUAD risk groups (Figure S28). LUAD patients have wide range of mutation numbers changing from 1518 / 1158 to 10s with median 167 and 172.5 for low-risk and high-risk groups, respectively. Missense mutation is the highest frequent mutation type, and C > A and C > T substitutions are the most frequent ones for both risk groups. LUAD risk groups have similar set of mutated genes with varying frequencies. TP53 is the highest frequently mutated gene with 45% and 53% for low-risk and high-risk groups, and the following ones are MUC16 (39%, 40%) and CSMD3 (38%, 35%) for both groups (Figure S24). SomInaClust analysis was performed to determine driver genes, and 39 genes and 19 genes are strong candidate driver genes for low-risk group and high-risk group, respectively (Table S5, S6). Interestingly, LUAD risk groups share 18 of these driver genes (Figure S28). SomInaClust calculates TSG and OG scores based on background mutation rate and hot spots, then classifies the genes based on TSG / OG scores and cancer gene census data (Figure S26). The driver genes determined in LUAD low-risk group are KRAS, TP53, EGFR, BRAF, STK11, MGA, NF1, RB1, PIK3CA, ATM, RBM10, SETD2, ARID1A, CTNNB1, CMTR2, SF3B1, CSMD3, ATF7IP, KEAP1, HMCN1, EPHA5, ARID2, TTK, SMAD4, KDM5C, SMARCA4, APC, NFE2L2, RIT1, DDX10, LTN1, CDH10, SPTA1, LRP1B, COL11A1, MAP3K12, USH2A, AKAP6 and RASA1. The driver genes determined in LUAD high-risk group are KRAS, TP53, STK11, EGFR, BRAF, RBM10, PIK3CA, SETD2, ARID2, NF1, RB1, MGA, KEAP1, CSMD3, SMARCA4, CTNNB1, KDM5C, IDH1 and ATM (Figure S26; Table S5, S6). TP53 and CSMD3 genes are the most frequently mutated genes with 47%, 56% and 41%, 37% frequencies, respectively for low-risk and high-risk groups (Figure 10 A, and B). More than half of the genes are mutated in less than 12% of patients. For common genes, LUAD high-risk group has mostly higher frequencies. TP53 has differential mutation types, while KRAS has mostly missense mutations. CSMD3 has multi hits (multiple mutations in one patient) in low-risk group than high-risk group. EGFR has in frame deletions in both risk groups and other common genes have similar mutation type pattern between risk groups (Figure 10 A, and B). Pathways of driver mutated genes are highly lung cancer-related pathways such as Non-small cell lung cancer, EGFR tyrosine kinase inhibitor resistance, Platinum drug resistance, MAPK signaling, mTOR signaling, Ras signaling pathway, PI3K-Akt signaling (Figure 10B and C) and other immunologic and metabolic pathways such as Signaling pathways regulating pluripotency of stem cells, FoxO signaling pathway, Rap1 signaling pathway, Central carbon metabolism in cancer, Proteoglycans in cancer, Human T-cell leukemia virus 1 infection, PD-L1 expression and PD-1 checkpoint pathway in cancer and Natural killer cell mediated cytotoxicity pathways, for both risk groups. Many common pathways are enriched because these mutated driver genes play role in many crucial important pathways. However, Wnt signaling pathway and Hippo signaling pathways are mutated only in low-risk group, while Gap junction, GnRH signaling pathway, C-type lectin receptor signaling pathway, T cell receptor signaling pathway, HIF-1 signaling pathway, Growth hormone synthesis, secretion and action and AMPK signaling pathways are mutated only in high-risk group.

**Figure 10.**
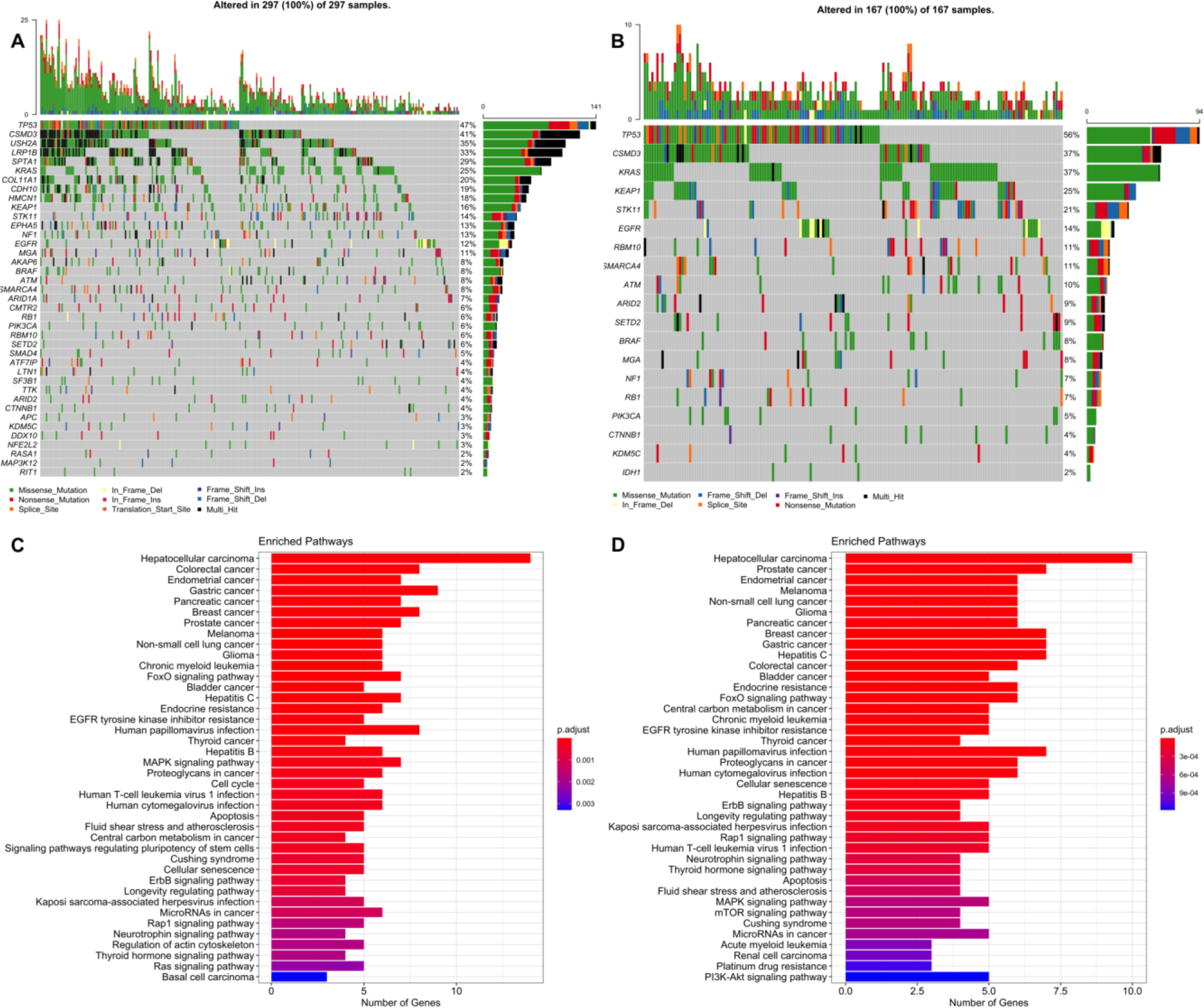
Oncoplot of potential driver genes containing significant SNVs of LUAD risk groups. (**A**) Oncoplot showing significant SNV genes in tumor samples of LUAD low-risk group patients. (**B**) Oncoplot showing significant SNV genes in tumor samples of LUAD high-risk group patients. (**C**) Pathway enrichment of the significant SNV genes of LUAD low-risk group. (**D**) Pathway enrichment of the significant SNV genes of LUAD high-risk group.

LUSC low-risk group has 14038 mutated genes, while LUSC low-risk group has 14616 mutated genes, and 11947 genes are common (Figure S28). LUSC patients have a range of mutation numbers from 2300 / 1488 to 10s with median 201 for low-risk and high-risk groups, respectively. Missense mutation is the highest frequent mutation type, and C > A and C > T substitutions are the most frequent ones for both risk groups. LUSC risk groups have overlapping list of mutated genes with varying frequencies. TP53 is the highest frequently mutated gene with 80% and 78% for low-risk and high-risk groups, and the following ones are CSMD3 (42%, 42%) and MUC16 (39%, 40%) for both groups (Figure S25). As candidate driver genes, 30 genes and 19 genes were identified for low-risk group and high-risk group, respectively (Table S7, S8). LUSC risk groups share 14 of these driver genes (Figure S28). The driver genes determined in LUSC low-risk group are TP53, KMT2D, NFE2L2, PIK3CA, CDKN2A, PTEN, RB1, FAT1, ARID1A, NF1, RASA1, CUL3, KDM6A, NRAS, KRT5, ZNF750, EP300, FGFR3, TAOK1, CSMD3, NSD1, HRAS, SI, PDS5B, KRAS, KEAP1, API5, HNRNPUL1, SLC16A1, FBXW7. The driver genes determined in LUSC high-risk group are TP53, NFE2L2, PIK3CA, KMT2D, FAT1, CDKN2A, RB1, PTEN, NOTCH1, ARID1A, RASA1, NF1, KMT2C, BRAF, PIK3R1, CSMD3, STK11, HRAS, KEAP1 (Figure S27; Table S7, S8). TP53 (83%, 82%), CSMD3 (44%, 44%) and KMT2D (25%, 23%) are most frequent mutated genes for low-risk and high-risk groups (Figure 11 A, and B). For common genes, risk groups have similar frequencies. TP53 and KMT2D genes have differential mutation types, while CSMD3 has mostly missense and multi hit mutations. CDKN2A has mostly truncating mutations in both risk groups and other common genes have similar mutation type pattern between risk groups (Figure 11 A, and B). Pathways of driver mutated genes are highly lung cancer-related pathways such as Non-small cell lung cancer, EGFR tyrosine kinase inhibitor resistance, Platinum drug resistance, MAPK signaling and Ras signaling (Figure 11B and C) and other immunologic and metabolic pathways such as FoxO signaling pathway, Central carbon metabolism in cancer, Proteoglycans in cancer, Hepatitis B, Hepatitis C, PD-L1 expression and PD-1 checkpoint pathway in cancer for both risk groups. Many common pathways are enriched because these mutated driver genes play role in many crucial important pathways. However, Gap junction and Ubiquitin mediated proteolysis pathways are mutated only in low-risk group, while HIF-1 signaling and TNF signaling pathways are mutated only in high-risk group.

**Figure 11.**
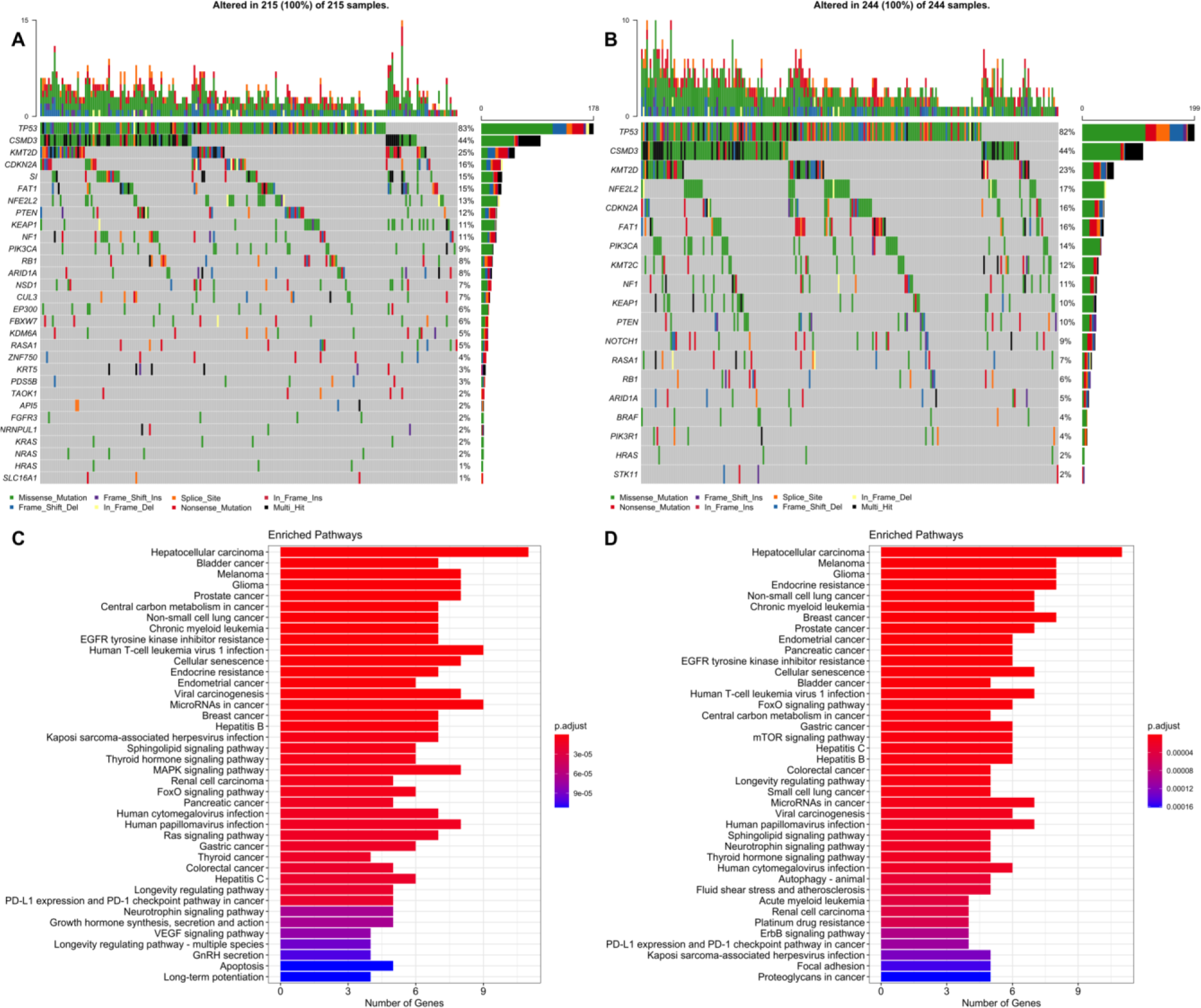
Oncoplot of potential driver genes containing significant SNVs of LUSC risk groups. (**A**) Oncoplot showing significant SNV genes in tumor samples of LUSC low-risk group patients. (**B**) Oncoplot showing significant SNV genes in tumor samples of LUSC high-risk group patients. (**C**) Pathway enrichment of the significant SNV genes of LUSC low-risk group. (**D**) Pathway enrichment of the significant SNV genes of LUSC high-risk group.

When venn diagram is drawn by using all driver genes, all cancer and risk groups have TP53, CSMD3, KEAP1, NF1, RB1 and PIK3CA mutations. KRAS, STK11, BRAF, ARID1A, NFE2L2 and RASA1 genes are shared by 3 different groups. LUAD high-risk group has only IDH1 oncogene as different from LUAD low-risk group while LUSC highrisk group has KMT2C, NOTCH1 and PIK3R1 tumor suppressor genes as different from LUSC low-risk group. EGFR, MGA and SMARCA4 are not driver genes in LUSC while CDKN2A, PTEN, HRAS and FAT1 are not driver genes in LUAD groups (Figure 12).

**Figure 12.**
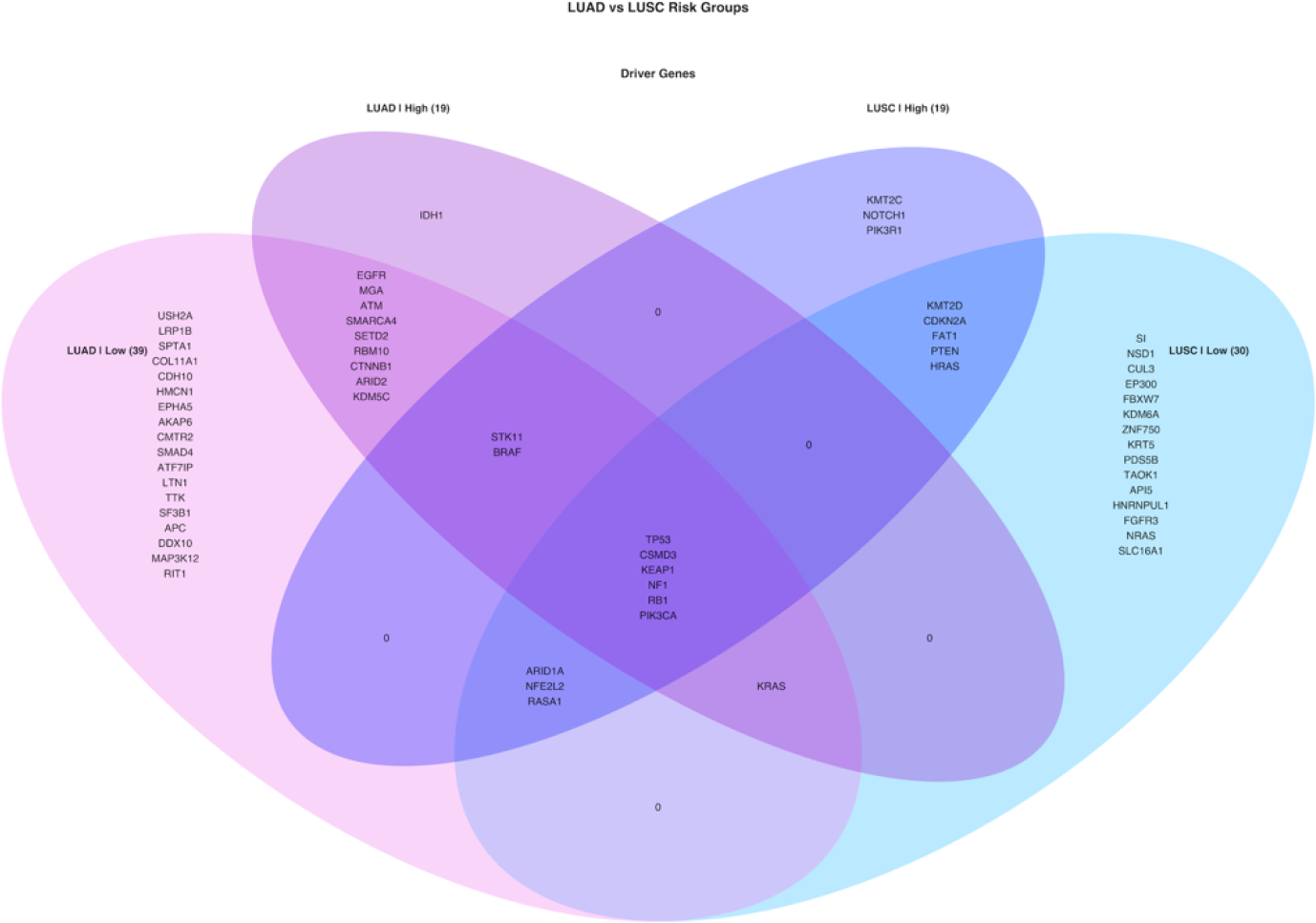
Venn diagram of driver genes containing SNVs.

Significant SNVs and CNVs on driver genes are codisplayed as OncoPrint. Although there exist some genes with both SNVs and significant CNVs while others have only SNVs. Moreover, some patients have only SNVs or only CNVs or both for a specific driver gene.

TP53, STK11, KEAP1, SMARCA4 and MGA genes have deletions while CSMD3 and PIK3CA genes have amplification beside SNVs in both LUAD risk group. KRAS and EGFR genes have amplification in high-risk group however they don’t have significant CNVs in low-risk group. Oncogenes tend to have amplifications while tumor suppressor genes tend to have deletions in both risk groups with exceptions (CSMD3, CDH10, HMCN1, AKAP6 and CTNNB1) (Figure 13).

**Figure 13.**
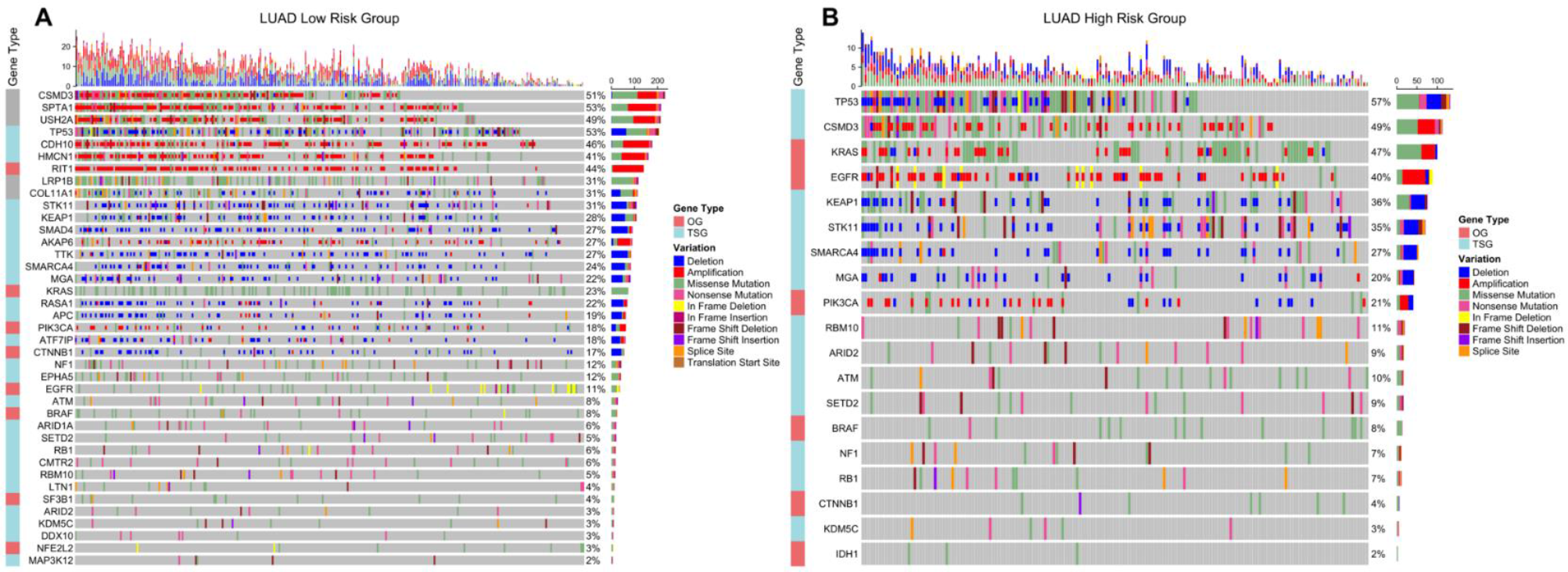
OncoPrint of driver genes containing significant SNVs and CNVs in LUAD risk groups. Significant SNVs and CNVs are plotted together on potential driver genes in tumor samples of LUAD risk groups. (**A**) OncoPrint of driver genes in LUAD low-risk group. (**B**) OncoPrint of driver genes in LUAD high-risk group.

OncoPrints in Figure 14 show that TP53, CDKN2A, FAT1, RASA1, ARID1A and HRAS genes have deletions while only PIK3CA gene has amplification beside SNVs in both LUSC risk groups. PIK3R1, KEAP1 and STK11 genes have deletions only in high-risk group while SI, CSMD3, ZNF750, KRAS genes have amplification and NSD1, FGFR3, PTEN, SLC16A1, NRAS and CUL3 have deletion only in low-risk group. Oncogenes tend to have amplifications while tumor suppressor genes tend to have deletions in both risk groups with exceptions (CSMD3, FGFR3, ZNF750, NRAS, HRAS, KEAP1) (Figure 14).

**Figure 14.**
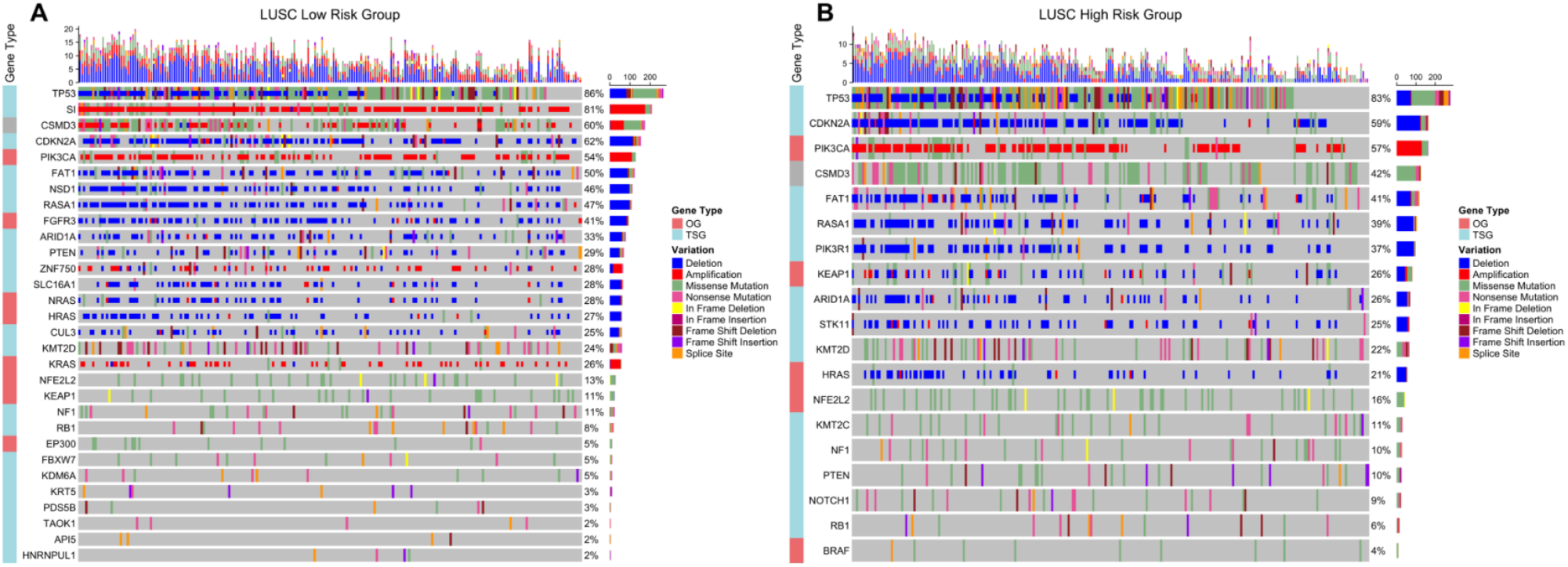
OncoPrint of driver genes containing significant SNVs and CNVs in LUSC risk groups. Significant SNVs and CNVs are plotted together on potential driver genes in tumor samples of LUSC risk groups. (**A**) OncoPrint of driver genes in LUSC low-risk group. (**B**) OncoPrint of driver genes in LUSC high-risk group.

Circos plots showing all non-synonymous SNVs in original data of risk groups and significant CNVs at genomic scale on chromosomes, were drawn to show the genomic alterations between risk groups of LUAD and LUSC.

LUAD low-risk group has more genome wide CNVs and SNVs than high-risk group. Low-risk group has more genomics regions containing missense, nonsense and frame shift insertions/deletions mutations. Moreover, low-risk group has extra deletions on chromosomes 1, 3, 5, 6, 12, 15 and X with extra amplifications on chromosomes 6, 10, 14 and 20. High-risk group has extra amplifications on chromosomes 7, 11, 12 and 17. The CNVs of high-risk group are localized mostly on 1, 3, 5, 6, 7, 8 and 17 whereas low-risk group has CNVs on more chromosomes (Figure 15).

**Figure 15.**
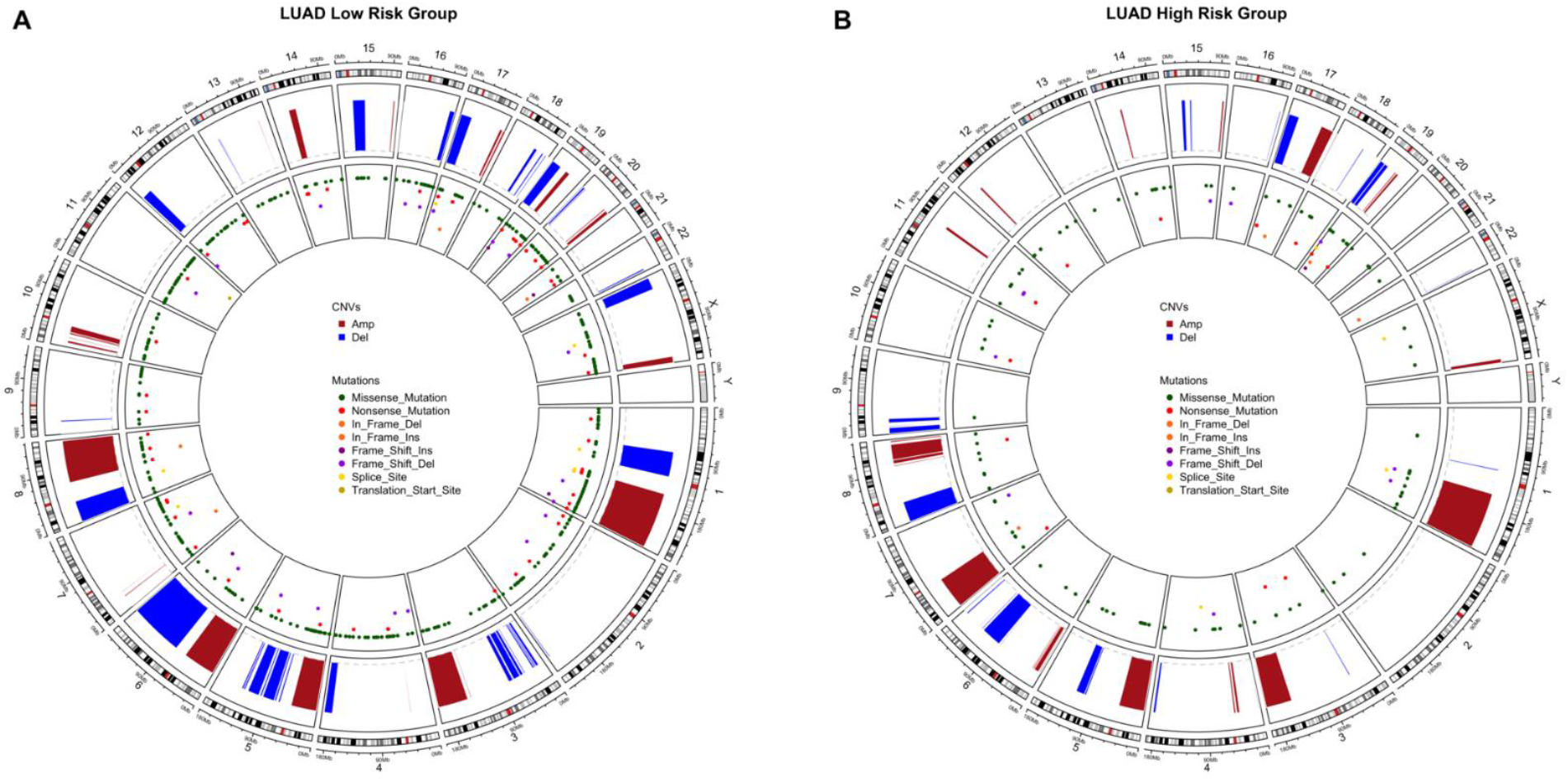
Circos plot of chromosome regions containing all SNVs and CNVs in LUAD risk groups. Significant CNVs (q <0.01) and all SNVs in original data are plotted together on chromosome regions in tumor samples of LUAD risk groups. (**A**) Circos plot of LUAD low-risk group. (**B**) Circos plot of LUAD high-risk group.

LUSC high-risk group has more genomics regions containing missense and nonsense mutations than low-risk group. However, they have similar amount of CNVs although with different localizations. High-risk group has extra amplifications on chromosomes 4, 6 and 11; has extra deletions on chromosomes 15, 19 and X. Low-risk group has only extra deletions on chromosomes 1, 5, 6, 11 and 16 (Figure 16).

**Figure 16.**
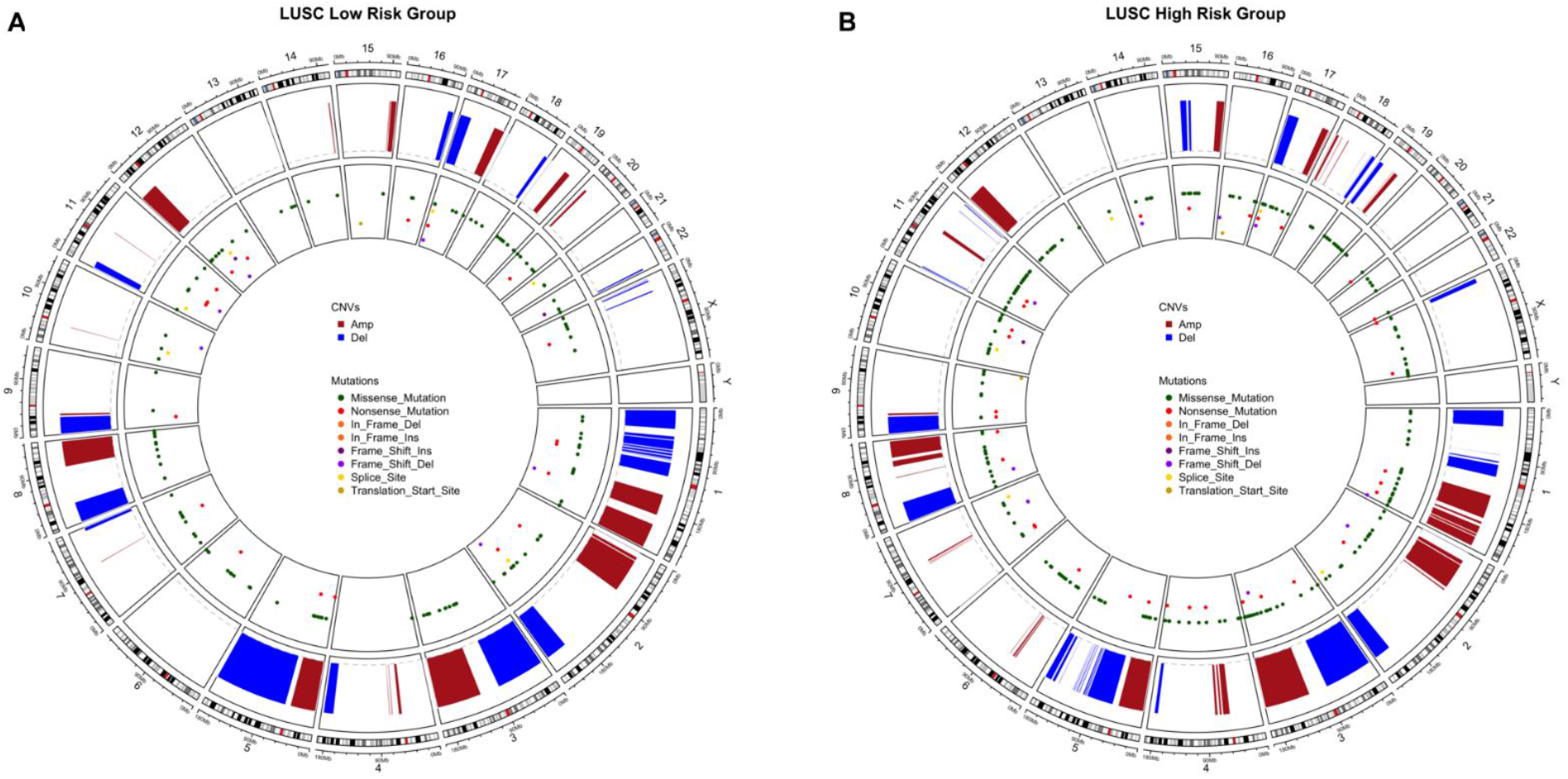
Circos plot of chromosome regions containing all SNVs and CNVs in LUSC risk groups. Significant CNVs (q <0.01) and all SNVs in original data are plotted together on chromosome regions in tumor samples of LUSC risk groups. (**A**) Circos plot of LUSC low-risk group. (**B**) Circos plot of LUSC high-risk group.

## Discussion

In order to profile the genetic differences between risk groups of LUAD and LUSC, gene expression signatures were generated and the patients were clustered into low-risk and high-risk groups and then significant DEGs, DEGs at active subnetworks, CNVs and SNVs were identified in each risk group. The biological alterations for these data types were compared between risk groups and between lung cancer subtypes.

Expression sffPCignature for LUAD consists of 35 gene which 27 of are protein-coding genes while 2 of them are long intergenic non-protein coding RNA, 1 is antisense RNA, 3 of them are pseudogenes and 2 of them are novel transcripts. Many of the coding genes are lung cancer or other cancer types related such as ADAMTS15, CCDC181 and IRX2 with potential tumor suppressor roles; ANGPTL4, Asb2, ASCL2, CCL20, DKK1, GRIK2, LDHA, RGS20, RHOQ, TLE1, WBP2 and ZNF835 with potential oncogenic roles; and CD200, CD200R1, EPHX1, GNPNAT1, LDLRAD3, STAP1, LINC00578 with prognostic potential. Moreover, MS4A1 is dysregulated in asbestos related lung squamous carcinoma, RAB9B is target of miR-15/16 which are highly related with lung cancer, LINC00539 is related with tumor immune response while long non-coding RNA OGFRP1 regulates non-small cell lung cancer progression [53]. The remaining signature genes, CPXM2, ENPP5, SAMD13, SLC52A1, ZNF682, ZNF571-AS1 and U91328.1, have not been related with carcinoma, yet. However, they showed highly prognostic power through risk score to distinguish low-risk and high-risk of overall survival in LUAD.

LUSC gene expression signature including 33 genes which are mostly noncharacterized or oncogenic. ALDH7A1, ALK, EDN1, FABP6, HKDC1, IGSF1, KBTBD11, NOS1, SLC9A9, UBB, ZNF703 have been shown with oncogenic relations while RGMA and STK24 genes with candidate tumor suppressors. ITIH3 and S100A5 has been related with prognostic biomarker potentials. Other cancer related genes are WASH8P, LINC01748, LPAL2, SRP14-AS1. Long intergenic non-protein coding RNA, LINC01426, promotes cancer progression via AZGP1 and predicts poor prognosis in patients with LUAD [54]. COL28A1 has prognostic values and is a potential tumor suppressor identified by *SomInaClust* R package in our previous analysis [55]. Many of the genes such as ADAMTS17, JHY, PLAAT1, PNMA8B, RPL37P6, SNX32, UGGT2 and Y_RNA have not been related with any cancer, yet.

Gene expression signatures of LUAD and LUSC share 8 pathways which are mostly metabolic pathways. LUAD signature plays role in immune related pathways as different from those in LUSC. However, pathway enrichment shows us that risk prediction works on metabolic pathways, therefore if we put a name to important mutations as driver mutations, in this case we can say that active metabolic gene expressions are the fuel of the cancer. The differential expression on them with immune system effect in count can hold the passage of cancer.

High risk groups of both LUAD and LUSC have more immune pathways including downregulated genes and metabolic pathways including upregulated genes. On the other hand, low risk groups have both upregulated and down regulated genes on cancer-related pathways. Although LUAD and LUSC seem to have similar characteristics of risk groups, close signature gene pathways and similar differential expression pathways sharing 2106 DEGs in total, they are displayed separately in PCA, especially at analysis of test groups. At CNV level both risk groups and cancer subtypes have huge number of genes with amplifications or deletions which can cause genomic instability and uncontrolled regulation. Both LUAD and LUSC risk groups have important gene alterations such as CDKN2A and CDKN2B deletions with different frequencies while SOX2 amplification in LUSC and PSMD4 amplification in LUAD. CNVs also play role in metabolic and immune related pathways which can differ between risk groups and cancer subtypes. If we look from a higher perspective, LUAD low risk group has much more CNVs and SNVs on its genome than high-risk group. So, the high-risk group seems more oncogene dependent group even if it has less driver SNV and CNV mutations. On the other hand, LUSC high-risk group has more SNVs than low-risk group while CNVs don’t vary too much as in LUAD. SNV analysis gives similar results with literature for example EGFR and KRAS mutations are mutually exclusive in LUAD samples that is confirmed again. Additionally, EGFR, MGA, SMARCA4, ATM, RBM10 and KDM5C genes are mutated only in LUAD but not in LUSC. On the other way, CDKN2A, PTEN and HRAS genes are mutated only in LUSC. In general, low risk groups have more mutated genes for both LUAD and LUSC samples. When SNV and CNV genes are plotted together, it can be seen that LUAD highrisk group has obvious oncogene amplifications and tumor suppressor deletions, while LUAD low-risk group has both tumor suppressor deletions and tumor suppressor amplifications with a few oncogene amplifications. This SNV and copy number differential pattern can cause differential gene expression profiles and characteristics of tumor. LUSC patients have mostly deletions on driver genes with only PIK3CA and KRAS oncogene amplifications. Both LUSC risk groups have obvious TP53 and CDKN2A deletions, but CSMD3 amplification does not occur in LUSC high-risk group. Again, only these driver genes which have differential alterations and frequencies can create the risk difference based on gene expression levels.

## Conclusions

This study has been performed to profile the genomic and transcriptomic differences not only between LUAD and LUSC also between risk groups to understand the driving differences between them. It is not still patient based analysis, but we are getting closer, parallel with development of targeted therapies against the biomarkers or important alterations which identified more specific cluster-based analyses. Nowadays, many groups and government institutions are working on the integration of the drug bioactivity and molecular data to investigate the potential of a more effective molecularly targeted therapeutics for individual patients following the personalized medicine trend worldwide.

## Supplementary Materials

The supplementary data is available online at www.mdpi.com/xxx/s1

## Author Contributions

Methodology, T.Z.; formal analysis, T.Z.; resources, T.Z., T.Ö-S.; data curation, T.Z.; writing—original draft preparation, T.Z.; writing—review and editing, T.Ö-S.; visualization, T.Z.; project administration, T.Ö-S. All authors have read and agreed to the published version of the manuscript.

## Funding

“T.Z. and T.Ö-S were partially funded by Turkish National Institutes of Health (TÜSEB) grant number 4583”.

## Informed Consent Statement

Not applicable.

## Data Availability Statement

The datasets supporting the conclusions of this article are publicly available and can be downloaded from TCGA data portal (https://portal.gdc.cancer.gov) or by using *TCGAbiolinks* R package [26]. The R code used in this study is available upon request.

## Acknowledgments

In this section, you can acknowledge any support given which is not covered by the author contribution or funding sections. This may include administrative and technical support, or donations in kind (e.g., materials used for experiments).

## Conflicts of Interest

The authors declare no conflict of interest.

## Notes

### Competing Interest Statement

The authors have declared no competing interest.

### Summary of Updates

Journal name is removed

