## Supplementary Material for "Comprehensive profiling of genomic and transcriptomic differences between risk groups of lung adenocarcinoma and lung squamous cell carcinoma"

### SUPPLEMENTARY DATA

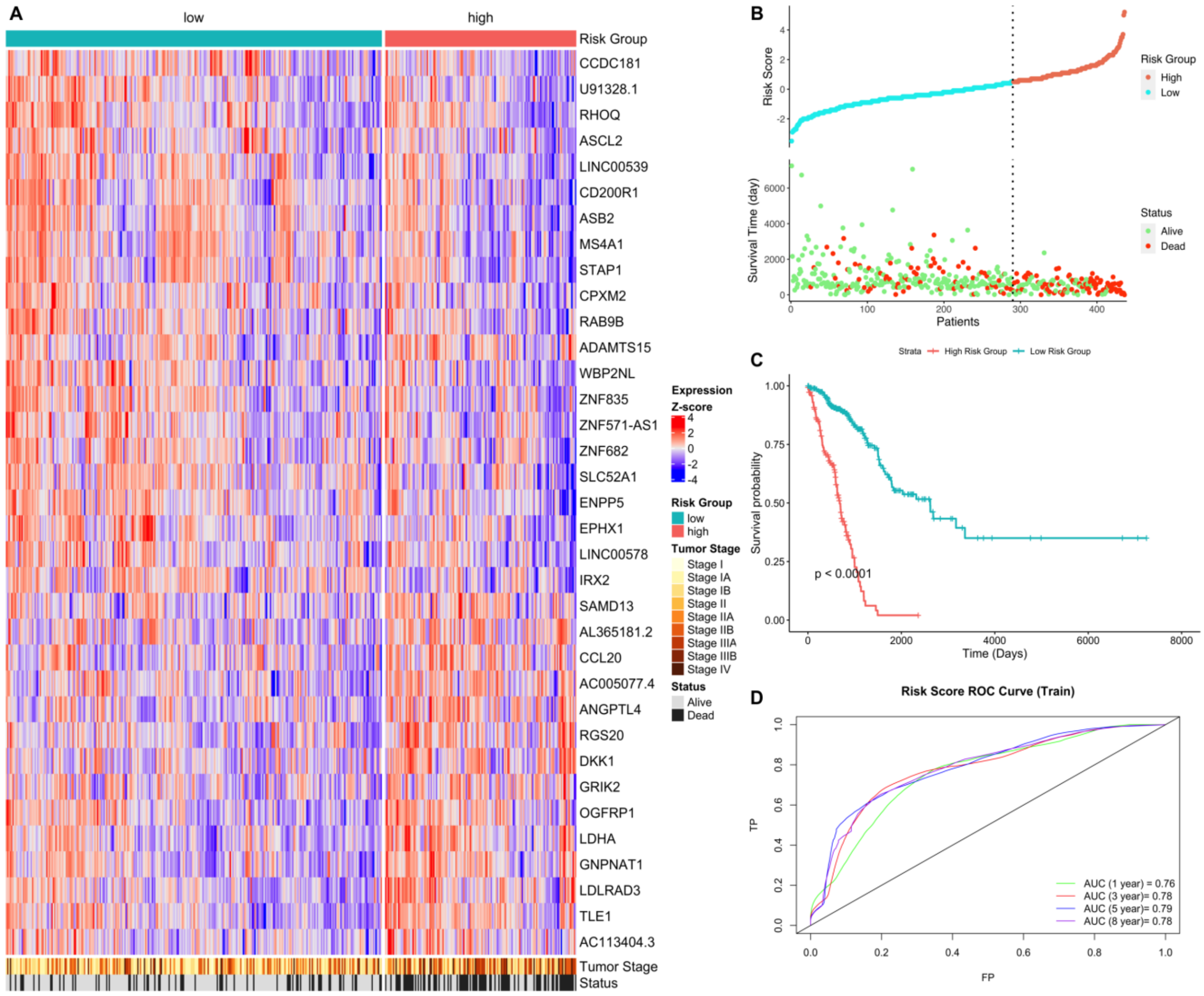


**Figure S1.** Gene expression signature and risk clustering of LUAD training dataset. (**A**) Expression heatmap of the signature genes in tumor samples of LUAD patients in train dataset. Train dataset patients were clustered into high-risk and low-risk groups; (**B**) Scatter plot showing risk scores, survival time and separation point of the patients into risk groups. (**C**) KM survival plot showing the overall survival probability between risk groups. (**D**) ROC curve showing prediction power of risk score in train dataset for 1, 3, 5 and 8 years.

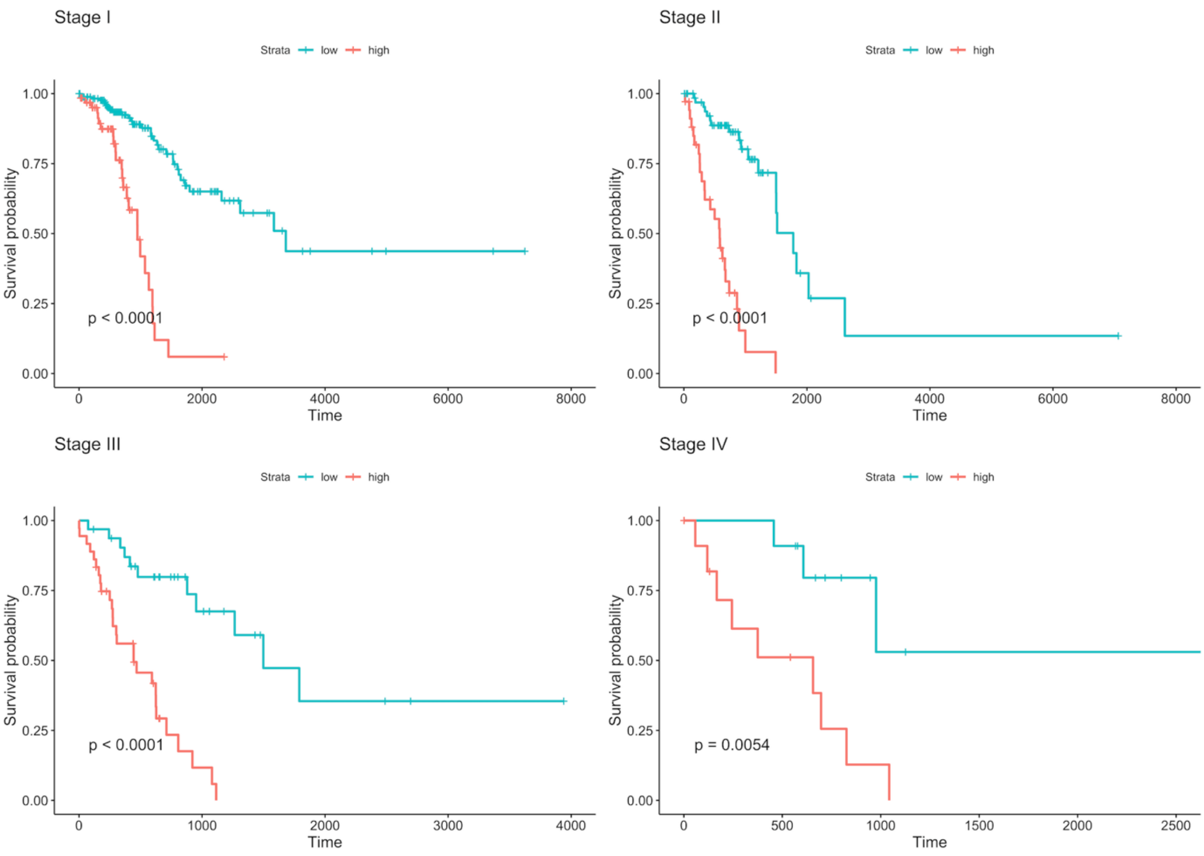


**Figure S2.** Survival analysis of risk groups clustered by using signature gene expression at different tumor stages in LUAD training dataset.

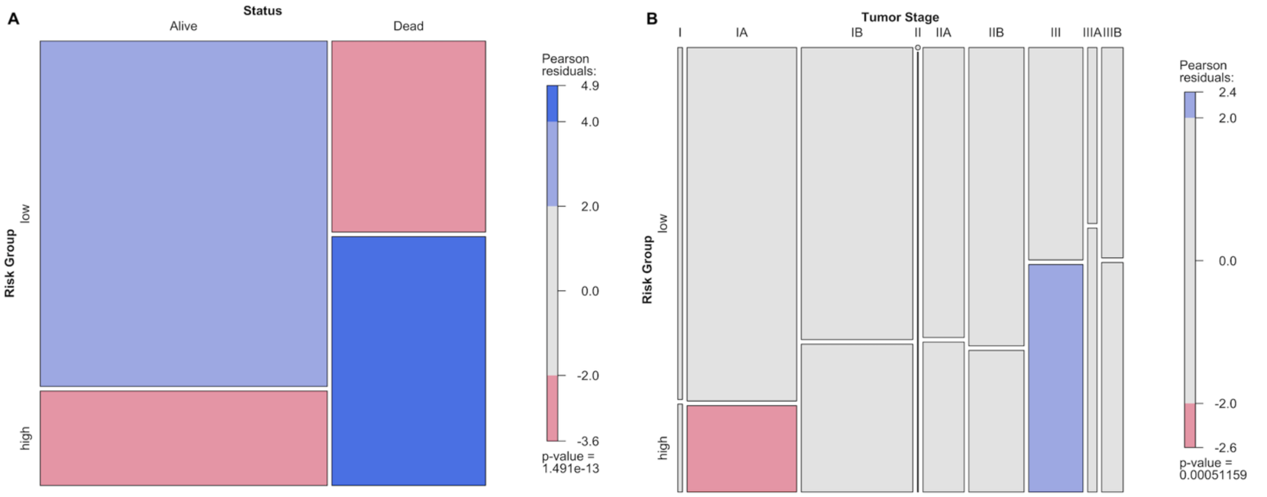


**Figure S3.** Mosaic plots showing association analysis of categorical variables for LUAD training dataset. Pearson residuals show the positive (blue) or negative (red) association between levels of categories. (**A**) The statistical relationships between risk groups and living status of patients; (**B**) The statistical relationships between risk groups and tumor stages of patients.

| Variable | | N | Hazard ratio | p |
|---|---|---|---|---|
| **age** | | 108 | 1.01 (0.94, 1.09) | 0.772 |
| **gender** | female | 65 | Reference | |
| | male | 43 | 5.53 (1.37, 22.42) | 0.017 |
| **tissue_or_organ_of_origin** | Lower lobe, lung | 34 | Reference | |
| | Lung, NOS | 1 | 2.02 (0.07, 58.03) | 0.681 |
| | Main bronchus | 1 | 0.00 (0.00, Inf) | 0.998 |
| | Middle lobe, lung | 5 | 0.15 (0.01, 3.61) | 0.244 |
| | Overlapping lesion of lung | 1 | 0.00 (0.00, Inf) | 0.999 |
| | Upper lobe, lung | 66 | 0.85 (0.25, 2.95) | 0.799 |
| **primary_diagnosis** | Acinar cell carcinoma | 10 | Reference | |
| | Adenocarcinoma with mixed subtypes | 21 | 1.63 (0.09, 28.63) | 0.737 |
| | Adenocarcinoma, NOS | 65 | 1.96 (0.17, 22.05) | 0.585 |
| | Bronchio-alveolar carcinoma, mucinous | 1 | 0.00 (0.00, Inf) | 1.000 |
| | Bronchiolo-alveolar adenocarcinoma, NOS | 1 | 0.00 (0.00, Inf) | 1.000 |
| | Bronchio-alveolar carcinoma, non-mucinous | 5 | 0.00 (0.00, Inf) | 0.999 |
| | Micropapillary carcinoma, NOS | 0 | | |
| | Mucinous adenocarcinoma | 1 | 0.00 (0.00, Inf) | 1.000 |
| | Papillary adenocarcinoma, NOS | 3 | 20.83 (0.34, 1271.95) | 0.148 |
| | Signet ring cell carcinoma | 0 | | |
| | Solid carcinoma, NOS | 1 | 0.00 (0.00, Inf) | 1.000 |
| **ajcc_pathologic_t** | T1 | 18 | Reference | |
| | T1a | 16 | 24.86 (0.34, 1831.08) | 0.143 |
| | T1b | 16 | 7.47 (0.09, 600.86) | 0.369 |
| | T2 | 12 | 2.21 (0.17, 28.93) | 0.546 |
| | T2a | 26 | 4.14 (0.08, 224.14) | 0.486 |
| | T2b | 9 | 2.48 (0.02, 247.73) | 0.699 |
| | T3 | 8 | 0.41 (0.01, 33.34) | 0.693 |
| | T4 | 3 | 0.00 (0.00, Inf) | 1.000 |
| | TX | 0 | | |
| **ajcc_pathologic_n** | N0 | 77 | Reference | |
| | N1 | 17 | 4.58 (0.78, 26.94) | 0.093 |
| | N2 | 12 | 1.13 (0.16, 8.20) | 0.905 |
| | N3 | 1 | 0.00 (0.00, Inf) | 1.000 |
| | NX | 1 | | |
| **ajcc_pathologic_m** | M0 | 66 | Reference | |
| | M1 | 1 | | |
| | M1a | 1 | 0.00 (0.00, Inf) | 1.000 |
| | M1b | 2 | 0.01 (0.00, 1.36) | 0.065 |
| | MX | 38 | 3.15 (0.92, 10.76) | 0.068 |
| **ajcc_staging_system_edition** | 3rd | 1 | Reference | |
| | 4th | 2 | 7.11 (0.04, 1137.62) | 0.449 |
| | 5th | 2 | 14.67 (0.09, 2412.90) | 0.302 |
| | 6th | 21 | 2.62 (0.07, 98.45) | 0.603 |
| | 7th | 82 | 1.08 (0.01, 196.55) | 0.977 |
| **prior_malignancy** | no | 93 | Reference | |
| | yes | 15 | 1.98 (0.41, 9.48) | 0.395 |
| **synchronous_malignancy** | No | 104 | Reference | |
| | Yes | 4 | 0.00 (0.00, Inf) | 0.999 |
| **race** | american indian or alaska native | 0 | Reference | |
| | asian | 1 | 0.00 (0.00, Inf) | 1.000 |
| | black or african american | 13 | 0.43 (0.06, 2.92) | 0.386 |
| | white | 94 | | |
| **ethnicity** | hispanic or latino | 2 | Reference | |
| | not hispanic or latino | 106 | 253872393.00 (0.00, Inf) | 0.999 |
| **years_smoked** | | 108 | 1.02 (0.96, 1.09) | 0.497 |
| **pack_years_smoked** | | 108 | 1.02 (0.99, 1.05) | 0.201 |
| **risk_score** | | 108 | 3.99 (1.72, 9.23) | 0.001 |

**Figure S4.** Multivariate Cox Regression results of clinical variables and risk score in LUAD training dataset. Only risk score has significant result when all clinical variables are included into multivariate analysis.

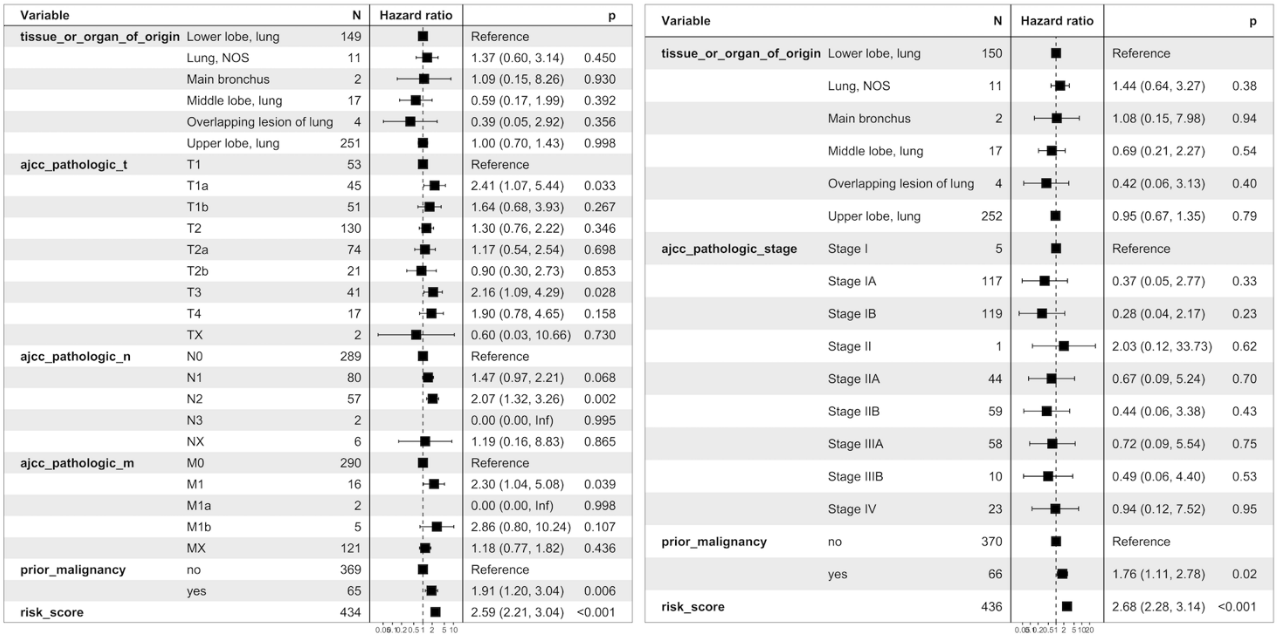

| Variable | | N | Hazard ratio | | p |
|---|---|---|---|---|---|
| tissue_or_organ_of_origin | Lower lobe, lung | 149 | | Reference | |
| | Lung, NOS | 11 | | 1.37 (0.60, 3.14) | 0.450 |
| | Main bronchus | 2 | | 1.09 (0.15, 8.26) | 0.930 |
| | Middle lobe, lung | 17 | | 0.59 (0.17, 1.99) | 0.392 |
| | Overlapping lesion of lung | 4 | | 0.39 (0.05, 2.92) | 0.356 |
| | Upper lobe, lung | 251 | | 1.00 (0.70, 1.43) | 0.998 |
| ajcc_pathologic_t | T1 | 53 | | Reference | |
| | T1a | 45 | | 2.41 (1.07, 5.44) | 0.033 |
| | T1b | 51 | | 1.64 (0.68, 3.93) | 0.267 |
| | T2 | 130 | | 1.30 (0.76, 2.22) | 0.346 |
| | T2a | 74 | | 1.17 (0.54, 2.54) | 0.698 |
| | T2b | 21 | | 0.90 (0.30, 2.73) | 0.853 |
| | T3 | 41 | | 2.16 (1.09, 4.29) | 0.028 |
| | T4 | 17 | | 1.90 (0.78, 4.65) | 0.158 |
| | TX | 2 | | 0.60 (0.03, 10.66) | 0.730 |
| ajcc_pathologic_n | N0 | 289 | | Reference | |
| | N1 | 80 | | 1.47 (0.97, 2.21) | 0.068 |
| | N2 | 57 | | 2.07 (1.32, 3.26) | 0.002 |
| | N3 | 2 | | 0.00 (0.00, Inf) | 0.995 |
| | NX | 6 | | 1.19 (0.16, 8.83) | 0.865 |
| ajcc_pathologic_m | M0 | 290 | | Reference | |
| | M1 | 16 | | 2.30 (1.04, 5.08) | 0.039 |
| | M1a | 2 | | 0.00 (0.00, Inf) | 0.998 |
| | M1b | 5 | | 2.86 (0.80, 10.24) | 0.107 |
| | MX | 121 | | 1.18 (0.77, 1.82) | 0.436 |
| prior_malignancy | no | 369 | | Reference | |
| | yes | 65 | | 1.91 (1.20, 3.04) | 0.006 |
| risk_score | | 434 | | 2.59 (2.21, 3.04) | <0.001 |

| Variable | | N | Hazard ratio | | p |
|---|---|---|---|---|---|
| tissue_or_organ_of_origin | Lower lobe, lung | 150 | | Reference | |
| | Lung, NOS | 11 | | 1.44 (0.64, 3.27) | 0.38 |
| | Main bronchus | 2 | | 1.08 (0.15, 7.98) | 0.94 |
| | Middle lobe, lung | 17 | | 0.69 (0.21, 2.27) | 0.54 |
| | Overlapping lesion of lung | 4 | | 0.42 (0.06, 3.13) | 0.40 |
| | Upper lobe, lung | 252 | | 0.95 (0.67, 1.35) | 0.79 |
| ajcc_pathologic_stage | Stage I | 5 | | Reference | |
| | Stage IA | 117 | | 0.37 (0.05, 2.77) | 0.33 |
| | Stage IB | 119 | | 0.28 (0.04, 2.17) | 0.23 |
| | Stage II | 1 | | 2.03 (0.12, 33.73) | 0.62 |
| | Stage IIA | 44 | | 0.67 (0.09, 5.24) | 0.70 |
| | Stage IIB | 59 | | 0.44 (0.06, 3.38) | 0.43 |
| | Stage IIIA | 58 | | 0.72 (0.09, 5.54) | 0.75 |
| | Stage IIIB | 10 | | 0.49 (0.06, 4.40) | 0.53 |
| | Stage IV | 23 | | 0.94 (0.12, 7.52) | 0.95 |
| prior_malignancy | no | 370 | | Reference | |
| | yes | 66 | | 1.76 (1.11, 2.78) | 0.02 |
| risk_score | | 436 | | 2.68 (2.28, 3.14) | <0.001 |

**Figure S5.** Multivariate Cox Regression results of selected clinical variables (which have significant results in univariate cox analysis) and risk score in LUAD training dataset. Risk score, t, n, m stages and history of prior malignancy have significant effects on survival. When pathologic tumor stage is used instead of t, n, m stages, only risk score and history of prior malignancy show significant effect on survival.

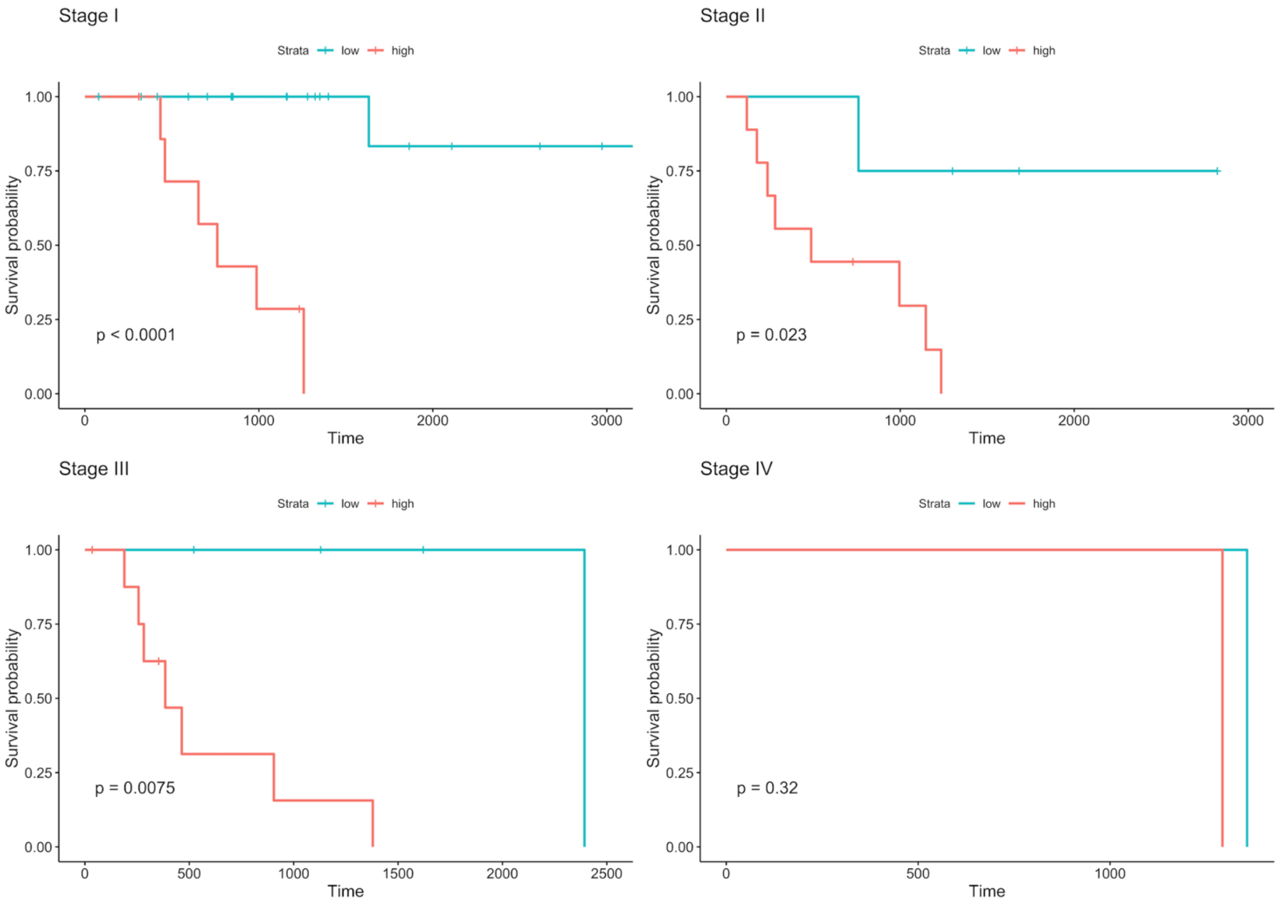


**Figure S6.** Survival analysis of risk groups clustered by using signature gene expression at different tumor stages in LUAD test dataset.

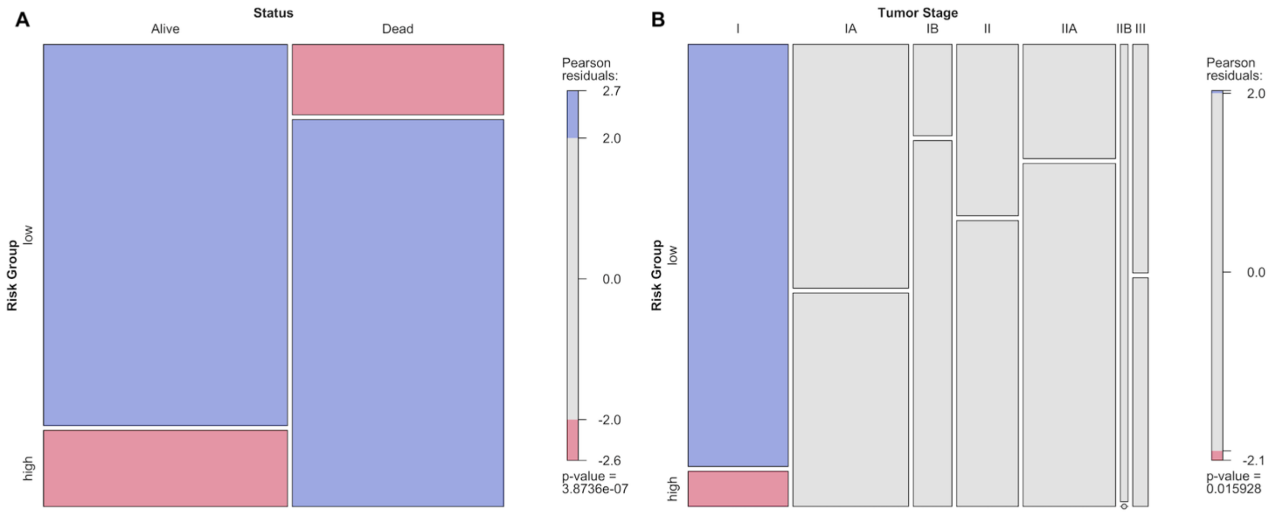


**Figure S7.** Mosaic plots showing association analysis of categorical variables for LUAD test dataset. Pearson residuals show the positive (blue) or negative (red) association between levels of categories. (**A**) The statistical relationships between risk groups and living status of patients; (**B**) The statistical relationships between risk groups and tumor stages of patients.

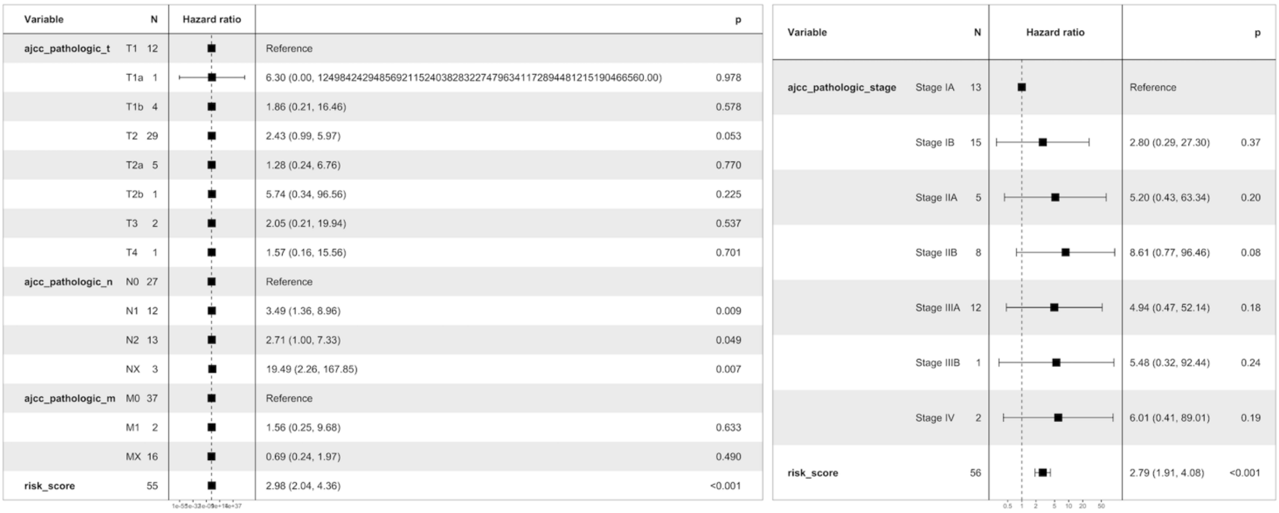


**Figure S8.** Multivariate Cox Regression results of selected clinical variables (which have significant results in univariate cox analysis) and risk score in LUAD test dataset. Risk score and n stages have significant effect on survival. When pathologic tumor stage is used instead of t, n, m stages, only risk score shows significant effect on survival.

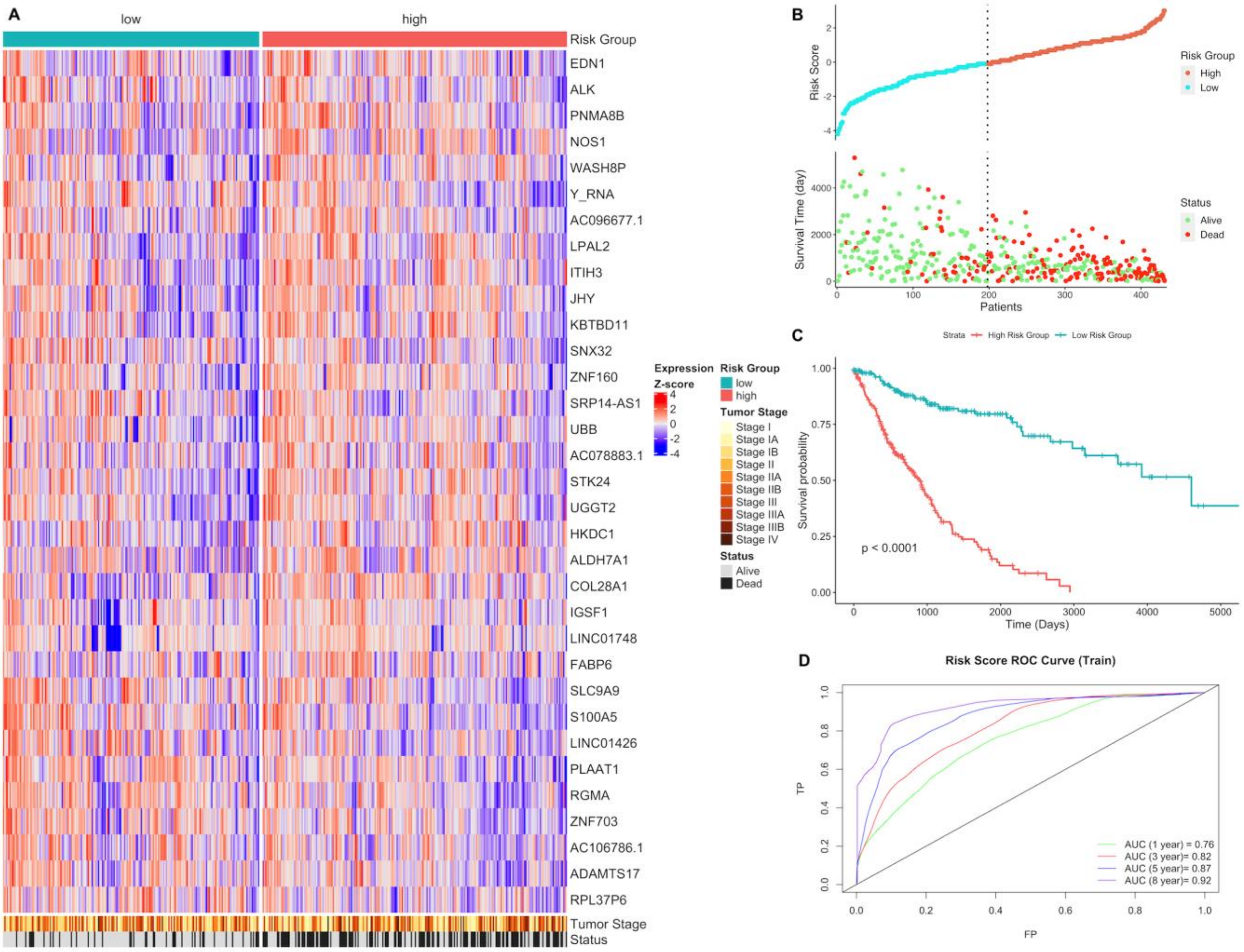


**Figure S9.** Gene expression signature and risk clustering of LUSC training dataset. (**A**) Expression heatmap of the signature genes in tumor samples of LUSC patients in train dataset. Train dataset patients were clustered into high-risk and low-risk groups; (**B**) Scatter plot showing risk scores, survival time and separation point of the patients into risk groups. (**C**) KM survival plot showing the overall survival probability between risk groups. (**D**) ROC curve showing prediction power of risk score in train dataset for 1, 3, 5 and 8 years.

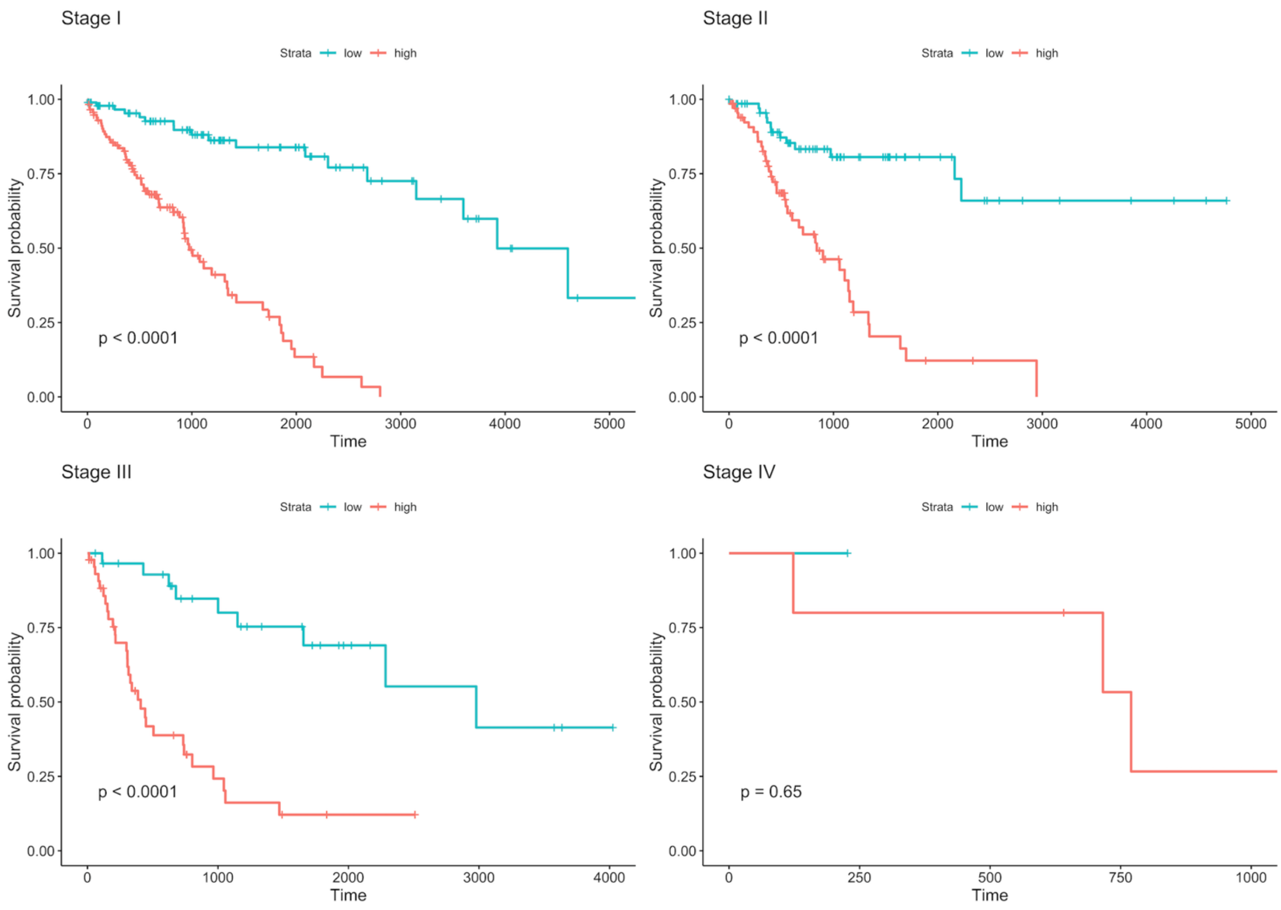


**Figure S10.** Survival analysis of risk groups clustered by using signature gene expression at different tumor stages in LUSC training dataset.

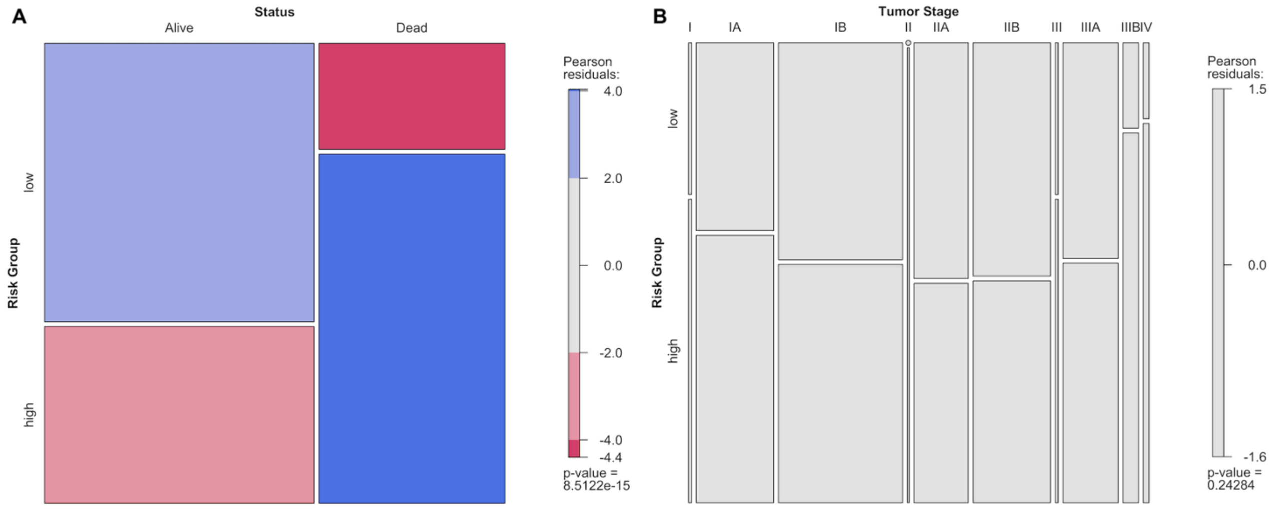


**Figure S11.** Mosaic plots showing association analysis of categorical variables for LUSC training dataset. Pearson residuals show the positive (blue) or negative (red) association between levels of categories. (**A**) The statistical relationships between risk groups and living status of patients; (**B**) The statistical relationships between risk groups and tumor stages of patients.

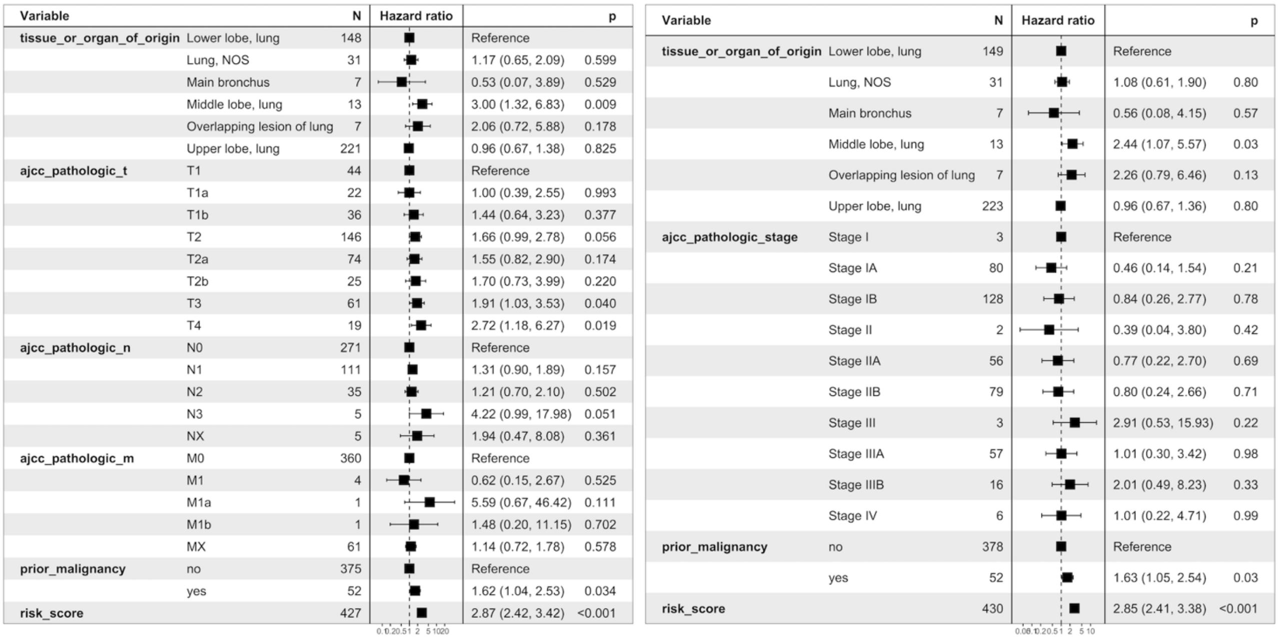

| Variable | | N | Hazard ratio | | p |
|---|---|---|---|---|---|
| tissue_or_organ_of_origin | Lower lobe, lung | 148 | | Reference | |
| | Lung, NOS | 31 | | 1.17 (0.65, 2.09) | 0.599 |
| | Main bronchus | 7 | | 0.53 (0.07, 3.89) | 0.529 |
| | Middle lobe, lung | 13 | | 3.00 (1.32, 6.83) | 0.009 |
| | Overlapping lesion of lung | 7 | | 2.06 (0.72, 5.88) | 0.178 |
| | Upper lobe, lung | 221 | | 0.96 (0.67, 1.38) | 0.825 |
| ajcc_pathologic_t | T1 | 44 | | Reference | |
| | T1a | 22 | | 1.00 (0.39, 2.55) | 0.993 |
| | T1b | 36 | | 1.44 (0.64, 3.23) | 0.377 |
| | T2 | 146 | | 1.66 (0.99, 2.78) | 0.056 |
| | T2a | 74 | | 1.55 (0.82, 2.90) | 0.174 |
| | T2b | 25 | | 1.70 (0.73, 3.99) | 0.220 |
| | T3 | 61 | | 1.91 (1.03, 3.53) | 0.040 |
| | T4 | 19 | | 2.72 (1.18, 6.27) | 0.019 |
| ajcc_pathologic_n | N0 | 271 | | Reference | |
| | N1 | 111 | | 1.31 (0.90, 1.89) | 0.157 |
| | N2 | 35 | | 1.21 (0.70, 2.10) | 0.502 |
| | N3 | 5 | | 4.22 (0.99, 17.98) | 0.051 |
| | NX | 5 | | 1.94 (0.47, 8.08) | 0.361 |
| ajcc_pathologic_m | M0 | 360 | | Reference | |
| | M1 | 4 | | 0.62 (0.15, 2.67) | 0.525 |
| | M1a | 1 | | 5.59 (0.67, 46.42) | 0.111 |
| | M1b | 1 | | 1.48 (0.20, 11.15) | 0.702 |
| | MX | 61 | | 1.14 (0.72, 1.78) | 0.578 |
| prior_malignancy | no | 375 | | Reference | |
| | yes | 52 | | 1.62 (1.04, 2.53) | 0.034 |
| risk_score | | 427 | | 2.87 (2.42, 3.42) | <0.001 |

| Variable | | N | Hazard ratio | | p |
|---|---|---|---|---|---|
| tissue_or_organ_of_origin | Lower lobe, lung | 149 | | Reference | |
| | Lung, NOS | 31 | | 1.08 (0.61, 1.90) | 0.80 |
| | Main bronchus | 7 | | 0.56 (0.08, 4.15) | 0.57 |
| | Middle lobe, lung | 13 | | 2.44 (1.07, 5.57) | 0.03 |
| | Overlapping lesion of lung | 7 | | 2.26 (0.79, 6.46) | 0.13 |
| | Upper lobe, lung | 223 | | 0.96 (0.67, 1.36) | 0.80 |
| ajcc_pathologic_stage | Stage I | 3 | | Reference | |
| | Stage IA | 80 | | 0.46 (0.14, 1.54) | 0.21 |
| | Stage IB | 128 | | 0.84 (0.26, 2.77) | 0.78 |
| | Stage II | 2 | | 0.39 (0.04, 3.80) | 0.42 |
| | Stage IIA | 56 | | 0.77 (0.22, 2.70) | 0.69 |
| | Stage IIB | 79 | | 0.80 (0.24, 2.66) | 0.71 |
| | Stage III | 3 | | 2.91 (0.53, 15.93) | 0.22 |
| | Stage IIIA | 57 | | 1.01 (0.30, 3.42) | 0.98 |
| | Stage IIIB | 16 | | 2.01 (0.49, 8.23) | 0.33 |
| | Stage IV | 6 | | 1.01 (0.22, 4.71) | 0.99 |
| prior_malignancy | no | 378 | | Reference | |
| | yes | 52 | | 1.63 (1.05, 2.54) | 0.03 |
| risk_score | | 430 | | 2.85 (2.41, 3.38) | <0.001 |

**Figure S12.** Multivariate Cox Regression results of selected clinical variables (which have significant results in univariate cox analysis) and risk score in LUSC training dataset. Risk score, tissue or organ of origin, t and n stages and history of prior malignancy have significant effects on survival. When pathologic tumor stage is used instead of t, n, m stages, tissue or organ of origin, risk score and history of prior malignancy show significant effect on survival.

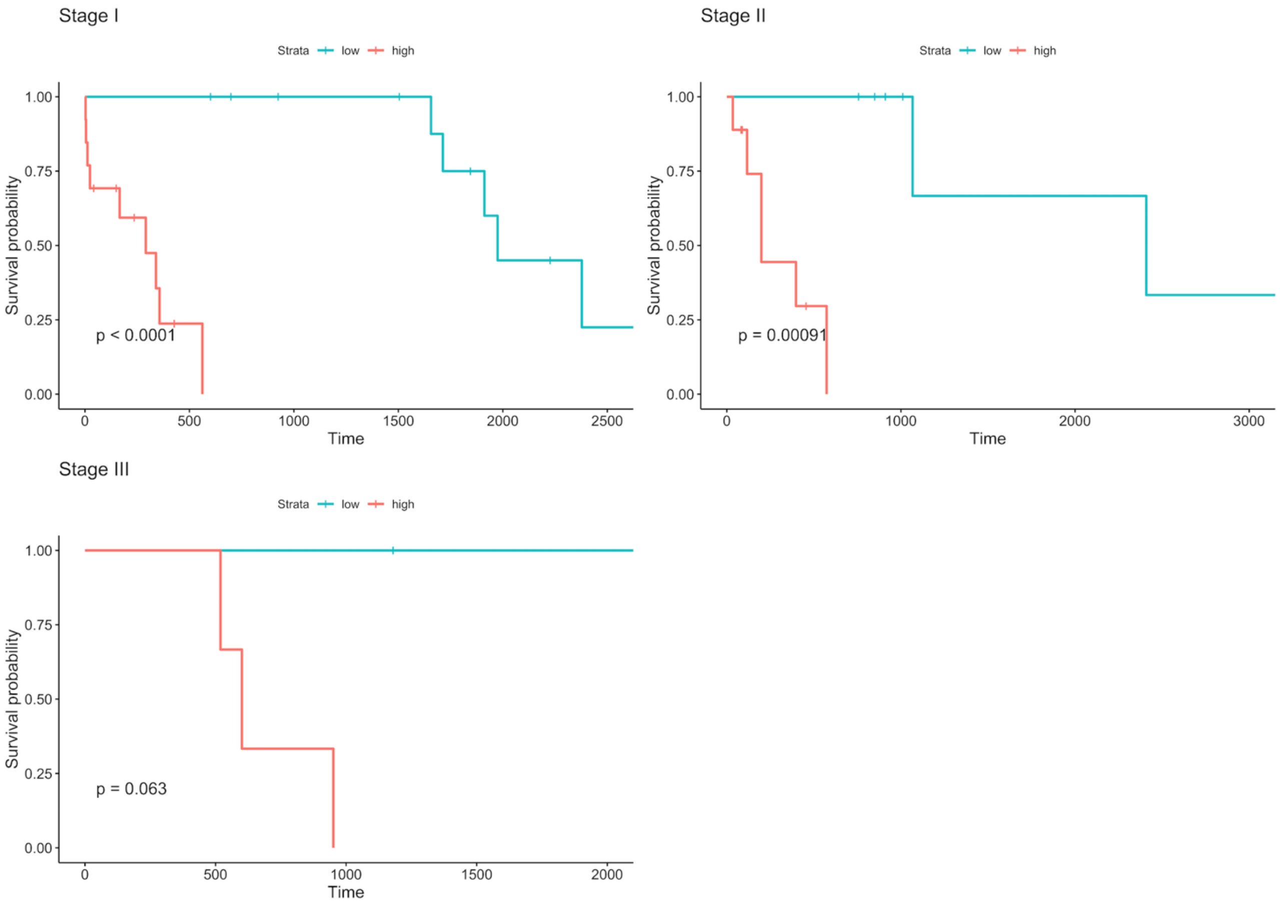


**Figure S13.** Survival analysis of risk groups clustered by using signature gene expression at different tumor stages in LUSC test dataset.

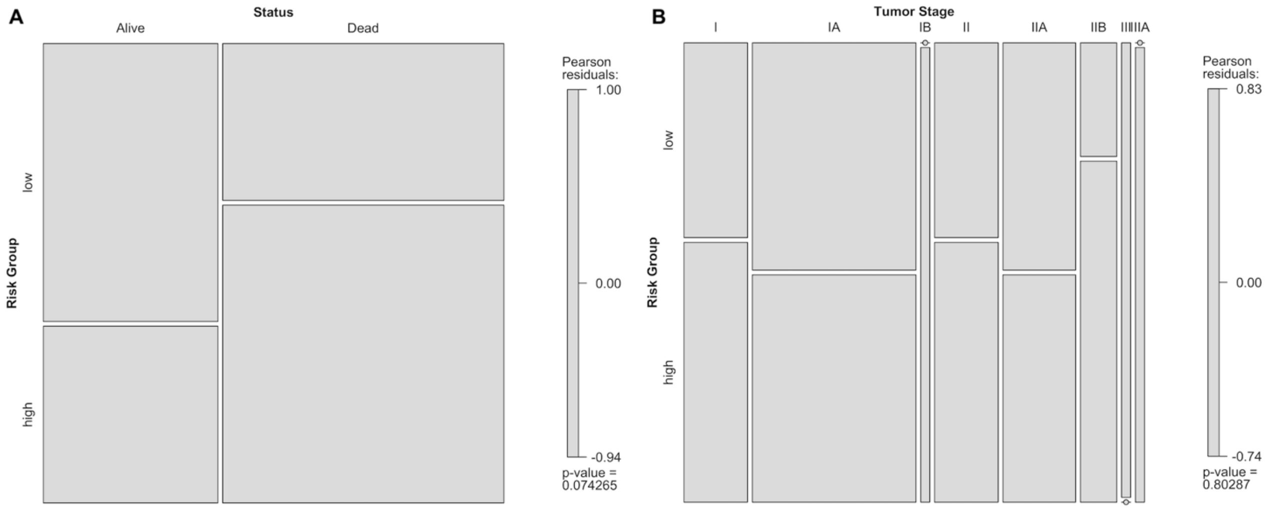


**Figure S14.** Mosaic plots showing association analysis of categorical variables for LUSC test dataset. Pearson residuals show the positive (blue) or negative (red) association between levels of categories. (**A**) The statistical relationships between risk groups and living status of patients; (**B**) The statistical relationships between risk groups and tumor stages of patients.

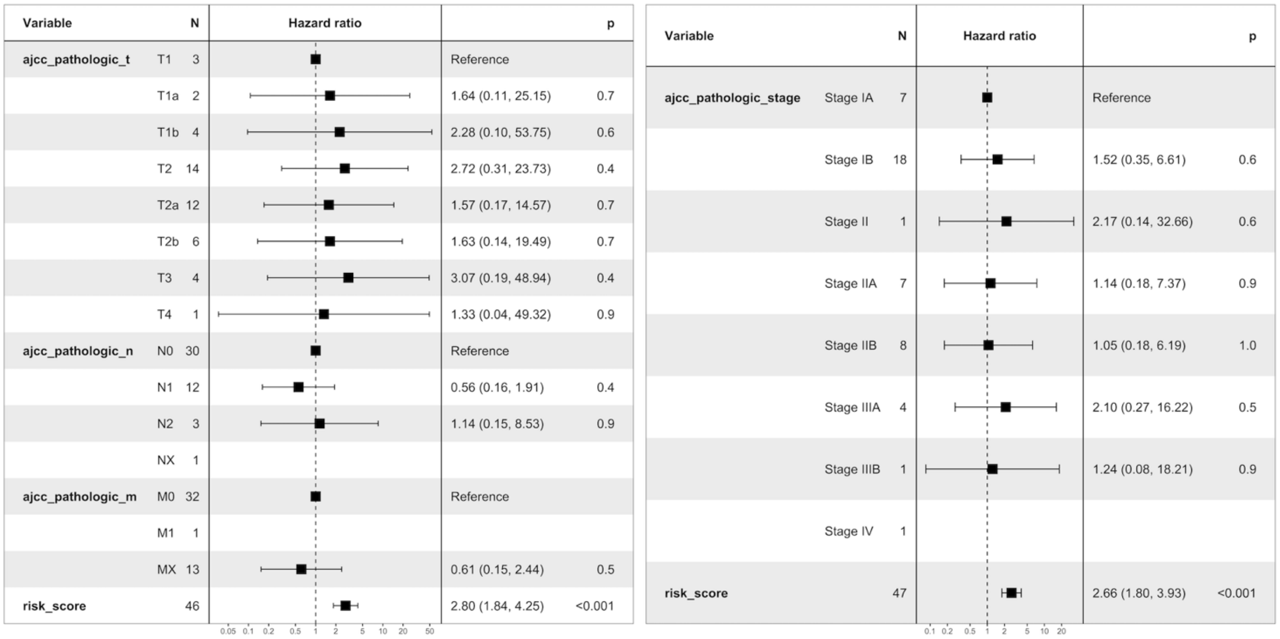


**Figure S15.** Multivariate Cox Regression results of selected clinical variables (which have significant results in univariate cox analysis) and risk score in LUSC test dataset. Only risk score has significant effect on survival either t, n, m stages or pathologic tumor stage is used instead of t, n, m stages.

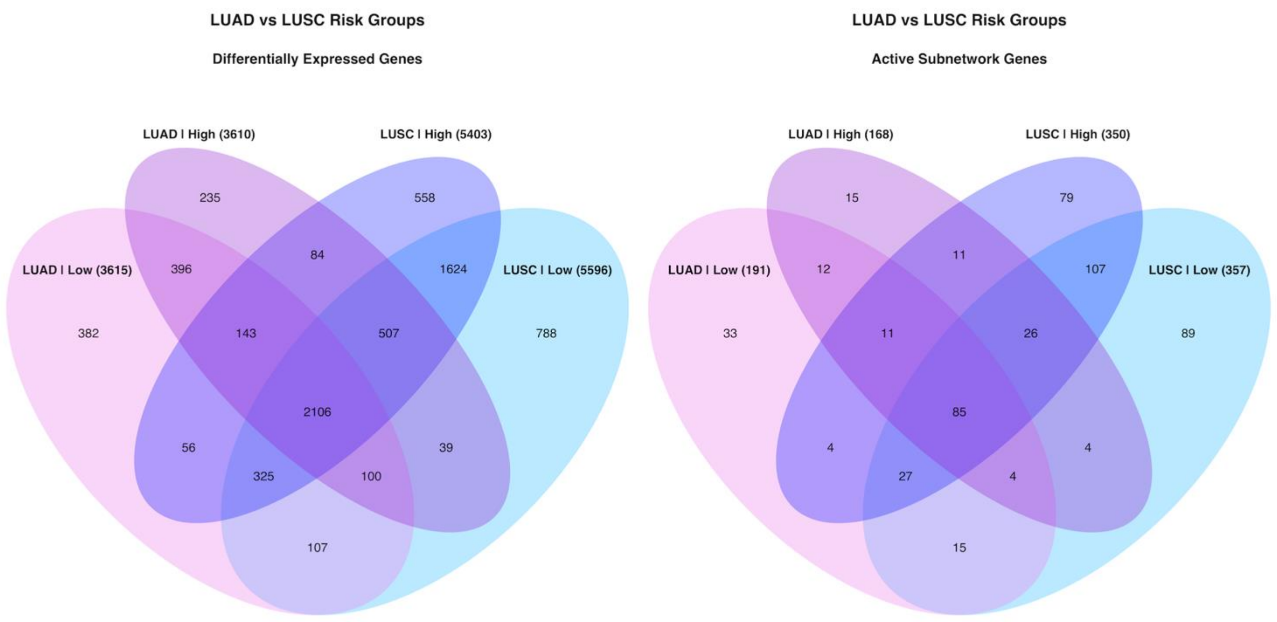


**Figure S16.** Venn diagram of differentially expressed genes in tumor samples of risk groups for LUAD and LUSC test groups.

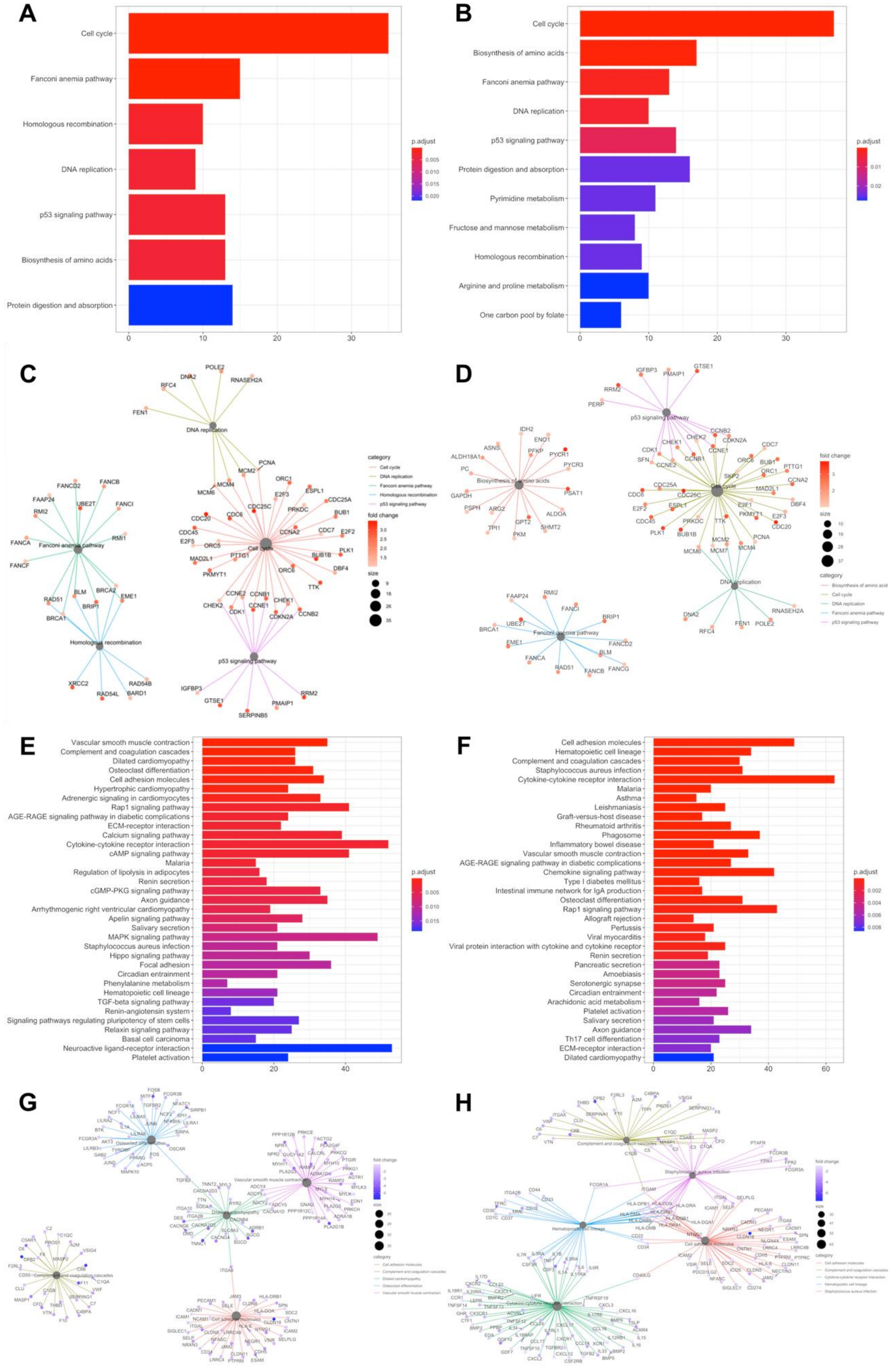


**Figure S17.** Pathway enrichment of DEGs of LUAD risk groups. **A**, **B**, **C**, **D**: Upregulated pathways; **E**, **F**, **G**, **H**: Downregulated pathways. **A**, **C**, **E**, **G**: LUAD low-risk group; **B**, **D**, **F**, **H**: LUAD high-risk group.

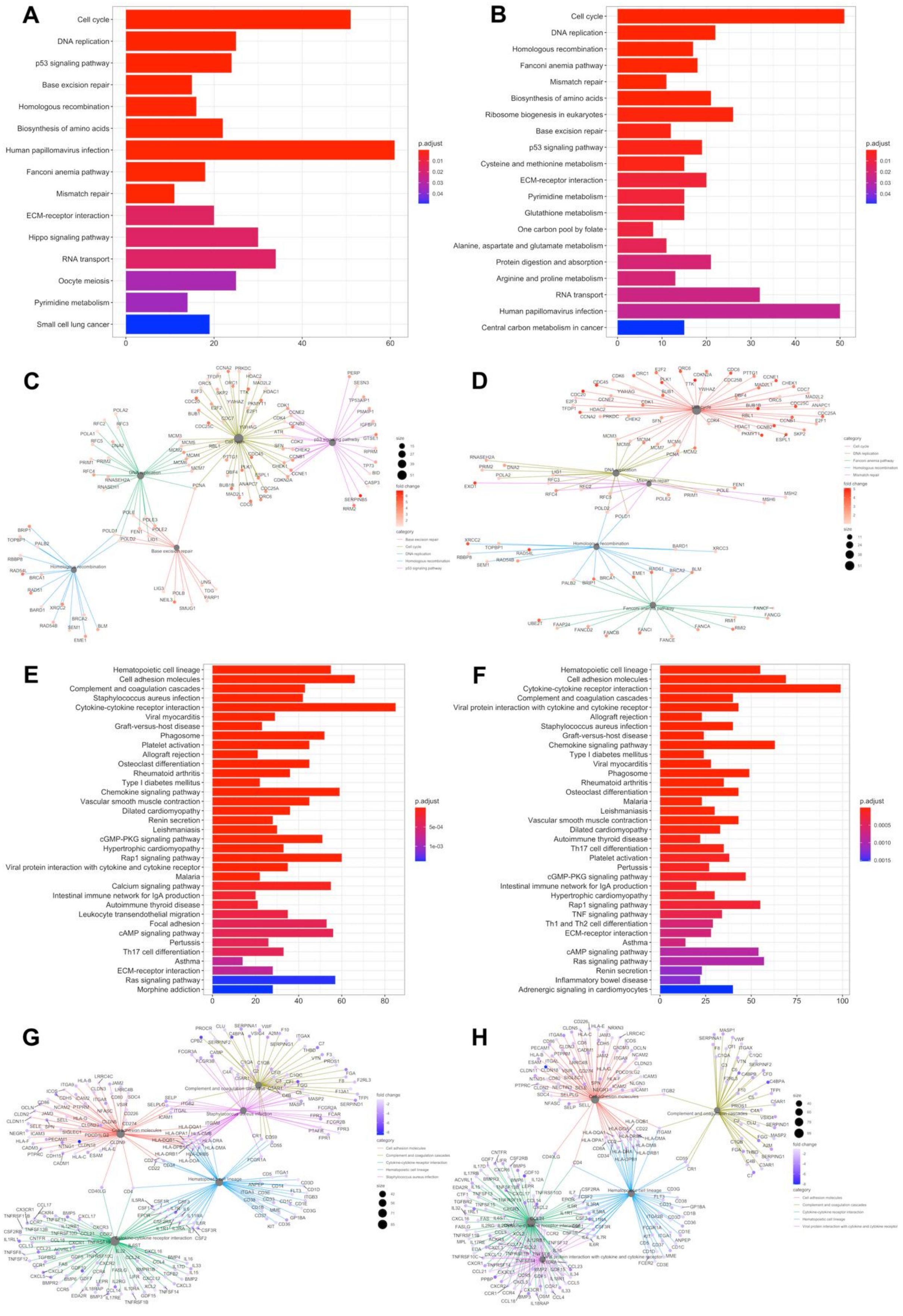


**Figure S18.** Pathway enrichment of DEGs of LUSC risk groups. **A**, **B**, **C**, **D**: Upregulated pathways; **E**, **F**, **G**, **H**: Downregulated pathways. **A**, **C**, **E**, **G**: LUSC low-risk group; **B**, **D**, **F**, **H**: LUSC high-risk group.

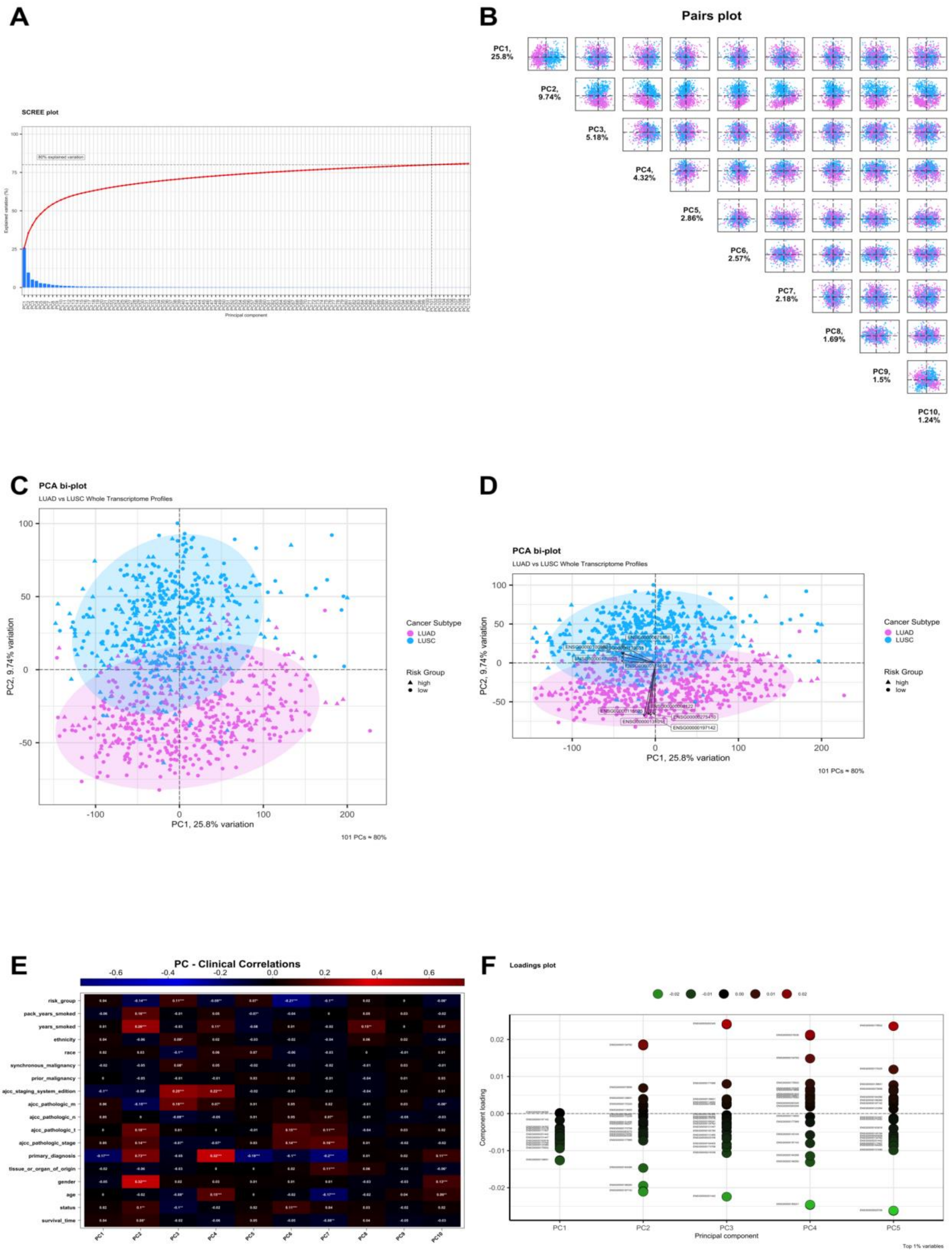


**Figure S19.** Principal Component Analysis (PCA) of whole transcriptome profiles of all patients in LUAD and LUSC projects. (**A**) Scree plot showing variances of the PCs. (**B**) Pairs plot of first 10 PCs (**C**) PCA biplot of PC1 and PC2 explaining 35.4% of variances among LUAD and LUSC samples. (**D**) PCA biplot of PC1 and PC2 with loadings (**E**) Clinical correlations of first 10 PCs (**F**) Loading plot of first 10 PCs.

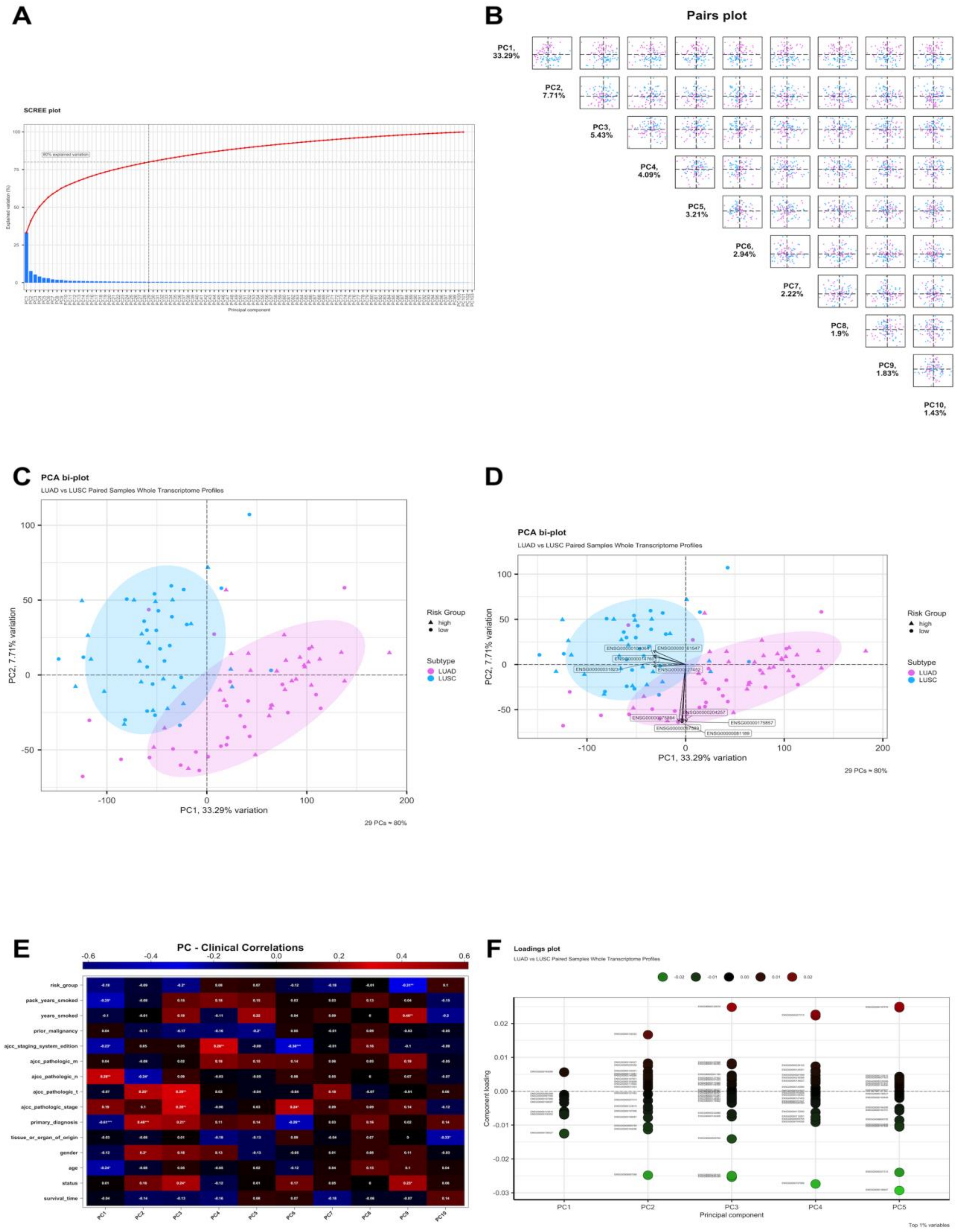


**Figure S20.** Principal Component Analysis (PCA) of whole transcriptome profiles of patients in LUAD and LUSC test groups. (**A**) Scree plot showing variances of the PCs. (**B**) Pairs plot of first 10 PCs (**C**) PCA biplot of PC1 and PC2 explaining 35.4% of variances among LUAD and LUSC samples. (**D**) PCA biplot of PC1 and PC2 with loadings (**E**) Clinical correlations of first 10 PCs (**F**) Loading plot of first 10 PCs.

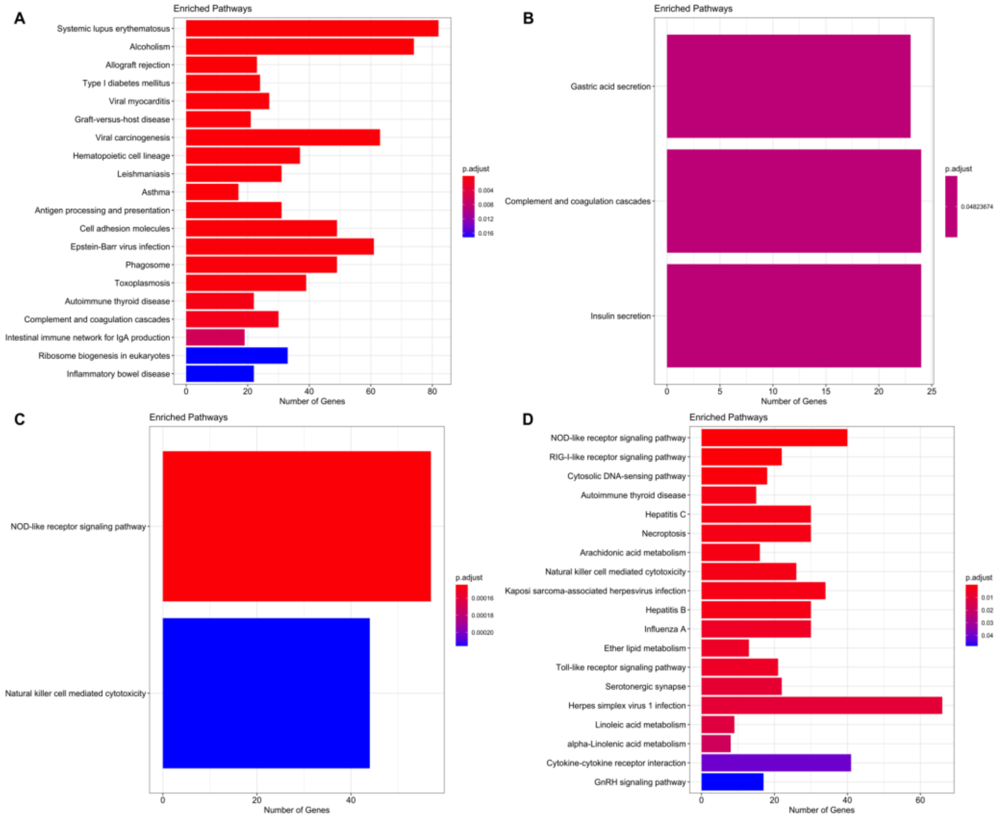


**Figure S21.** Pathway enrichment of CNV genes of LUAD risk groups. **A**, **B**: Pathways of amplified genes in tumor samples; **C**, **D**: Pathways of deleted genes in tumor samples. **A**, **C**: LUAD low-risk group; **B**, **D**: LUAD high-risk group.

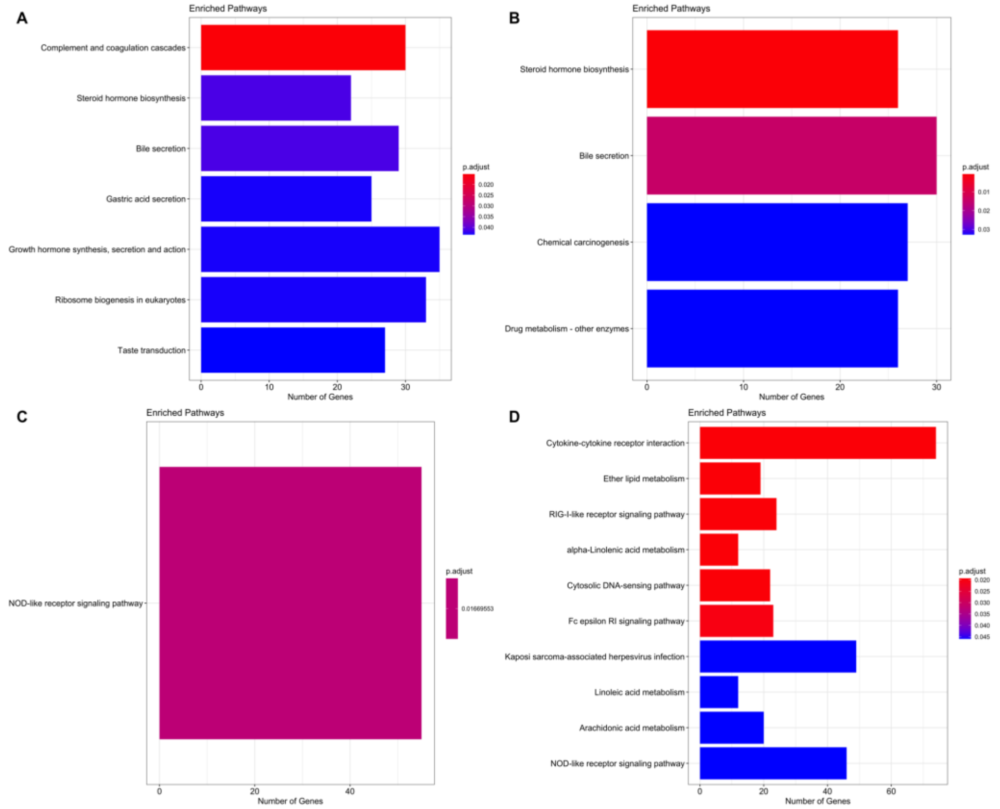


**Figure S22.** Pathway enrichment of CNV genes of LUSC risk groups. **A**, **B**: Pathways of amplified genes in tumor samples; **C**, **D**: Pathways of deleted genes in tumor samples. **A**, **C**: LUSC low-risk group; **B**, **D**: LUSC high-risk group.

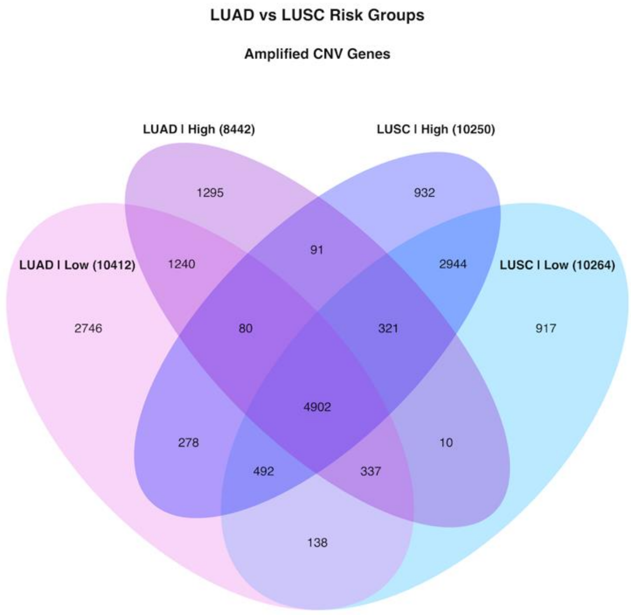


**Figure S23.** Venn diagram of genes which have significant copy number alterations in tumor samples of LUAD and LUSC risk groups.

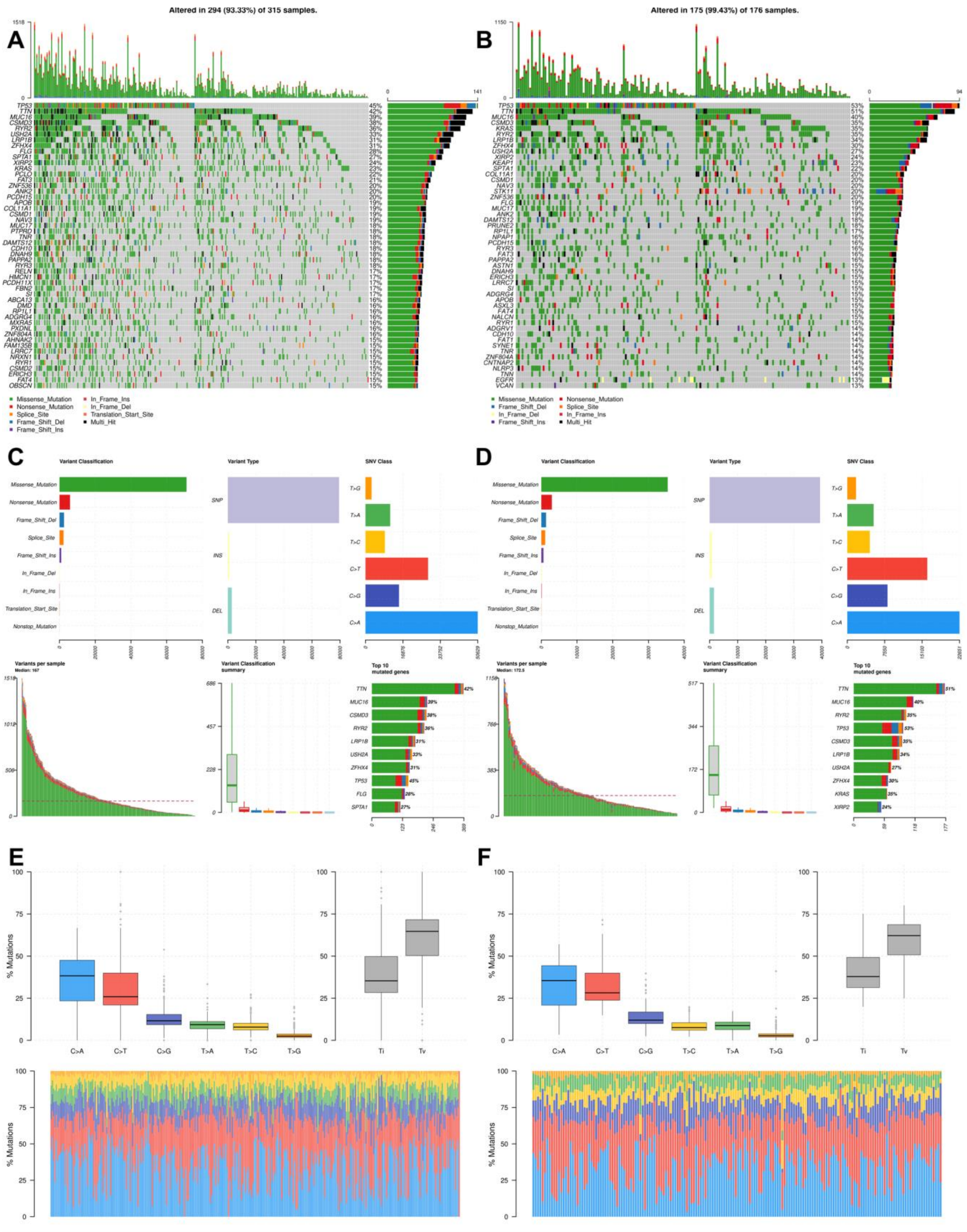


**Figure S24.** Summary of SNVs in LUAD risk groups. **A**, **B**: Oncoplot of top frequent genes in tumor samples of LUAD risk groups; **C**, **D**: Summary plot of SNVs in risk groups. **E, F**: Base substitutions and frequencies of SNVs in risk groups. **A**, **C, E**: LUAD low-risk group; **B**, **D, F**: LUAD high-risk group.

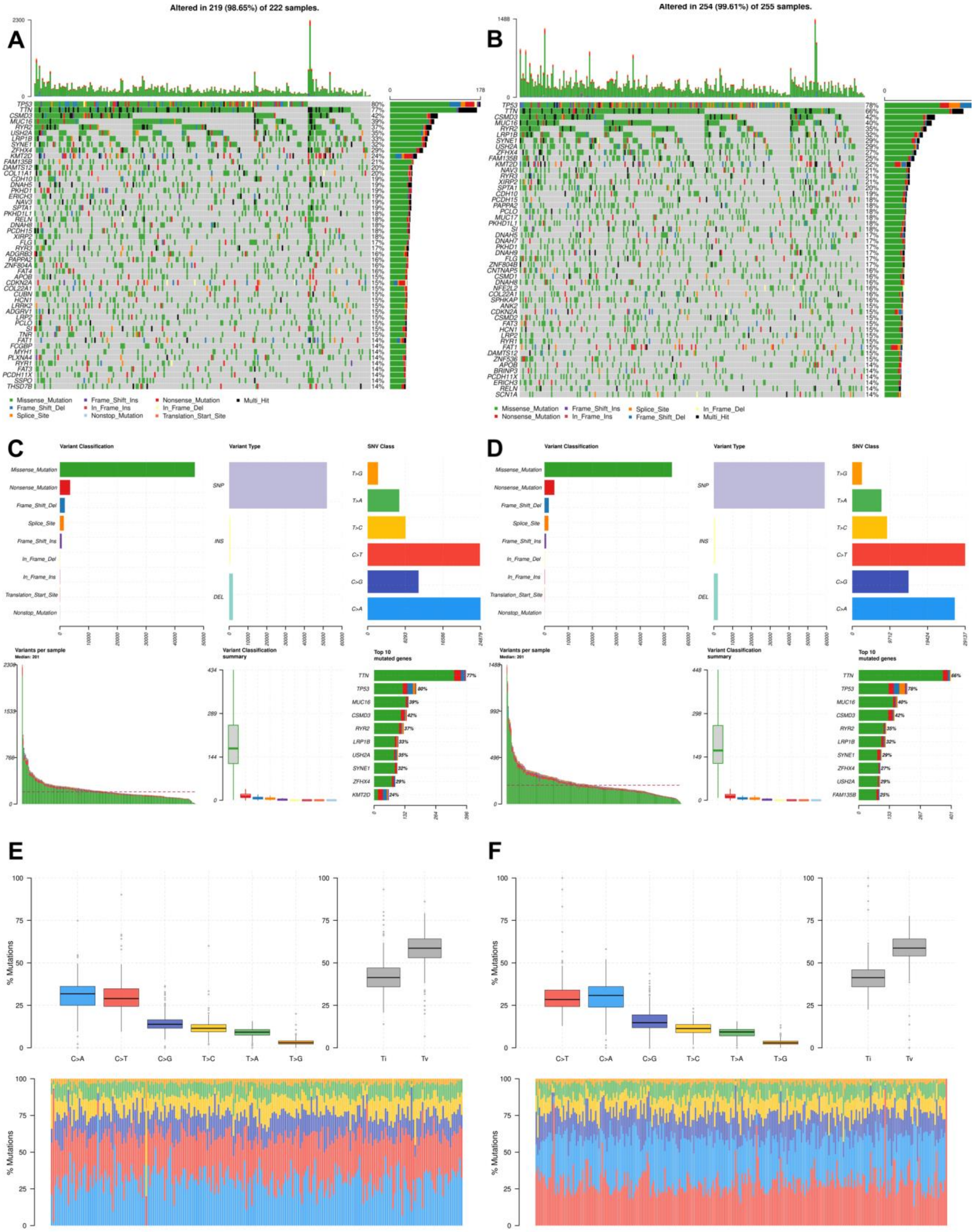


**Figure S25.** Summary of SNVs in LUSC risk groups. **A**, **B**: Oncoplot of top frequent genes in tumor samples of LUSC risk groups; **C**, **D**: Summary plot of SNVs in risk groups. **E, F**: Base substitutions and frequencies of SNVs in risk groups. **A**, **C**, **E**: LUSClow-risk group; **B**, **D, F**: LUSC high-risk group.

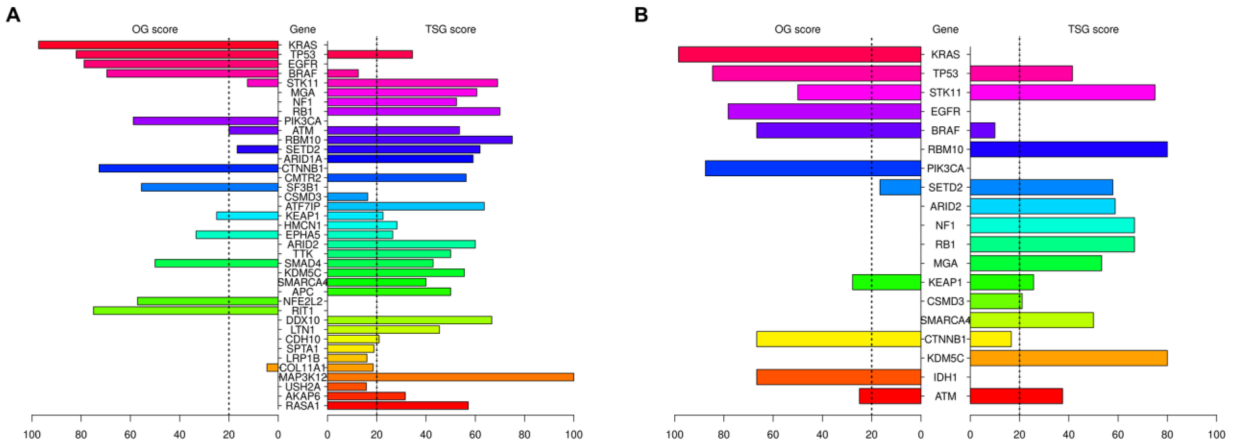


**Figure S26.** SomInaClust result of potential driver genes containing significant SNVs in LUAD risk groups. SomInaClust calculates oncogene (OG) score and tumor suppressor gene (TSG) score for each significant gene and classifies the gene according to the score threshold (20) and reference database. (**A**) Scores of driver genes in LUAD low-risk group. (**B**) Scores of driver genes in LUAD high-risk group.

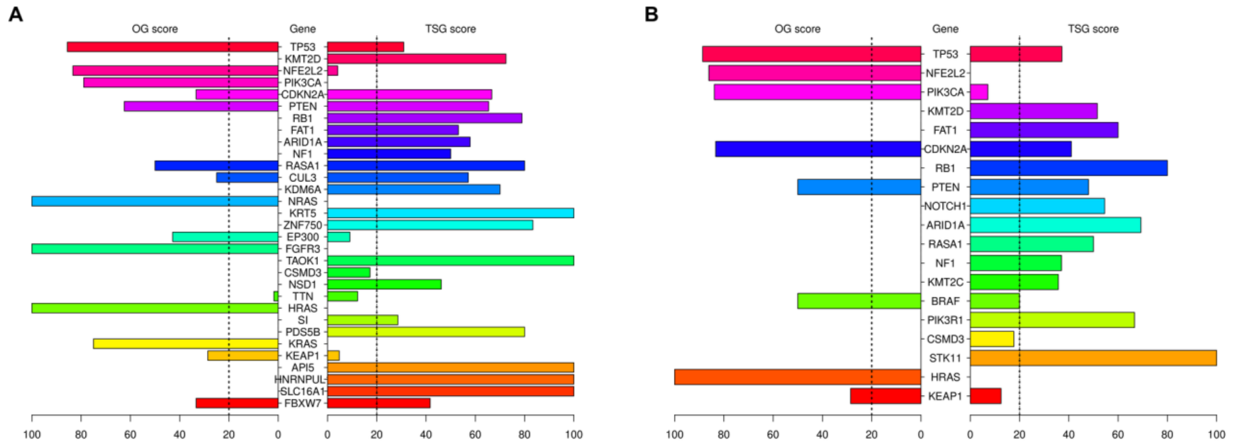


**Figure S27.** SomInaClust result of potential driver genes containing significant SNVs in LUSC risk groups. SomInaClust calculates oncogene (OG) score and tumor suppressor gene (TSG) score for each significant gene and classifies the gene according to the score threshold (20) and reference database. (**A**) Scores of driver genes in LUSC low-risk group. (**B**) Scores of driver genes in LUSC high-risk group.

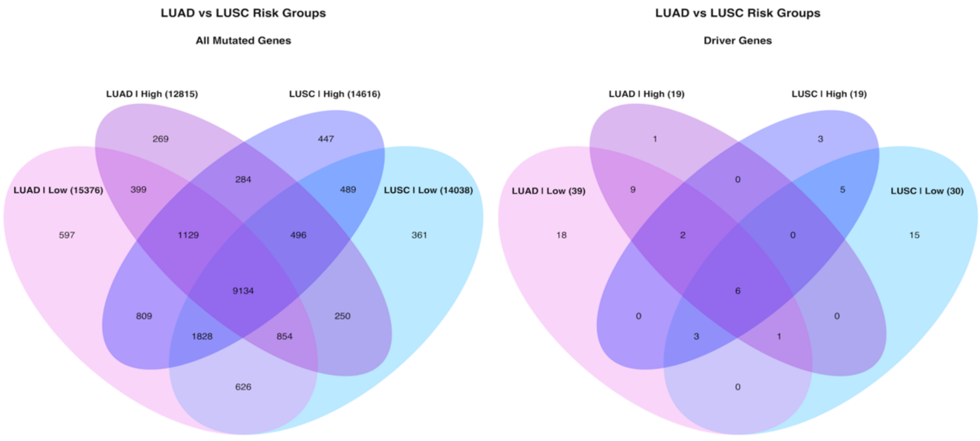


**Figure S28.** Venn diagram of all genes and potential driver genes containing SNVs of LUAD and LUSC risk groups.

**Table S1.** Gene list of expression signature in LUAD. Ensemble Gene IDs were used in signature analysis and then enriched by using BioMart database.

| Ensemble Gene ID | Entrez Gene ID | Gene Name | Description |
|---|---|---|---|
| ENSG00000035720 | 26228 | STAP1 | Signal transducing adaptor family member 1 |
| ENSG00000100522 | 64841 | GNPNAT1 | Glucosamine-phosphate N-acetyltransferase 1 |
| ENSG00000100628 | 51676 | ASB2 | Ankyrin repeat and SOCS box containing 2 |
| ENSG00000107984 | 22943 | DKK1 | Dickkopf WNT signaling pathway inhibitor 1 |
| ENSG00000112796 | 59084 | ENPP5 | Ectonucleotide pyrophosphatase/phosphodiesterase family member 5 |
| ENSG00000115009 | 6364 | CCL20 | C-C motif chemokine ligand 20 |
| ENSG00000117477 | 57821 | CCDC181 | Coiled-coil domain containing 181 |
| ENSG00000119729 | 23433 | RHOQ | Ras homolog family member Q |
| ENSG00000121898 | 119587 | CPXM2 | Carboxypeptidase X, M14 family member 2 |
| ENSG00000123570 | 51209 | RAB9B | RAB9B, member RAS oncogene family |
| ENSG00000127903 | 90485 | ZNF835 | Zinc finger protein 835 |
| ENSG00000132517 | 55065 | SLC52A1 | Solute carrier family 52 member 1 |
| ENSG00000134333 | 3939 | LDHA | Lactate dehydrogenase A |
| ENSG00000143819 | 2052 | EPHX1 | Epoxide hydrolase 1 |
| ENSG00000147509 | 8601 | RGS20 | Regulator of G protein signaling 20 |
| ENSG00000156738 | 931 | MS4A1 | Membrane spanning 4-domains A1 |
| ENSG00000163606 | 131450 | CD200R1 | CD200 receptor 1 |
| ENSG00000164418 | 2898 | GRIK2 | Glutamate ionotropic receptor kainate type subunit 2 |
| ENSG00000166106 | 170689 | ADAMTS15 | ADAM metallopeptidase with thrombospondin type 1 motif 15 |
| ENSG00000167772 | 51129 | ANGPTL4 | Angiopoietin like 4 |
| ENSG00000170561 | 153572 | IRX2 | Iroquois homeobox 2 |
| ENSG00000179241 | 143458 | LDLRAD3 | Low density lipoprotein receptor class A domain containing 3 |
| ENSG00000182057 | - | OGFRP1 | Opioid growth factor receptor pseudogene 1 |
| ENSG00000183066 | 164684 | WBP2NL | WBP2 N-terminal like |
| ENSG00000183734 | 430 | ASCL2 | Achaete-scute family bHLH transcription factor 2 |
| ENSG00000196781 | 7088 | TLE1 | TLE family member 1, transcriptional corepressor |
| ENSG00000197124 | 91120 | ZNF682 | Zinc finger protein 682 |
| ENSG00000203943 | 148418 | SAMD13 | Sterile alpha motif domain containing 13 |
| ENSG00000224429 | - | LINC00539 | Long intergenic non-protein coding RNA 539 |
| ENSG00000228221 | 100505566 | LINC00578 | Long intergenic non-protein coding RNA 578 |
| ENSG00000230882 | - | AC005077.4 | Hypothetical protein LOC285908 (LOC285908) pseudogene |
| ENSG00000254893 | - | AC113404.3 | RAP1B, member of RAS oncogene family pseudogene |
| ENSG00000267470 | 100507433 | ZNF571-AS1 | ZNF571 antisense RNA 1 |
| ENSG00000272068 | - | AL365181.2 | Novel transcript |
| ENSG00000272462 | - | U91328.1 | Novel transcript |

**Table S2.** KEGG pathway enrichment of expression signature gene list in LUAD by using KEGG Mapper tool

| Pathway ID | Description | Gene ID | Count |
|---|---|---|---|
| hsa01100 | Metabolic pathways | GNPNAT1, LDHA | 2 |
| hsa05230 | Central carbon metabolism in cancer | LDHA | 1 |
| hsa00010 | Glycolysis / Gluconeogenesis | LDHA | 1 |
| hsa04922 | Glucagon signaling pathway | LDHA | 1 |
| hsa04066 | HIF-1 signaling pathway | LDHA | 1 |
| hsa00620 | Pyruvate metabolism | LDHA | 1 |
| hsa00270 | Cysteine and methionine metabolism | LDHA | 1 |
| hsa00640 | Propanoate metabolism | LDHA | 1 |
| hsa03320 | PPAR signaling pathway | ANGPTL4 | 1 |
| hsa04979 | Cholesterol metabolism | ANGPTL4 | 1 |
| hsa00520 | Amino sugar and nucleotide sugar metabolism | GNPNAT1 | 1 |
| hsa04910 | Insulin signaling pathway | RHOQ | 1 |
| hsa04640 | Hematopoietic cell lineage | MS4A1 | 1 |
| hsa05204 | Chemical carcinogenesis | EPHX1 | 1 |
| hsa00980 | Metabolism of xenobiotics by cytochrome P450 | EPHX1 | 1 |
| hsa04976 | Bile secretion | EPHX1 | 1 |
| hsa04668 | TNF signaling pathway | CCL20 | 1 |
| hsa04657 | IL-17 signaling pathway | CCL20 | 1 |
| hsa04060 | Cytokine-cytokine receptor interaction | CCL20 | 1 |
| hsa05323 | Rheumatoid arthritis | CCL20 | 1 |
| hsa04061 | Viral protein interaction with cytokine and cytokine receptor | CCL20 | 1 |
| hsa04062 | Chemokine signaling pathway | CCL20 | 1 |
| hsa05132 | Salmonella infection | RAB9B | 1 |
| hsa05162 | Measles | RAB9B | 1 |
| hsa05168 | Herpes simplex virus 1 infection | ZNF682 | 1 |
| hsa05167 | Kaposi sarcoma-associated herpesvirus infection | CD200R1 | 1 |
| hsa05022 | Pathways of neurodegeneration - multiple diseases | DKK1 | 1 |
| hsa05010 | Alzheimer disease | DKK1 | 1 |
| hsa04310 | Wnt signaling pathway | DKK1 | 1 |
| hsa04080 | Neuroactive ligand-receptor interaction | GRIK2 | 1 |
| hsa04724 | Glutamatergic synapse | GRIK2 | 1 |

**Table S3.** Gene list of expression signature in LUSC. Ensemble Gene IDs were used in signature analysis and then enriched by using BioMart database.

| Ensemble Gene ID | Entrez Gene ID | Gene Name | Description |
|---|---|---|---|
| ENSG00000078401 | 1906 | EDN1 | Endothelin 1 |
| ENSG00000089250 | 4842 | NOS1 | Nitric oxide synthase 1 |
| ENSG00000102572 | 8428 | STK24 | Serine/threonine kinase 24 |
| ENSG00000102595 | 55757 | UGGT2 | UDP-glucose glycoprotein glucosyltransferase 2 |
| ENSG00000109944 | 79864 | JHY | Junctional cadherin complex regulator |
| ENSG00000127252 | 57110 | PLAAT1 | Phospholipase A and acyltransferase 1 |
| ENSG00000140470 | 170691 | ADAMTS17 | ADAM metallopeptidase with thrombospondin type 1 motif 17 |
| ENSG00000147255 | 3547 | IGSF1 | Immunoglobulin superfamily member 1 |
| ENSG00000156510 | 80201 | HKDC1 | Hexokinase domain containing 1 |
| ENSG00000162267 | 3699 | ITIH3 | Inter-alpha-trypsin inhibitor heavy chain 3 |
| ENSG00000164904 | 501 | ALDH7A1 | Aldehyde dehydrogenase 7 family member A1 |
| ENSG00000170231 | 2172 | FABP6 | Fatty acid binding protein 6 |
| ENSG00000170315 | 7314 | UBB | Ubiquitin B |
| ENSG00000170949 | 90338 | ZNF160 | Zinc finger protein 160 |
| ENSG00000171094 | 238 | ALK | ALK receptor tyrosine kinase |
| ENSG00000172803 | 254122 | SNX32 | Sorting nexin 32 |
| ENSG00000176595 | 9920 | KBTBD11 | Kelch repeat and BTB domain containing 11 |
| ENSG00000181804 | 285195 | SLC9A9 | Solute carrier family 9 member A9 |
| ENSG00000182175 | 56963 | RGMA | Repulsive guidance molecule BMP co-receptor A |
| ENSG00000183779 | 80139 | ZNF703 | Zinc finger protein 703 |
| ENSG00000196420 | 6276 | S100A5 | S100 calcium binding protein A5 |
| ENSG00000204851 | 57469 | PNMA8B | PNMA family member 8B |
| ENSG00000207383 | - | Y_RNA | Y RNA |
| ENSG00000213071 | 80350 | LPAL2 | Lipoprotein(a) like 2, pseudogene |
| ENSG00000215018 | 340267 | COL28A1 | Collagen type XXVIII alpha 1 chain |
| ENSG00000223652 | - | AC106786.1 | Novel transcript |
| ENSG00000224536 | - | AC096677.1 | Novel transcript |
| ENSG00000225205 | - | AC078883.1 | Novel transcript |
| ENSG00000226210 | 100288778 | WASH8P | WAS protein family homolog 8, pseudogene |
| ENSG00000226476 | 105378763 | LINC01748 | Long intergenic non-protein coding RNA 1748 |
| ENSG00000234380 | 100506385 | LINC01426 | Long intergenic non-protein coding RNA 1426 |
| ENSG00000241431 | - | RPL37P6 | Ribosomal protein L37 pseudogene 6 |
| ENSG00000248508 | 100131089 | SRP14-AS1 | SRP14 antisense RNA1 (head to head) |

**Table S4.** KEGG pathway enrichment of expression signature gene list in LUSC by using *clusterProfiler* R package.

| Pathway ID | Description | P value | Gene ID | Count |
|---|---|---|---|---|
| hsa00330 | Arginine and proline metabolism | 0,002 | NOS1/ALDH7A1 | 2 |
| hsa00010 | Glycolysis / Gluconeogenesis | 0,004 | HKDC1/ALDH7A1 | 2 |
| hsa04066 | HIF-1 signaling pathway | 0,011 | EDN1/HKDC1 | 2 |
| hsa04926 | Relaxin signaling pathway | 0,015 | EDN1/NOS1 | 2 |
| hsa00220 | Arginine biosynthesis | 0,032 | NOS1 | 1 |
| hsa00340 | Histidine metabolism | 0,032 | ALDH7A1 | 1 |
| hsa00053 | Ascorbate and aldarate metabolism | 0,044 | ALDH7A1 | 1 |
| hsa00410 | beta-Alanine metabolism | 0,044 | ALDH7A1 | 1 |
| hsa00052 | Galactose metabolism | 0,045 | HKDC1 | 1 |
| hsa00051 | Fructose and mannose metabolism | 0,048 | HKDC1 | 1 |
| hsa05131 | Shigellosis | 0,050 | HKDC1/UBB | 2 |
| hsa00500 | Starch and sucrose metabolism | 0,052 | HKDC1 | 1 |
| hsa00620 | Pyruvate metabolism | 0,056 | ALDH7A1 | 1 |
| hsa00260 | Glycine, serine and threonine metabolism | 0,058 | ALDH7A1 | 1 |
| hsa00380 | Tryptophan metabolism | 0,061 | ALDH7A1 | 1 |
| hsa00071 | Fatty acid degradation | 0,063 | ALDH7A1 | 1 |
| hsa04930 | Type II diabetes mellitus | 0,066 | HKDC1 | 1 |
| hsa04973 | Carbohydrate digestion and absorption | 0,068 | HKDC1 | 1 |
| hsa00280 | Valine, leucine and isoleucine degradation | 0,069 | ALDH7A1 | 1 |
| hsa00520 | Amino sugar and nucleotide sugar metabolism | 0,069 | HKDC1 | 1 |
| hsa04730 | Long-term depression | 0,086 | NOS1 | 1 |
| hsa00561 | Glycerolipid metabolism | 0,087 | ALDH7A1 | 1 |
| hsa00310 | Lysine degradation | 0,090 | ALDH7A1 | 1 |
| hsa04137 | Mitophagy - animal | 0,097 | UBB | 1 |
| hsa04924 | Renin secretion | 0,098 | EDN1 | 1 |
| hsa05230 | Central carbon metabolism in cancer | 0,099 | HKDC1 | 1 |
| hsa05223 | Non-small cell lung cancer | 0,102 | ALK | 1 |
| hsa03320 | PPAR signaling pathway | 0,110 | FABP6 | 1 |
| hsa05235 | PD-L1 expression and PD-1 checkpoint pathway in cancer | 0,125 | ALK | 1 |
| hsa05410 | Hypertrophic cardiomyopathy | 0,126 | EDN1 | 1 |
| hsa04970 | Salivary secretion | 0,130 | NOS1 | 1 |
| hsa04350 | TGF-beta signaling pathway | 0,131 | RGMA | 1 |
| hsa04713 | Circadian entrainment | 0,135 | NOS1 | 1 |
| hsa04933 | AGE-RAGE signaling pathway in diabetic complications | 0,139 | EDN1 | 1 |
| hsa04916 | Melanogenesis | 0,140 | EDN1 | 1 |
| hsa04974 | Protein digestion and absorption | 0,143 | COL28A1 | 1 |
| hsa04668 | TNF signaling pathway | 0,154 | EDN1 | 1 |
| hsa05022 | Pathways of neurodegeneration - multiple diseases | 0,154 | NOS1/UBB | 2 |
| hsa01200 | Carbon metabolism | 0,162 | HKDC1 | 1 |

**Table S5.** SomInaClust result of SNV data in tumor samples of LUAD low-risk group.

| Gene | Number of Mutations | Q-value | OG Score | TSG Score | Classification | CGC[1] |
|---|---|---|---|---|---|---|
| KRAS | 74 | 1.24e-128 | 97.3 | 0 | OG | Dom |
| TP53 | 148 | 1.16e-68 | 82 | 34.5 | TSG | Rec |
| EGFR | 40 | 7.46e-39 | 78.8 | 0 | OG | Dom |
| BRAF | 26 | 2.87e-27 | 69.6 | 12.5 | OG | Dom |
| STK11 | 43 | 1.26e-26 | 12.5 | 69 | TSG | Rec |
| MGA | 36 | 4.25e-16 | 0 | 60.6 | TSG | NA |
| NF1 | 45 | 4.25e-16 | 0 | 52.4 | TSG | Rec |
| RB1 | 20 | 4.81e-12 | 0 | 70 | TSG | Rec |
| PIK3CA | 18 | 8.18e-11 | 58.8 | 0 | OG | Dom |
| ATM | 30 | 1.05e-10 | 20 | 53.6 | TSG | Rec |
| RBM10 | 17 | 1.05e-10 | 0 | 75 | TSG | NA |
| SETD2 | 22 | 3.49e-10 | 16.7 | 61.9 | TSG | Rec |
| ARID1A | 22 | 7.28e-10 | 0 | 59.1 | TSG | Rec |
| CTNNB1 | 12 | 3.13e-08 | 72.7 | 0 | OG | Dom |
| CMTR2 | 18 | 5.35e-06 | 0 | 56.2 | TSG | NA |
| SF3B1 | 12 | 1.53e-05 | 55.6 | 0 | OG | Dom |
| CSMD3 | 219 | 7.36e-05 | 0 | 16.3 | NA | NA |
| ATF7IP | 13 | 8.25e-05 | 0 | 63.6 | TSG | NA |
| KEAP1 | 47 | 0.000259 | 25 | 22.5 | TSG | NA |
| HMCN1 | 65 | 0.000464 | 0 | 28.2 | TSG | NA |
| EPHA5 | 47 | 0.00048 | 33.3 | 26.5 | TSG | NA |
| ARID2 | 11 | 0.00108 | 0 | 60 | TSG | Rec |
| TTK | 13 | 0.00411 | 0 | 50 | TSG | NA |
| SMAD4 | 14 | 0.0101 | 50 | 42.9 | TSG | Rec |
| KDM5C | 10 | 0.0121 | 0 | 55.6 | TSG | Rec |
| SMARCA4 | 26 | 0.0158 | 0 | 40 | TSG | Rec |
| APC | 10 | 0.0209 | 0 | 50 | TSG | Rec |
| NFE2L2 | 8 | 0.0211 | 57.1 | 0 | OG | Dom |
| RIT1 | 5 | 0.0211 | 75 | 0 | OG | NA |
| DDX10 | 8 | 0.0302 | 0 | 66.7 | TSG | Dom |
| LTN1 | 14 | 0.0322 | 0 | 45.5 | TSG | NA |
| CDH10 | 66 | 0.0381 | 0 | 20.9 | TSG | NA |
| SPTA1 | 114 | 0.0381 | 0 | 18.9 | NA | NA |
| LRP1B | 174 | 0.0384 | 0 | 16 | NA | Rec |
| COL11A1 | 79 | 0.0394 | 4.5 | 18.5 | NA | NA |
| MAP3K12 | 5 | 0.0394 | 0 | 100 | TSG | NA |
| USH2A | 161 | 0.0394 | 0 | 15.8 | NA | NA |
| AKAP6 | 29 | 0.0468 | 0 | 31.6 | TSG | NA |
| RASA1 | 7 | 0.0483 | 0 | 57.1 | TSG | NA |

[1] COSMIC Cancer Gene Census (Dom: Dominant, Rec: Recessive)

**Table S6.** SomInaClust result of SNV data in tumor samples of LUAD high-risk group.

| Gene | Number of Mutations | Q-value | OG Score | TSG Score | Classification | CGC[1] |
|---|---|---|---|---|---|---|
| KRAS | 64 | 3.76e-113 | 98.4 | 0 | OG | Dom |
| TP53 | 96 | 8.46e-50 | 84.6 | 41.5 | TSG | Rec |
| STK11 | 37 | 1.45e-27 | 50 | 75 | TSG | Rec |
| EGFR | 31 | 6.01e-26 | 78.3 | 0 | OG | Dom |
| BRAF | 16 | 6.42e-16 | 66.7 | 10 | OG | Dom |
| RBM10 | 21 | 1.59e-15 | 0 | 80 | TSG | NA |
| PIK3CA | 8 | 6.8e-09 | 87.5 | 0 | OG | Dom |
| SETD2 | 20 | 1.24e-07 | 16.7 | 57.9 | TSG | Rec |
| ARID2 | 18 | 6.33e-07 | 0 | 58.8 | TSG | Rec |
| NF1 | 13 | 7.39e-06 | 0 | 66.7 | TSG | Rec |
| RB1 | 12 | 7.39e-06 | 0 | 66.7 | TSG | Rec |
| MGA | 16 | 7.5e-05 | 0 | 53.3 | TSG | NA |
| KEAP1 | 41 | 8.07e-05 | 27.8 | 25.7 | TSG | NA |
| CSMD3 | 93 | 0.00504 | 0 | 21.1 | TSG | NA |
| SMARCA4 | 21 | 0.00514 | 0 | 50 | TSG | Rec |
| CTNNB1 | 8 | 0.00743 | 66.7 | 16.7 | OG | Dom |
| KDM5C | 6 | 0.0159 | 0 | 80 | TSG | Rec |
| IDH1 | 3 | 0.0277 | 66.7 | 0 | OG | Dom |
| ATM | 17 | 0.0296 | 25 | 37.5 | TSG | Rec |

[1] COSMIC Cancer Gene Census (Dom: Dominant, Rec: Recessive)

**Table S7.** SomInaClust result of SNV data in tumor samples of LUSC low-risk group.

| Gene | Number of Mutations | Q-value | OG Score | TSG Score | Classification | CGC[1] |
|---|---|---|---|---|---|---|
| TP53 | 185 | 4.35e-90 | 85.6 | 30.9 | TSG | Rec |
| KMT2D | 63 | 7.77e-41 | 0 | 72.4 | TSG | Rec |
| NFE2L2 | 29 | 5.5e-25 | 83.3 | 4.2 | OG | Dom |
| PIK3CA | 20 | 3.19e-20 | 78.9 | 0 | OG | Dom |
| CDKN2A | 34 | 2.81e-19 | 33.3 | 66.7 | TSG | Rec |
| PTEN | 26 | 1.66e-16 | 62.5 | 65.4 | TSG | Dom |
| RB1 | 19 | 2.3e-14 | 0 | 78.9 | TSG | Rec |
| FAT1 | 38 | 2.73e-12 | 0 | 53.1 | TSG | NA |
| ARID1A | 19 | 7.73e-08 | 0 | 57.9 | TSG | Rec |
| NF1 | 25 | 8.65e-08 | 0 | 50 | TSG | Rec |
| RASA1 | 11 | 6.17e-07 | 50 | 80 | TSG | NA |
| CUL3 | 15 | 3.1e-05 | 25 | 57.1 | TSG | NA |
| KDM6A | 11 | 3.1e-05 | 0 | 70 | TSG | Rec |
| NRAS | 4 | 7.98e-05 | 100 | 0 | OG | Dom |
| KRT5 | 14 | 0.000151 | 0 | 100 | TSG | NA |
| ZNF750 | 8 | 0.000807 | 0 | 83.3 | TSG | NA |
| EP300 | 12 | 0.000867 | 42.9 | 9.1 | OG | Rec |
| FGFR3 | 4 | 0.00139 | 100 | 0 | OG | Dom |
| TAOK1 | 5 | 0.00301 | 0 | 100 | TSG | NA |
| CSMD3 | 140 | 0.00301 | 0 | 17.2 | NA | NA |
| NSD1 | 16 | 0.00631 | 0 | 46.2 | TSG | Dom |
| TTN | 394 | 0.00636 | 1.8 | 12.2 | NA | NA |
| HRAS | 3 | 0.00948 | 100 | 0 | OG | Dom |
| SI | 41 | 0.0108 | 0 | 28.6 | TSG | NA |
| PDS5B | 6 | 0.011 | 0 | 80 | TSG | NA |
| KRAS | 4 | 0.0116 | 75 | 0 | OG | Dom |
| KEAP1 | 25 | 0.0399 | 28.6 | 4.8 | OG | NA |
| API5 | 5 | 0.0469 | 0 | 100 | TSG | NA |
| HNRNPUL1 | 5 | 0.0469 | 0 | 100 | TSG | NA |
| SLC16A1 | 3 | 0.0469 | 0 | 100 | TSG | NA |
| FBXW7 | 12 | 0.0487 | 33.3 | 41.7 | TSG | Rec |

[1] COSMIC Cancer Gene Census (Dom: Dominant, Rec: Recessive)

**Table S8.** SomInaClust result of SNV data in tumor samples of LUSC high-risk group.

| Gene | Number of Mutations | Q-value | OG Score | TSG Score | Classification | CGC[1] |
|---|---|---|---|---|---|---|
| TP53 | 211 | 9.03e-117 | 88.6 | 37.2 | TSG | Rec |
| NFE2L2 | 42 | 4.44e-41 | 86.1 | 0 | OG | Dom |
| PIK3CA | 34 | 3.59e-38 | 83.9 | 7.1 | OG | Dom |
| KMT2D | 69 | 1.22e-24 | 0 | 51.6 | TSG | Rec |
| FAT1 | 42 | 7.42e-17 | 0 | 60 | TSG | NA |
| CDKN2A | 40 | 1.63e-14 | 83.3 | 41 | TSG | Rec |
| RB1 | 15 | 4.32e-11 | 0 | 80 | TSG | Rec |
| PTEN | 25 | 9.24e-10 | 50 | 48 | TSG | Dom |
| NOTCH1 | 24 | 3.11e-08 | 0 | 54.5 | TSG | NA |
| ARID1A | 13 | 4.45e-07 | 0 | 69.2 | TSG | Rec |
| RASA1 | 19 | 2.22e-05 | 0 | 50 | TSG | NA |
| NF1 | 29 | 0.000106 | 0 | 37 | TSG | Rec |
| KMT2C | 31 | 0.000145 | 0 | 35.7 | TSG | Rec |
| BRAF | 9 | 0.000274 | 50 | 20 | OG | Dom |
| PIK3R1 | 10 | 0.000496 | 0 | 66.7 | TSG | Rec |
| CSMD3 | 155 | 0.000554 | 0 | 17.7 | NA | NA |
| STK11 | 4 | 0.00322 | NaN | 100 | TSG | Rec |
| HRAS | 4 | 0.0113 | 100 | 0 | OG | Dom |
| KEAP1 | 28 | 0.0444 | 28.6 | 12.5 | OG | NA |

[1] COSMIC Cancer Gene Census (Dom: Dominant, Rec: Recessive)
